# Flexible foraging decisions by configuration of orthogonal integrator dynamics

**DOI:** 10.1101/2025.11.14.688540

**Authors:** Lyle Kingsbury, Grace Zhang, Juan Ignacio Sanguinetti-Scheck, Naoshige Uchida

**Affiliations:** Dept. of Molecular and Cellular Biology, Harvard University, Cambridge, MA, USA; Center for Brain Science, Harvard University, Cambridge, MA, USA; Dept. of Psychology, University of Pennsylvania, Philadelphia, PA

## Abstract

The capacity to adapt behavior and cognition across contexts is fundamental to intelligence. Context-dependent decision making is an exemplar of flexible cognition in animals, yet its biological basis remains poorly understood. At the neural level, a critical mechanistic question is whether context largely alters the inputs to a common decision process or reconfigures decision activity itself to create context-specific functional modes. To study this problem, we developed a patch foraging task in which mice forage in two environment contexts defined by distinct reward dynamics. Mice made patch leaving decisions across environments using different context-specific parameterizations of a drift-diffusion integrator process. Using high-density acute and chronic neural recordings, we find that foraging environments recruit separate activity subspaces which organize orthogonal coding of context-specific decision variables. Unlike the context-invariant decision coding used for perceptual choice problems, foraging decision variables were encoded in orthogonal population vectors within largely separate neural subpopulations – a subspace configuration mechanism which can support modular learning, independent readout, and flexible toggling of decision strategies across environments. In the dorsal frontal cortex, orthogonal integrators emerge with experience, are preserved across three distinct task settings, and are required for rapid and volitional strategy switching. Our findings establish orthogonal integrator dynamics as a general solution for context-dependent decision making and open a new avenue to investigate the cell- and circuit-level basis of flexible cognition in the mammalian brain.

## Introduction

The capacity to adapt cognition and behavior across different contexts is a core feature of intelligence and is crucial for surviving in dynamic natural environments. A major goal of neuroscience is to understand how cognitive processes such as context-dependent decision making arise from underlying biological mechanisms. In mammals, the frontal cortex is critical for flexible cognition, which is thought to involve context-dependent configuration of neural networks for information processing and decision computations.^1–5^ Theoretical studies give insight into how functionally distinct network modes may be organized by neuron physiology and microcircuit architectures,^6–10^ but despite these advances, experimental progress in grounding models of context-dependent processes at the cellular and circuit level has been remarkably limited.

A fundamental question for mechanistic understanding of context-dependent decision making is whether context largely shapes upstream inputs to a common (i.e. context-invariant) decision process or directly reconfigures the decision process itself (**Fig. 1**). To date, investigations of this problem in animal models have mostly used classical, rule-based perceptual choice tasks.^11–15^ This research program has clarified a diversity of circuit mechanisms that operate upstream of decision formation and can incorporate context contingencies through attentional shifting,^13,14,16–18^ sensorimotor re-mapping,^12,19^ and altering recurrent input dynamics.^11,15,20^

**Figure 1:**
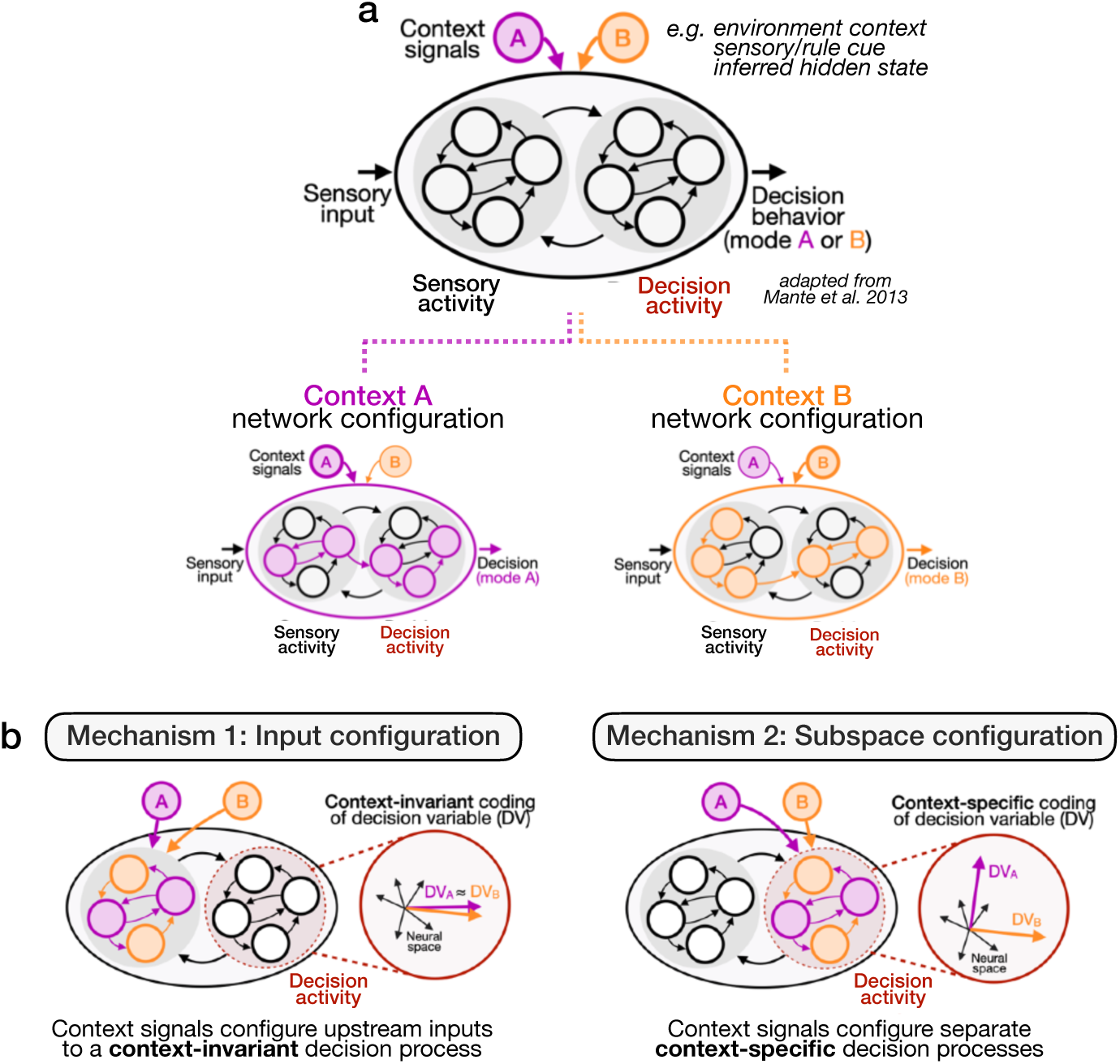
Conceptual models and hypothetical mechanisms for context-dependent decision making. a) Top, conceptual model of neural network control of context-dependent decision making (adapted from Mante et al. 2013) - context (“A” or “B”) signals configure different operating modes for a decision process by altering neural dynamics for sensory processing and/or decision formation. Bottom, cartoon of context A and context B neural network configurations which process common sensory inputs differently to produce contingent decision behavior. b) Illustration of hypothetical neural network mechanisms for context-dependent decision making. **Input configuration** mechanisms operate on processing steps upstream of decision formation, preserving context-invariant decision activity, as observed for rule-based perceptual decisions. Alternatively, **Subspace configuration** mechanisms operate on decision activity itself, parsing decision variables into context-specific coding vectors contained in separate subspaces.

However, while it has led to foundational insights, this focus on perceptual choice tasks may also limit the explanatory scope of mechanistic models and be blind to other possibilities.^21,22^ For example, other studies^7,10,23–28^ suggest that context-dependency may depend on orthogonal coding of decision activity in separate subspaces. In theory, this neural organization can support modular learning, non-interfering readout of decision computations, and rapid toggling between decision strategies – yet empirical evidence for it is scant. Additionally, it is not known whether mechanisms for perceptual choice problems generalize outside that domain, nor how neural solutions for flexible cognition operate in naturalistic environments that require rapid strategy shifting.

To address these questions, we studied context-dependent decision making in the ecological setting of foraging – a ubiquitous survival challenge faced across the animal kingdom.^21,22,29,30^ In the natural world, animals routinely make decisions about where to forage and when to leave depleting resource locations (patches) in search of alternatives (the *patch leaving problem*),^31,32^ adjusting strategies across changing environments. Because of its intuitiveness and ecological grounding, foraging has gained recent attention in neuroscience, and studies have begun to uncover the neural basis of stay-or-leave foraging decisions.^33,34^ Evidence suggests that patch leaving decisions may be solved through an integrator process akin to bounded drift-diffusion, linking natural cognition with classical decision models – but it is not known whether or how these computations are reconfigured at the neural level to support flexible decision strategies.^35–39^

We developed an experimental paradigm in which mice flexibly toggle foraging strategies across environment contexts using different configurations of an integrator decision process. Using an equivalent virtual task for wide neural sampling, we find that environment contexts recruit distinct, distributed neural states with separated activity subspaces. In contrast to the context-invariant decision activity observed for rule-based perceptual choices, integrator dynamics for foraging decisions are parsed into context-specific, orthogonal coding vectors. In the dorsal frontal cortex (dFC), this coding structure is shaped over days and preserved across diverse task settings, even when mice make repeated, volitional context switches. Inactivation experiments demonstrate causal necessity of dFC for rapid reconfiguration of decision modes. Together, our results reveal a novel neural solution for context-dependent decision making in orthogonal integrator dynamics and open a new avenue to study the biological mechanisms of cognition in the mammalian brain.

## Main

### Context-dependent decision making through configuration of neural network modes

Context-dependent decision making is an exemplar of flexible cognition in animals. Its defining structural feature is contingency: common sensory inputs must be processed differently in different contexts, leading to a behavioral output (a decision) that is contingent on context identity (**Fig. 1a**, top).^8,11^ Theoretically, this can be achieved by a neural network with context-specific configurations (**Fig. 1a**, bottom), yet how this occurs in the brain is not well understood. At the neural level, context could shape the inputs to a general context-invariant decision process (**Fig. 1b**, *input configuration*), as has been observed for rule-based perceptual choice. Context signals could also reconfigure the decision process itself, creating context-specific neural modes for decision computations (**Fig. 1b**, *subspace configuration*). To study the neural basis of context-dependent decision making in an ecological setting, we developed a foraging task for mice that requires different decision strategies across environments, and we used high-density electrophysiology to investigate the structure and function of the underlying neural dynamics.

### A patch foraging task for context-dependent decision making in mice

In patch foraging,^31,32^ animals navigate an environment for resource patches that deplete with consumption and must decide when to leave a current patch in search of alternatives (**Fig. 2a**). We developed a task in which mice forage in two environments containing patches with distinct reward dynamics, encouraging context-specific decision strategies. Mice navigate an arena with two chambers (environment contexts) for water rewards from multiple ports (patches, **Fig. 2b,c**). Mice can nose-poke to initiate a trial and immediately (t = 0s) receive a 2µL water drop, after which they can stay for more rewards. Patches in the Stochastic context (**Fig. 2b**, top) yield decaying-probability rewards each second (**Extended Data Fig. 1a**), whereas patches in the Deterministic context (**Fig. 2b**, bottom) always yield three fixed rewards (at t = 0, 1, 2s). 15% of Stochastic trials were programmed with this return (“Stc:012” trials with rewards at t = 0, 1, 2s) to provide reward-matched trial types for direct comparison of behavior across contexts.

**Figure 2:**
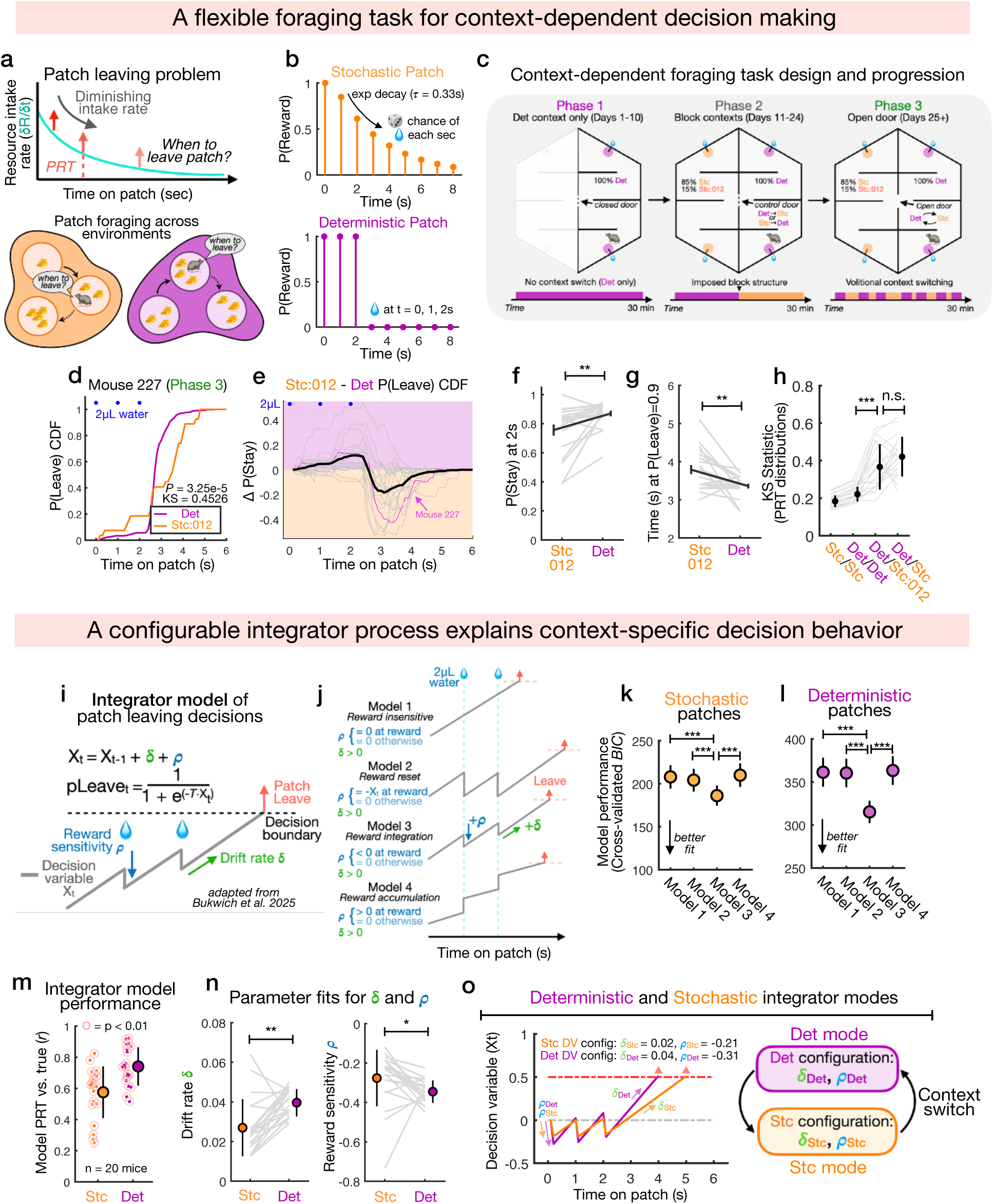
Mice flexibly use context-specific foraging strategies across different environments. a) Illustration of the classical patch leaving problem from behavioral ecology (when to leave a patch with depleting resources). Animals make patch leaving decisions in different environment contexts that contain distinct resource distributions or dynamics. b) Reward returns (2µL water drops with probability each second) for Stochastic (orange) and Deterministic (purple) patches (see also **Extended Data** Fig. 1a). c) Progression through flexible foraging task, from Deterministic environment (Phase 1, 7-10 days) to block structured environments (Phase 2, 14 days, alternating the starting context) and unconstrained volitional transitions with the separating door left open (Phase 3, 4+ days). d) CDF of leaving probabilities P(Leave)_t_ for Deterministic and Stc:012 patches from example Mouse 227 (KS test, *P* = 3.25e-5). e) Difference in stay probability P(Stay) on Deterministic vs. Stc:012 patches for each mouse (gray lines), mouse 227 (pink line), and average across mice (black line, *n* = 20 mice). f) Comparison of P(Stay) at t = 2 seconds for Deterministic vs. Stc:012 patches (mean ± SEM, SR test, *P* = 0.0013, *n* = 20 mice). g) Comparison of time (seconds) on patch when P(Leave) = 0.9 for Deterministic vs. Stc:012 patches (mean ± SEM, SR test, *P* = 0.0038, *n* = 20 mice). h) Kolmogorov-Smirnov (KS) statistics comparing PRT distributions across different trial types (mean ± s.d., SR tests, *P* = 2.19e-4, Det/Det vs. Det/Stc:012, *P* = 0.0522, Det/Stc:012 vs. Det/ Stc, *n* = 20 mice). i) Schematic of drift-diffusion integrator model for patch-leaving decisions (based on previous work by Bukwich et al. 2025). Decision variable (DV) X_t_ is incremented each time step by drift rate δ and perturbed by ρ upon reward receipt (ρ = 0 otherwise), accumulating to a boundary to trigger a leaving decision. pLeave_t_ is generated by a logistic function transform of X_t_ with temperature *T* (see **Methods**). j) Parameter configurations (δ, ρ) for the integrator model (Fig. 2i) implement different algorithms for computing patch leaving decisions (Models 1-4). k) Performance (cross-validated Bayesian information criterion, BIC) of integrator Models 1-4 (Fig. 2j) fit to mouse PRT behavior and tested on held-out data for Stochastic patch trials (mean ± SEM, SR test between models 1v3, 2v3, 3v4, *P* = 0.857e-5 for each, *n* = 20 mice). l) Model performance as in (Fig. 2k) for Deterministic patch trials (mean ± SEM, SR test between models 1v3, 2v3, 3v4; *P* = 0.857e-5 for each, *n* = 20 mice). m) Performance of “reward integration” Model 3 (correlation *r* between predicted and true PRTs, mean ± s.d.) for Stochastic and Deterministic patch trials. Each mouse is one dot; red circles indicate independent statistical significance (*P* < 0.01) of regression on predicted vs. true PRTs for that mouse (see also **Extended Data** Fig. 2k**,l**). n) Left, comparison of fit drift rate parameter δ to Model 3 for Stochastic vs. Deterministic trials (mean ± s.d., SR test, *P* = 0.005, *n* = 20 mice). Right, comparison of reward sensitivity parameter ρ for Stochastic vs. Deterministic trials (mean ± s.d., SR test, *P* = 0.027, *n* = 20 mice). o) Left, rollout of Model 3 DV for average (across mice) integrator parameters fit to Stochastic or Deterministic patches (Fig. 2n). Right, distinct parameter configurations produce different modes for patch leaving decisions that can explain context-specific patterns of PRT distributions. Standard deviation (s.d.); Standard error of the mean (SEM); Kolmogorov-Smirnov test (KS test); Wilcoxon signed-rank test (SR test); \*\*\**P* < 0.001; \*\**P* < 0.01; \**P* < 0.05; n.s. *P* > 0.05.

Mice first foraged in the Deterministic context to learn a decision strategy for Deterministic rewards (**Fig. 2c**, left). Mice foraged in the first session and rapidly learned to stay at patches for additional rewards (**Extended Data Fig. 1b-e**). Analysis of patch residence time (PRT, how long mice stay at a given patch trial) distributions revealed calibration of the decision process over days, with PRTs clustered after the final reward receipt (**Extended Data Fig. 1f-g**). In Phase 2, mice explored both Deterministic and Stochastic contexts once per session in blocks (**Fig. 2c**, middle) to learn context-specific strategies, and in Phase 3, the door was left open so they could switch environments volitionally (**Fig. 2c**, right). When given the choice, mice preferred and collected more rewards at Deterministic patches (**Extended Data Figs. 1h,i**), possibly reflecting risk attitudes or small differences in reward expectation. Nonetheless, mice explored both environments and switched frequently (about 1-4 times/min, **Extended Data Fig. 1j,k**). On Stochastic patches, mice scaled their PRTs with reward frequency, indicating value-sensitive decisions that reflect the decaying reward rate structure (**Extended Data Fig. 2a-b**).

### Mice exhibit context-specific decision strategies across foraging environments

To test whether mice use context-dependent decision strategies, we compared PRT distributions for Deterministic and reward-matched Stc:012 trials by plotting the cumulative density function (CDF) of P(Leave) over time. Despite identical reward returns, mice showed distinct patterns of leaving behavior (**Fig. 2d**). For example, while Mouse 227 waited to collect all Deterministic rewards and left shortly after, it did not exhibit this step-like pattern on Stc:012 patches, more often leaving before the final reward or staying seconds after. This pattern was consistent across mice (**Fig. 2e**), and mice were generally more likely to collect all rewards on Deterministic trials and faster to leave after the final reward (**Fig. 2f-g**). To gain insight into the decision strategies, we simulated patch leaving decisions by applying a reward-maximizing policy based on the Marginal Value Theorem (MVT, **Extended Data Fig. 2c-e, Methods**). This replicated the pattern of decision behavior under the assumption that Deterministic leaving decisions are driven by expected reward rates (reflecting knowledge of the reward structure), while Stochastic patch decisions are driven by local experienced rewards.

To quantify this decision flexibility, we measured the difference between Deterministic and Stc:012 PRT distributions using the Kolmogorov-Smirnov (KS) statistic (**Extended Data Fig. 2f**). These diverged significantly more than PRTs across subsampled Deterministic trials, despite identical rewards, confirming that the behavioral difference is driven by context (**Fig. 2h**). Decision flexibility increased over time, indicating calibration of decision strategies that are maintained through Phase 3 (**Extended Data Fig. 2g**). While this shows that mice use distinct strategies, they could be reliant on recent trial outcomes to update their decision process. To test this, we analyzed trial sequences in which mice performed a Stc:012 followed by a Deterministic trial and observed similar PRT distributions (**Extended Data Fig. 2h-j).** Furthermore, mice trained on a task variant with hidden rather than spatial contexts could not adapt their decisions, suggesting that they use the environment as a cue to switch decision strategies rather than online experience (**Extended Data Fig. 3**). Together, these results demonstrate that mice calibrate context-specific decision strategies to reward dynamics in different environments and rapidly toggle between them.

### Configurations of a common integrator process explain decision flexibility

We have previously shown that patch leaving decisions in mice are well described by an integrator process that may be instantiated by neural dynamics oppositely weighing time and reward gains (**Fig. 2i**).^35,38^ We reasoned that this model may similarly explain decisions in the present task and may additionally explain context-dependent decision flexibility. To test this, we modeled leaving decisions as a drift-diffusion decision variable (DV) which increments each time step (drift rate ***δ***) and is perturbed on reward receipt (reward sensitivity **ρ**), then used to generate a time-varying leaving probability P(Leave). In this integrator model, as in models of perceptual decisions,^36,37^ ***δ*** and **ρ** govern the DV dynamics with different parameterizations producing distinct algorithms for computing leaving decisions (**Fig. 2j**). For example, while **ρ** = 0 yields a reward-insensitive integrator tracking time (Model 1),^31^ **ρ** < 0 integrates time investment vs. rewards (Model 3), and **ρ** > 0 (with small ***δ***) yields a counting algorithm that can track reward quantity (Model 4).^35^

In principle, mice could use different algorithms to compute leaving decisions in Stochastic vs. Deterministic contexts. To test which algorithm best explained their behavior, we fit different formulations of the integrator model (Models 1-4, **Fig. 2j**) to empirical PRTs from mice and measured predictions on held-out trials. Consistent with our previous study,^38^ we found that the “reward integration” algorithm (Model 3) significantly outperformed others (**Fig. 2k,l**). Despite clearly distinct behavior (**Fig. 2d-h**), this algorithm captured PRTs for both Deterministic and Stochastic patches (**Fig. 2m**), raising the possibility that context-specificity could be driven by different configurations (i.e. parameters ***δ*** and **ρ**) of this common decision algorithm.

To test this, we compared ***δ*** and **ρ** for models fit to Stochastic and Deterministic patches and found clear differences in parameterization: while Deterministic patches had significantly larger drift rates ***δ*** (**Fig. 2n**, left), they also had larger negative values for reward sensitivity **ρ** (**Fig. 2n**, right). To understand how these configurations could produce the observed behavior, we plotted DVs following a Deterministic return (rewards at t = 0, 1, 2s) using the average parameters across mice (**Fig. 2o**, left). This produced distinct DV rollouts which translate into different P(Leave) time-courses on Deterministic vs. Stochastic patches, effectively calibrating leaving decisions for distinct reward dynamics. Indeed, sampling PRTs from model-fit DVs for individual mice qualitatively replicated their behavioral patterns (**Extended Data Fig. 2k,l**). These results raise the possibility that flexible patch leaving decisions may be driven by different decision modes defined by context-specific configurations of a common integrator process (**Fig. 2o**, right).

### An equivalent head-fixed virtual task for wide sampling of neural activity

To test this idea, we next investigated the organization of neural dynamics underlying context-dependent foraging decisions. While the free-moving (FM) setting was critical for understanding natural behavior,^21,22^ constraint with head-fixation enables wide sampling of neural activity in a more controlled setting with cleaner interpretability (**Fig. 3a**).^40^ We therefore developed a structural equivalent virtual-reality^38^ (VR) version of our task (**Fig. 3b**), with the goal of identifying neural mechanisms that we could study across a range of task designs with different constraints. In VR, mice run on a treadmill to navigate a virtual corridor and stop at patches (visual/odor-cued landmarks) to collect rewards. Mice encountered Stochastic and Deterministic patches in distinct environments. Deterministic patches yielded three rewards at fixed times (at t = 0, 2, 4s, **Fig. 3c**), and 10% of Stochastic trials had this same return (“Stc:024” trials); otherwise, Stochastic patches gave decaying-probability rewards each second (**Fig. 3c**, **Extended Data Fig. 4a**). Mice followed a similar two-phase task progression as in the FM task (**Fig. 3d**), and in general, they performed the foraging task similarly as in our previous study (**Extended Data Fig. 4b-d**).^38^

**Figure 3.**
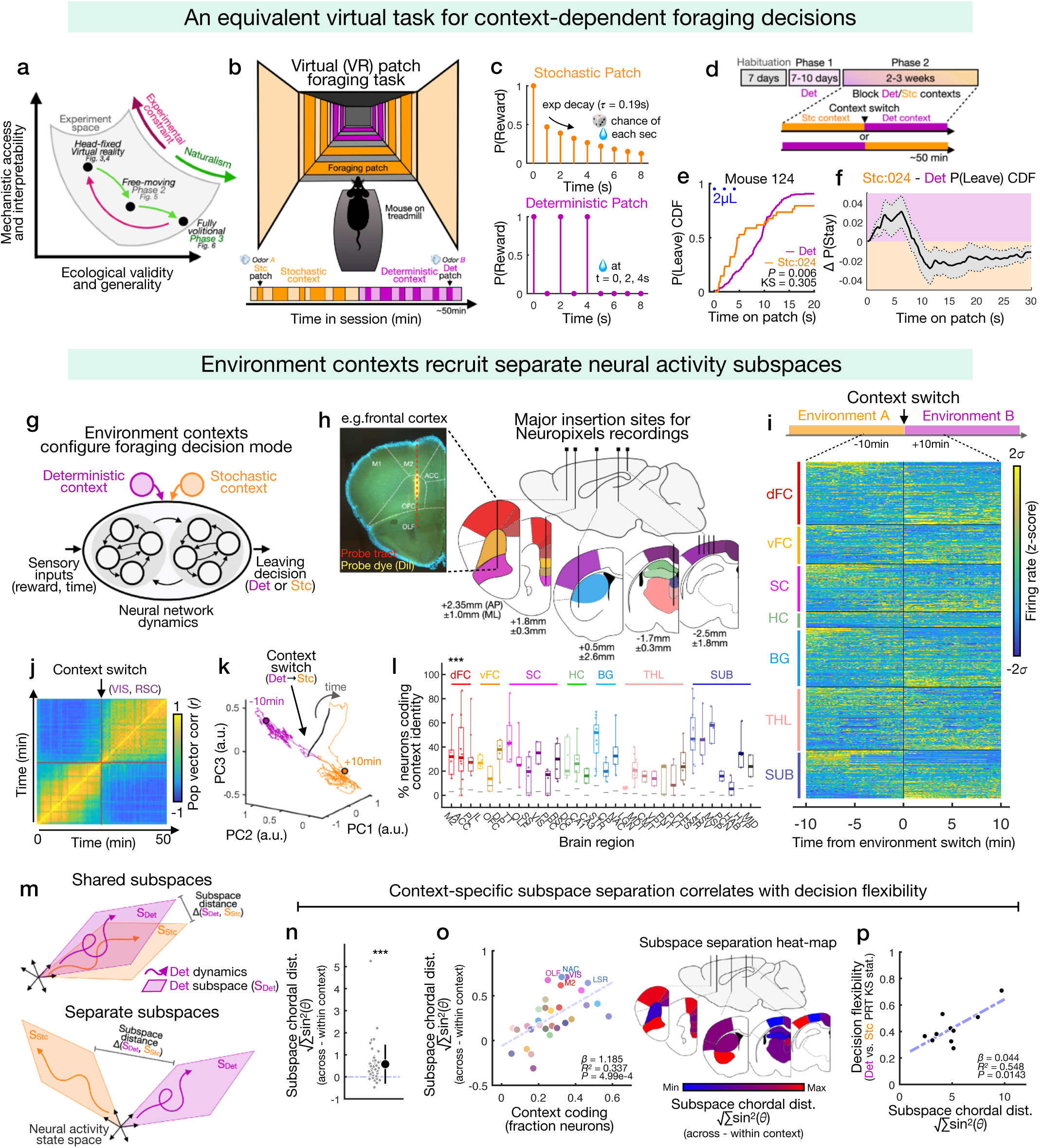
Foraging environments in an equivalent virtual task recruit distinct neural activity states with context-specific dynamics. a) Cartoon illustrating the fundamental tradeoff in experiment design between constraint (such as head-fixation in virtual-reality), which increases mechanistic access and interpretability, and naturalism, which increases ecological validity and generality. We used complementary task designs to clarify neural dynamics mechanisms for flexible decision making (Fig. 3**,4**) and study their operation under settings of increasing ecological complexity (Fig. 5**,6**). b) Structural equivalent virtual-reality (VR) task in which mice forage for water rewards and encounter Stochastic (Stc) and Deterministic (Det) patches in different environment contexts indicated by distinct visuals and odors. c) Reward returns for Deterministic and Stochastic patches with similar structure as in the free-moving (FM) task (Fig. 2b, see also **Extended Data** Fig. 4a). d) Timeline for VR task. Mice are habituated to head fixation and running wheel, first exposed to the Deterministic environment (Phase 1, 7-10 days), and then forage in block-structured environments (Phase 2, alternating the starting context) with one context transition per session. e) CDF of P(Leave)_t_ for Deterministic and Stc:024 patches from example Mouse 124 (KS test, *P* = 0.002). f) Average difference in P(Stay) on Deterministic vs. Stc:024 patches (mean ± SEM, *n* = 36 mice), showing similar context-dependent behavior as in the FM task (Fig. 2e). g) Cartoon illustrating configuration of neural network modes by context signals indicating Deterministic/Stochastic environment for context-dependent foraging decisions (see also **Extended Data** Fig. 6 for comparison of related task designs in previous studies). h) Example histological section showing location of a Neuropixels probe insertion through frontal cortex (dotted red line, probe tract, yellow, probe dye; see also **Extended Data** Fig. 7) and schematic showing major insertion locations with coordinates for acute recording experiments. i) Heat map showing activity of all recorded neurons (*n* = 7804, rows) over a 20-minute window centered around the context transition (change of virtual environment, t = 0, *σ* = s.d. of firing rate). Neurons were recorded from 32 brain regions (see **Extended Data** Fig. 8a and **Supplementary Table 1**) and sorted into 7 “macro-region” categories: dFC (dorsal Frontal cortex), vFC (ventral Frontal cortex), SC (sensory cortex), HC (hippocampal), BG (basal ganglia), THL (thalamic), and SUB (subcortical). j) Correlation matrix of population vectors from an example recording session in higher visual/ retrosplenial cortex. Block structure indicates stable within-context activity patterns that are reorganized at the time of context transition, indicating two distinct neural activity states. k) Principal component (PC) projections of activity from session shown in Fig. 3j showing transient deviation at the time of context transition followed by a persistent change in population activity (gray arrow indicates direction of time). l) Single neuron coding of environment/block identity (fraction of significant neurons for each session/region sample) across regions (median IQR and range, one-way anova, *P* = 5.08e-4, *n* = 187 samples, see **Methods f**or description of statistical tests). m) Schematic illustrating possible relationships between neural activity subspaces. Canonical correlation analysis (CCA, see also **Extended Data** Fig. 10) its used to quantify distance between subspaces, which could be largely shared (top) or separated (bottom). n) Normalized (between - within context) subspace distances (the “chordal distance” √∑sin^2^(*ϴ*) of angles *ϴ* between canonical vectors/canonical correlations ) for Deterministic and Stochastic-context subspaces (mean ± s.d., t-test, *P* = 5.15e-5, *n* = 47 sessions). o) Left, correlation across regions between single-neuron coding of environment identity (Fig. 3l) and subspace chordal distance (linear regression, *P* = 4.95e-4, *n* = 32 regions); right, heat-map visualizing regional heterogeneity in subspace distance (scaled min-max in blue-red). p) Correlation across mice between subspace difference (chordal distance) and context-dependent decision behavior measured using the KS statistic of Stochastic vs. Deterministic PRT distributions (linear regression, *P* = 0.0143, *n* = 10 mice, see also **Extended Data** Fig. 10k). Standard deviation (s.d.); Standard error of the mean (SEM); Kolmogorov-Smirnov test (KS test); Wilcoxon signed-rank test (SR test); \*\*\**P* < 0.001; \*\**P* < 0.01; \**P* < 0.05; n.s. *P* > 0.05.

Like in the FM task, P(Leave) CDFs for matched trials revealed clear context-dependency in the decision process (**Fig. 3e**). On average, mice followed a similar pattern of time-varying differences in P(Leave) (**Fig. 3f**), and Deterministic vs. Stc:024 PRT distributions were significantly more divergent than within-context comparisons (**Extended Data Fig. 4e**). To test whether this decision flexibility could similarly be captured by different configurations of an integrator process, we fit models as before and found that time/reward integration (Model 3, **Fig. 2j**) was again the best fit algorithm and exhibited similar differences in ***δ*** (drift rate) and **ρ** (reward sensitivity) across patch types (**Extended Data Fig. 4f-i**). Behavior of male and female mice was not significantly different in either task (**Extended Data Fig. 5**). Together, this shows that mice use similar foraging strategies across FM and VR tasks, demonstrating robust context-dependent decision making and enabling study of the underlying neural activity across experimental settings.

### Environment contexts recruit distinct brain-wide neural activity states

To enable different modes for context-dependent decisions, context signals must configure neural processes that instantiate or feed into decision computations (**Fig. 1a, 3g**).^8,9,11^ In most classical task designs, context is either inferred from experience or presented as a transient sensory cue (**Extended Data Fig. 6**). Instead, our task uses environment context – an external state which is continuously and unambiguously identified. To investigate how population and decision activity are shaped by environment contexts, we performed high-density *in vivo* electrophysiological recordings using Neuropixels^41^ probes targeting a wide range of brain regions (**Fig. 3h, Extended Data Fig. 7**). We sampled the frontal cortex, which is known to encode integrator DVs for patch foraging,^38,39^ as well as the basal ganglia, subcortical and thalamic nuclei, and sensory cortices spanning olfactory, somatosensory, visual and associative domains. We included a total of 7804 units recorded from 47 sessions across 10 mice for analysis (**Extended Data Fig. 8a**).

To gain an initial impression of the neural data and guide closer investigation, we first explored large-scale activity patterns in seven macro-region categories: dorsal frontal cortex (dFC), ventral frontal cortex (vFC), sensory cortex (SC), hippocampal (HC), basal ganglia (BG), thalamic (THL), and subcortical (SUB) areas (**Supplementary Table 1**). Plots of single-neuron traces around the context transition (**Fig. 3i, Extended Data Fig. 9a**) revealed persistent changes in activity aligned to the transition in every macro-region, indicating two distinct neural states corresponding to the two foraging environments. In individual sessions, these states were evident in the correlation structure and low-dimensional projections of population activity (**Fig. 3j,k**) and were not spuriously driven by slow-timescale fluctuations or drift artifacts (**Extended Data Fig. 9b**). To characterize the distribution of context-specific activity states across the brain, we identified single neurons that exhibited an activity change at the context transition which persisted throughout the block (**Fig. 3l, Methods**). Encoding of context identity was widespread, including in sensory areas (as expected through visual/odor cues), as well as subcortical, motor, and frontal areas. Because environment contexts are identified by continuous sensory information, we cannot disambiguate sensory responses from encoding of context-specific abstract variables (e.g. cognitive rules). Nonetheless, these data show that environment contexts recruit widely distributed activity states which could organize context-specific neural processes for decision computations.

### Foraging contexts organize population activity into separated subspaces

Although information indicating context identity was widespread, it remains unclear whether or how context alters the structure of decision-related activity (**Fig. 1a, 3g**). Decision-related activity in Stochastic and Deterministic contexts could lie within shared subspaces, indicating similar population-level structure (**Fig. 3m**, top); alternatively, it could be organized into separate subspaces, creating context-specific modes for sensory processing and/or decision formation (**Fig. 3m**, bottom).^7,25,42,43^ To first examine whether the overall structure of population activity is altered across contexts, we quantified the alignment of Deterministic- vs. Stochastic-context activity subspaces. We used PCA to identify orthonormal bases (i.e. subspaces) capturing within-context activity, and applied canonical correlation analysis^44,45^ (**Extended Data Fig. 10a**, **Methods**) to obtain vector pairs, and their canonical correlations, that geometrically capture subspace alignment. This provides measures of orthogonal dimensions between subspaces as well as overall subspace distance (e.g. the chordal distance, **Extended Data Fig. 10b-f**), which closely track an alternative metric based on explained variance.^46^ We quantified these for each session, normalizing by subtraction of within-context measurements.

Subspace distances (**Fig. 3n**) and orthogonal dimensionality (**Extended Data Fig. 10g**) were significantly higher than within-context controls, indicating that environment contexts organize population activity into separated subspaces. Interestingly, we observed substantial variability in subspace distances across brain regions, which was itself correlated with single-neuron coding of environment context (**Fig. 3o, Extended Data Fig. 10h**) and greater than expected from the effect of context coding itself (**Extended Data Fig. 10i,j**). This indicates that brain regions encoding context identity exhibit more separated activity subspaces, potentially supporting functional modes for decision making. Strikingly, subspace distances were also significantly correlated with behavioral measure of decision flexibility (**Fig. 3p, Extended Data Fig. 10k**), suggesting a close connection between subspace separation at the neural level and context-dependent cognition.

### Context-specific decision variables are encoded in orthogonal neural populations

To understand the neural basis of context-dependent foraging decisions, we next focused on how context shapes decision activity itself. One possibility is that context largely affects upstream processes, configuring inputs to a general decision process (**Fig. 1b**, **Fig. 4a**, *input configuration*).^11–14,20^ The neural signature of this would be context-invariant DV coding, as has been previously observed for rule-based perceptual choice. Context could also parse decision activity into context-specific coding vectors residing in separate subspaces (**Fig. 4a**, *subspace configuration*), which could enable modular learning and independent readout of decision computations.^7,8,25,27^ To distinguish these possibilities in our task, we examined DV coding in single-neurons and population-level geometry, which are related by coding correlation structure. That is, while correlated single-neuron DV coding corresponds to aligned population-level coding vectors and indicates input configuration (**Fig. 4b**, left), uncorrelated single-neuron coding corresponds to orthogonal coding vectors (**Fig. 4b**, right), indicating subspace configuration.

**Figure 4.**
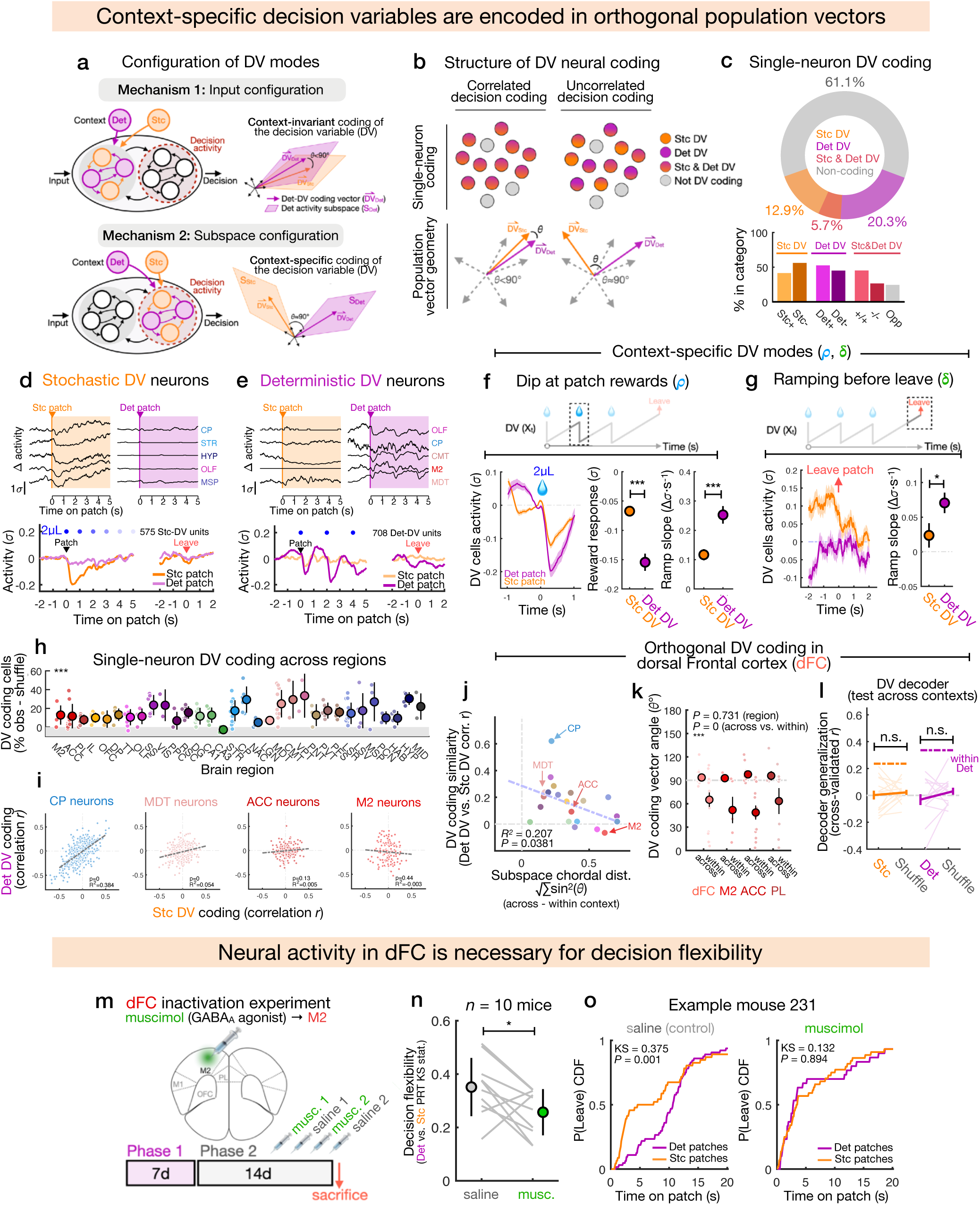
Context-specific integrator decision variables are orthogonally encoded in separate neural subpopulations. a) Illustration of **input configuration** vs. **subspace configuration** mechanisms for how context structures integrator dynamics for foraging decisions. Input configuration operates upstream of decision formation and preserves context-invariant decision dynamics, as observed in rule-based perceptual decision tasks. Subspace configuration exhibits context-specific decision coding vectors that lie in separate activity subspaces and may be orthogonal. b) Different possibilities for cellular-level coding structure underlying population geometry of decision activity. Orthogonal coding vectors (population level) correspond to uncorrelated single-neuron decision coding, which could be distributed in a common neural population or separate subpopulations. c) Top, percentage breakdown of Stochastic (Stc) decision variable (DV), Deterministic (Det) DV, both DVs, and non-coding neurons from among all recorded; bottom, breakdown of positive and negative DV coding cells within each category. For cells encoding both Stc and Det DVs, “opp” indicates opposite-sign coding across Deterministic vs. Stochastic contexts. d) Traces of trial-averaged activity for example Stc DV cells (top) and population average on Stochastic or Deterministic patch trials (bottom, *n* = 575 cells, *σ* = s.d. of firing rate). e) Traces of trial-averaged activity for example Det DV cells (top) and population averages on patch trials (bottom, *n* = 708 cells, *σ* = s.d. of firing rate). f) Left, average response of Stc DV (*n* = 575) and Det DV (*n* = 708) cells at patch rewards after trial initiation (at t > 0s, mean ± SEM); middle, comparison of reward response magnitudes across cell types (t-test, *P* = 3.37e-7); right, comparison of ramping slope (Δ activity *σ* / second) following reward response (t-test, *P* = 2.59e-4). g) Left, average response of Stc DV (*n* = 524) and Det DV (*n* = 622) cells around patch leave (subset of trials with no rewards in the 4 seconds preceding patch leave, mean ± SEM); right, comparison of ramping (Δ activity *σ* / second) slope before patch leave (t-test, *P* = 0.0392). h) Breakdown of single-neuron DV coding across regions (average % significant [α = 0.05 with Bonferroni correction] in Deterministic or Stochastic context above shuffle control, see **Supplementary Table 1** for region definitions, mean ± s.d., one-way ANOVA, *P* = 6e-4). i) Correlation (*r*) of Stc DV and Det DV coding across single neurons from example regions ([CP, Caudoputamen, *R^2^* = 0.385, *P* = 3.48e-56, *n* = 515]; [MDT, Mediodorsal Thalamus, *R^2^* = 0.056, *P* = 7.19e-7, *n* = 350]; [ACC, Anterior cingulate, *R^2^* = 0.009, *P* = 0.13, *n* = 268]; [M2, Secondary motor cortex, *R^2^* = 0.004, *P* = 0.44, *n* = 141]). j) Correlation across regions between normalized subspace distance and DV coding similarity (correlation *r* between Stc DV and Det DV coding), omitting regions with fewer than 50 DV coding neurons across sessions (linear regression, *R^2^* = 0.207, *P* = 0.0381, *n* = 21 regions). k) Angle (*ϴ* degrees) between Det DV and Stc DV coding vectors, and comparisons with within-context controls, for dFC neurons and each dFC region separately (mean ± SEM, t-tests against *ϴ* = 90°, *P* = 0.1444 [dFC], *P* = 0.2244 [M2], *P* = 0.0276 [ACC], *P* = 0.3598 [PL]; 2-way anova, *P* = 0 (within vs. between), *P* = 0.7308, region, *n* = 50 session samples). l) Performance of Stc DV and Det DV linear decoders to predict DVs from patch trials in the opposite unseen context (mean ± SEM, dashed lines indicate within-context decoder performance from **Extended Data** Fig. 11g, SR tests against shuffle controls, *P* = 0.6257, *n* = 14 sessions [Stochastic DV decoder], *P* = 0.3396, *n* = 13 sessions [Deterministic DV decoder]). m) Design of dFC inactivation experiments. After training in Phase 2 of the VR task, mice were injected unilaterally in the M2 region of dFC with saline or Muscimol (alternating across four days) and then tested in the task each day (see **Methods**). n) Comparison of differences between Deterministic and Stochastic PRT distributions (KS statistic, see also **Extended Data** Fig. 12f for comparison of Wasserstein metrics) under Muscimol vs. saline control (mean ± s.d., SR test, *P* = 0.0137, *n* = 10 mice). o) Deterministic and Stochastic PRT distributions, as the CDF of P(Leave)_t_, for example Mouse 231 under saline (left, KS test, *P* = 7e-4) and Muscimol (right, KS test, *P* = 0.8944), showing collapse of context-specific PRT behavior under M2 inactivation. Standard deviation (s.d.); Standard error of the mean (SEM); Kolmogorov-Smirnov test (KS test); Wilcoxon signed-rank test (SR test); \*\*\**P* < 0.001; \*\**P* < 0.01; \**P* < 0.05; n.s. *P* > 0.05.

We first quantified the correlation between single-neuron activity and model-fit DVs to identify single neurons encoding the integrator DV (**Methods**). 38.9% of all neurons encoded the DV on Stochastic patches, Deterministic patches, or both (**Fig. 4c**), with the majority specific to one context (12.9% Stc DV; 20.3% Det DV). Overlap between Stc and Det DV cells was smaller than within-context controls, and DV coding was not driven by movement or reward confounds (**Extended Data Fig. 11a,b**). From this analysis, we could readily identify single neurons that tracked the DV in one context but were unresponsive in the other (**Fig. 4d,e**, top), and averaging these yielded context-specific encoding of the Stochastic or Deterministic DV (**Fig. 4d,e**, bottom).

This shows that context organizes mixed coding of decision activity, largely in context-specific neural populations, consistent with subspace configuration (**Fig. 4a-b**). However, if context-specific DV neurons genuinely track different configurations of the DV, then they should additionally exhibit distinct dynamics on Deterministic vs. Stochastic trials, instantiating different configuration modes (***δ*** and **ρ, Extended Data Fig. 11c**). To test this, we examined whether: **1)** Det DV neurons exhibit faster ramping activity than Stc DV neurons (reflecting a larger drift rate ***δ***), and **2)** Det DV neurons exhibit larger reward responses (reflecting larger reward sensitivity **ρ**).

We first analyzed neural responses around patch rewards. Water rewards elicited stronger negative deflections in Det DV vs. Stc DV neurons and faster ramping after the reward, even across reward-matched trials (**Fig. 4f, Extended Data Fig. 11d**). We observed a similar difference in ramping after the first reward, but Stc DV neurons showed a larger response at trial initiation (**Extended Data Fig. 11e**). To eliminate the potential confound of extended reward responses, we also examined activity leading up to the leaving decision in trials with no preceding rewards – still, Det DV neurons exhibited faster ramping (**Fig. 4g**). These results demonstrate that context-specific DVs are encoded in neural populations which instantiate distinct integrator DV modes (i.e. configurations of ***δ*** and **ρ** in the model), potentially driving context-specific decision strategies.

### Neural activity subspaces structure context-specific decision coding

We next sought to investigate how encoding of foraging DVs is distributed across the brain. At the single-neuron level, we found that most regions contained detectable DV coding (**Fig. 4h, Extended Data Fig. 12a**), consistent with previous reports of distributed activity for perceptual decisions.^47,48^ However, while DVs were generally widespread, context-specific coding may be regionally graded or localized.^8^ To test this, we quantified DV similarity across contexts (for each region) as the correlation between single-neuron Stc DV and Det DV coding strength, and observed substantial heterogeneity in context-specificity (**Fig. 4i**): while some regions exhibited correlated DV tuning (e.g. Caudoputamen, CP), indicating context-invariant DV coding, others exhibited completely uncorrelated, context-specific DV coding (e.g. secondary motor cortex, M2).

How might this heterogeneity arise? One possibility is that the separation of activity subspaces decorrelates and orthogonalizes the coding structure of DVs across contexts. To test this, we examined the relationship across regions between overall subspace distance and DV coding similarity (**Fig. 4j**). Indeed, DV coding similarity was negatively correlated with subspace distance across regions, suggesting that subspace separation may parse decision activity into context-specific coding vectors. To further investigate this coding structure and its potential role in flexible foraging decisions, we narrowed our focus to a subset of exemplar regions in the dorsal frontal cortex (dFC – including M2, ACC, and PL), which exhibited high subspace divergence and uncorrelated DV coding, and have been linked to computations related to foraging.^38,39,49–51^

### dFC encodes orthogonal decision variables and supports context-dependent decisions

Focusing on the dFC, we first quantitatively tested whether context-specific DV coding vectors are geometrically orthogonal. A linear decoder analysis confirmed that both DVs could both be read out from dFC activity and generalized well within context (**Extended Data Fig. 12b,c**). If Stc and Det DVs are genuinely orthogonal, then their coding vectors should be: **1)** geometrically perpendicular (i.e. angle *θ* ≈ 90°), and **2)** mutually uninformative, indicating non-interfering readout of decision computations. We measured the angle *θ* between DV coding vectors and found these to be close to 90° and greater than within-context comparisons (**Fig. 4k**), indicating orthogonality. To rule out the possibility that this is driven spuriously by the concentration of measure in high-dimensional space, we repeated and confirmed the result in low-dimensional projections of population activity (**Extended Data Fig. 12d-f**). Further, while linear decoders were stable within context, they could not generalize across contexts (**Fig. 4l**), indicating that context-specific DVs reside in orthogonal coding vectors that can be readout independently.

As dFC receives input from areas that encode sensory context (**Fig. 3l**), we wondered to what extent this orthogonal coding structure could be driven by context-specific sensory responses in the same population. We tested this experimentally by recording dFC neurons in a separate cohort of naïve mice on the first 3 days of the task (**Extended Data Fig. 13a, Extended Data Fig. 8b**), which experience identical sensory conditions as expert mice. We analyzed subspace alignment in early vs. late sessions and observed a significant increase in subspace separation with experience (**Extended Data Fig. 13b,c**). Critically, single-neuron DV coding was initially correlated across contexts and only uncorrelated in experienced mice (**Extended Data Fig. 13d**), demonstrating that while integrator DVs are present in the naïve state, the organization of separated subspaces and orthogonal DV coding is learned and not an obligate consequence of simultaneous sensory coding.

This orthogonal DV structure positions dFC as a potentially key functional node for contextual configuration of foraging decisions. To test the causal contribution of dFC to decision flexibility, we injected the GABAA receptor agonist muscimol into M2 (a region within dFC) to locally silence neural activity (**Fig. 4m**). While M2 inactivation had no effect on average PRTs or engagement (**Extended Data Fig. 13e-f**), indicating intact foraging behavior, it significantly decreased decision flexibility compared with control sessions, at the group level and in individual mice (**Fig. 4n-o, Extended Data Fig. 13g**). Taken together, these results establish that dFC encodes orthogonal, context-specific DVs and is required for context-specific decision flexibility.

### dFC coding structure is preserved in real environments and organized over days

Thus far, our investigation has revealed a unique neural solution for context-dependent foraging decisions: environment contexts recruit separate activity subspaces that organize orthogonal integrator dynamics for decision computations (*subspace configuration*, **Fig. 1b, 4a**). At the same time, the artificiality and constraint of VR experiments make it difficult to address other questions that are critical for a comprehensive mechanistic picture. In particular, we wondered **1)** whether and how orthogonal DV coding generalizes to more naturalistic settings (e.g. when mice move freely in real environments), and **2)** how this coding structure is organized progressively as mice learn context-specific strategies.^21,22,30^ To address these questions, we returned to our original FM task which allows for unrestrained, volitional foraging (**Fig. 3a**, **5a**). We implanted chronic Neuropixels probes^52^ in the frontal cortex of four mice, recorded across daily sessions, and analyzed 2220 dFC units from 50 sessions across Phase 2 (**Fig. 5b, Extended Data Fig. 8c**).

**Figure 5.**
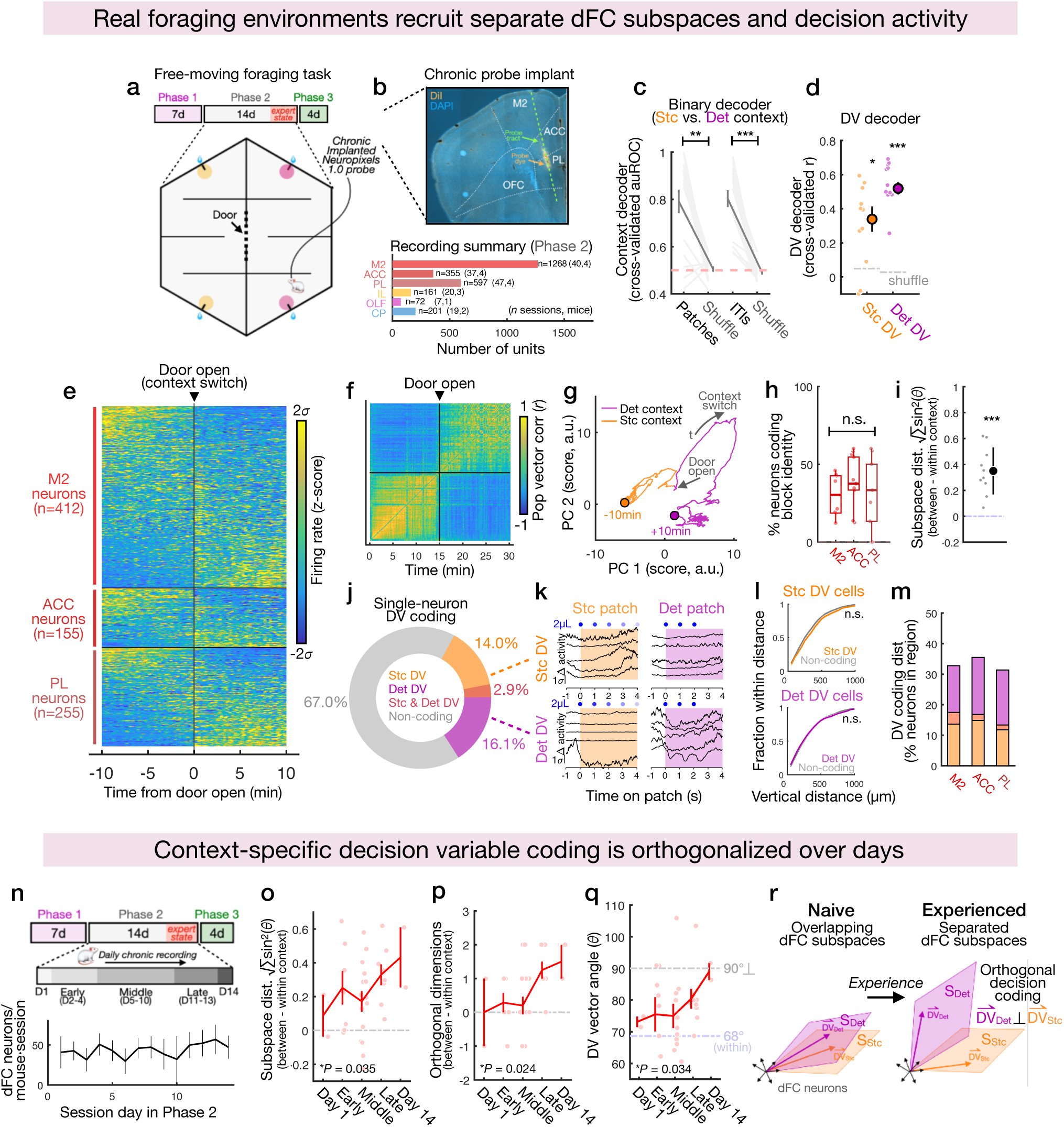
Orthogonal integrator dynamics in dFC are structured with experience and preserved in real foraging environments. a) Schematic of chronic Neuropixels recordings in during FM foraging task Phase 2 (Fig. 2c), in which mice forage in Deterministic and Stochastic environments during two 15-minute blocks, with one transition in the middle of the session when mice cross chambers through a controllable door. Expert sessions are defined as days 11+ (the final four days) in Phase 2. b) Top, example histology showing location of chronic Neuropixels probe implant targeting the dorsal frontal cortex (dFC; orange, DiI probe tract; blue, DAPI); bottom, summary of recordings from all Phase 2 sessions. A total of *n* = 2220 dFC neurons were recorded from secondary motor cortex (M2), anterior cingulate (ACC), and prelimbic cortex (PL) across 50 sessions in 4 mice. c) Performance of binary decoders (Fisher’s linear discriminant) predicting Deterministic vs. Stochastic context identity from dFC population activity during patch trials (left) or inter-trial intervals (ITI, right), quantified using the area under ROC curve (cross-validated auROC) for predicted vs. true labels on held-out data (mean ± SEM, SR test against shuffle, *P* = 1.3e-3, trials, *P* = 4.378e-4, ITIs, *n* = 16 sessions). d) Performance (cross-validated correlation *r*) of linear decoders to predict Stochastic or Deterministic DVs from dFC activity on held-out trials (mean ± SEM, SR tests against shuffle, *P* = 0.034, Stochastic DV, *P* = 4.88e-4, Deterministic DV, *n* = 12 sessions). e) Heat map of single-neuron activity centered around the time of door opening/environment transition for each session (dFC neurons in expert sessions, *σ* = s.d. of firing rate). f) Correlation matrix of dFC population vectors sampled across an example session. Block structure indicates within-context similarity reorganized around the environment transition. g) Principal component (PC) projection of activity from example session in (Fig. 5f) showing persistent change in population activity upon context transition (arrow shows direction of time). h) Fraction of neurons encoding Stochastic vs. Deterministic block/environment identity across dFC regions (median IQR and range, one-way Anova, *P* = 0.5254, *n* = 12 sessions). i) Normalized (between -within context) substance distance for dFC population activity in Deterministic vs. Stochastic environments (mean ± s.d., t-test, *P* = 1.74e-4, *n* = 10 sessions). j) Breakdown of Stc DV (orange), Det DV (purple), both DVs (red), and non-coding neurons across dFC recordings from expert sessions (*n* = 822 neurons, permutation test at α = 0.05). k) Trial-averaged Stochastic and Deterministic patch activity for example context-specific Stc DV (top) and Det DV (bottom) neurons (*σ* = s.d. of firing rate, Δ from pre-trial baseline). l) Spatial clustering of Stc DV (top) and Det DV (bottom) neurons. For each cell type, the fraction of the full population (i.e. Stc DV cells, Det DV cells, or non-coding cells) contained within an expanding distance (µm) from individual sample cells. Stc DV and Det DV cells have the same spatial clustering (fraction as a function of distance) as non-coding cells (KS tests, *P* = 0.9923 [Stc DV vs. non-coding cells], *P* = 1 [Det DV vs. non-coding cells]). m) Breakdown of DV coding cell categories as in (Fig. 5j) for M2, ACC, and PL. n) Top, schematic of chronic recordings across the entire Phase 2 time-course, binned into day 1, early sessions (days 2-4), middle sessions (days 5-10), late sessions (days 11-13), and the final day 14. Bottom, average number of dFC neurons recorded per session and mouse for each day in Phase 2 (mean ± s.d., *n* = 50 total sessions). o) Subspace distance across contexts in dFC populations (normalized to within-session, as in Fig. 5i) across days during Phase 2 progression (mean ± s.d., linear regression against time bins, *P* = 0.035). p) Normalized number of orthogonal vectors across contexts in dFC populations across days during Phase 2 progression (mean ± s.d., linear regression against time bins, *P* = 0.024). q) Angle (*ϴ*°) between Stc DV and Det DV coding vectors in dFC populations across days during Phase 2 progression. Grey line indicates orthogonality at 90°, and light blue line indicates the average value of within-context control comparisons (*ϴ* ≈ 68°, see also **Extended Data** Fig. 13d**,e**, mean ± s.d., linear regression against time bins, *P* = 0.034). r) Cartoon illustration of progressive subspace divergence in dFC across days from the naive state (more overlapping subspaces) to the experienced state (separated subspaces and orthogonal DV coding vectors). Standard deviation (s.d.); Standard error of the mean (SEM); Wilcoxon signed-rank test (SR test); \*\*\**P* < 0.001; \*\**P* < 0.01; \**P* < 0.05; n.s. *P* > 0.05.

We focused first on recordings from expert mice and tested whether dFC neurons encoded environment context and integrator DVs in the FM setting. Decoders could predict context identity from dFC activity (**Fig. 5c**) and DVs could be read out in both contexts (**Fig. 5d**). Applying the same analyses as for the VR task, we observed strikingly similar large-scale neural structure: distinct activity states for Stochastic and Deterministic contexts were evident in single-neuron traces (**Fig. 5e**), population vector correlations (**Fig. 5f, Extended Data Fig. 14a**), and low-dimensional projections of population activity (**Fig. 5g**) following similar dynamics around the context transition (**Extended Data Fig. 14b,c**). Single neurons encoded environment identity across dFC regions (**Fig. 5h**), and subspace distances across environment contexts were larger than within-context comparisons (**Fig. 5i**). These observations raise the possibility that the orthogonal DV coding structure may also be preserved for foraging in real environments. Indeed, even across spatial contexts (which contain no explicit visual or odor cues), we found that most single neurons (91.2% of DV coding neurons) were context-specific (**Fig. 5j**), exhibiting integrator dynamics only in the Deterministic or Stochastic environment (**Fig. 5k**). DV neurons were not spatially clustered (**Fig. 5l**) and were similarly distributed across dFC regions (**Fig. 5m**).

How is this neural structure organized as mice learn context-specific decision strategies? To answer this, we leveraged our chronic recording dataset to analyze the evolution of dFC population activity across the full two-week time-course of Phase 2 (**Fig. 5n**). For each session, we computed subspace distance measures and the angle *θ* between Stochastic and Deterministic DV coding vectors. Remarkably, activity subspaces were initially overlapping and only progressively separated over several days with task experience (**Fig. 5o,p**). Further, while within-context DV coding vectors remained stable and smaller than 90° apart (mean = 67.9°, **Extended Data Fig. 14d**), context-specific coding vectors were continuously separated over time and nearly orthogonal by the final day of Phase 2 (**Fig. 5q-r, Extended Data Fig. 14e**). Taken together, these results establish orthogonal integrator dynamics as a general neural solution for context-dependent foraging decisions which, in the dFC, is organized with experience over days and preserved across both highly controlled (VR) and unconstrained naturalistic (FM) experiment settings.

### Rapid reconfiguration of dFC population dynamics supports volitional mode switching

One advantage of this neural structure may be to support rapid switching between cognitive modes: one learned, different configurations of the decision process instantiated in orthogonal coding vectors could be recruited selectively by control signals, potentially with little switching cost. Indeed, in Phase 3 of the FM task, mice repeatedly switch between decision strategies and do so immediately on context transitions without relying on recent experience (**Extended Data Fig. 2h-j).** Yet it has been extremely difficult to study such rapid switching at the neural level, as typical experimental constraints (such as block structure) make context transitions rare and often eliminate the volitional aspect of strategy switching. We took advantage of the unique design of Phase 3 (**Fig. 6a**), in which mice were allowed to switch environments freely, and we continued to record dFC activity through these sessions, retaining 712 units across 16 sessions from the same animals (**Fig. 6b**). Despite mice now switching contexts repeatedly (every 20-30 seconds, **Extended Data Fig. 1k**), we still observed robust encoding of context identity (**Fig. 6c**), integrator DVs (**Fig. 6d**), and orthogonal DV coding vectors (**Fig. 6e, Extended Data Fig. 14f,g**) in dFC, suggesting that mice rapidly toggle between decision modes on volitional context switches.

**Figure 6.**
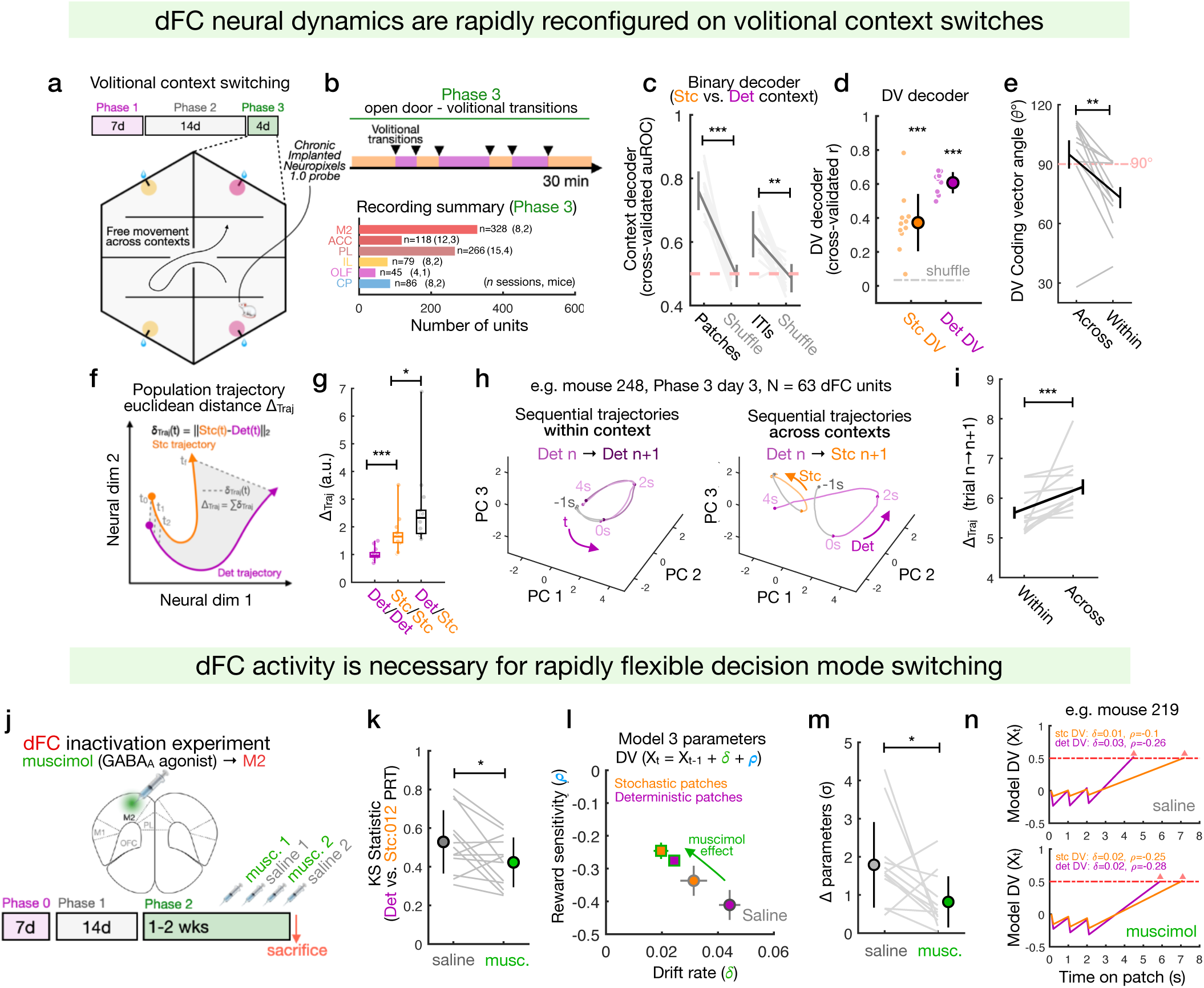
dFC dynamics are rapidly reconfigured on volitional context transitions and support flexible mode switching. a) Schematic of chronic Neuropixels recording in Phase 3 of the free-moving (FM) foraging task when mice are free to switch environments repeatedly and volitionally through the open door. b) Cartoon of volitional transitions in Phase 3 and recording summary (*n* = 712 dFC neurons, 16 sessions, 4 mice). c) Performance (cross-validated auROC) of Fisher’s discriminant binary decoders for context identity on patch trials (left) and inter-trial interval (ITI) periods (right) (mean ± SEM, SR tests against shuffle, *P* = 4.88e-4 [trials], *P* = 1.5e-3 [ITIs], *n* = 12 sessions). d) Performance (cross-validated correlation *r*) of Stc DV and Det DV decoders compared to shuffle controls (mean ± SEM, SR tests against shuffle, *P* = 4.88e-4 [Stc DV], *P* = 4 .88e-4 [Det DV], *n* = 12 sessions). e) Angle *ϴ* between context-specific DV coding vectors compared to within-context controls (SR test, *P* = 0.0034, *n* = 12 sessions, see **Methods**). f) Illustration of neural trajectory analysis and quantification of ΔTraj (the sum of time point-matched Euclidean distances between patch trial trajectories in dFC populations). g) Comparison of ΔTraj across within- and across-context trial-average trajectories (median IQR and range, U tests, *P* = 4.78e-4 [within Det vs. within Stc], *P* = 0.01 [within Stc vs. across contexts], *n* = 12 sessions). h) Average of trajectories for Det→Det (left) and Det→Stc (right) sequential trial pairs from one example session (Mouse 251, session 17); arrow indicates direction of time. i) Comparison of session average ΔTraj for within- vs. across-context sequence pairs as visualized for one session in Fig. 6h (median ± SEM, SR test, *P* = 9.76e-4, *n* = 12 sessions). j) Schematic of dFC Muscimol inactivation experiments and timeline. Muscimol or saline were injected unilaterally into M2 on alternating days after mice were trained through the full task (Fig. 2c) including several days in Phase 3. k) Comparison of Deterministic vs. Stc:012 PRT distributions (KS statistic) for Muscimol vs. saline sessions (mean ± s.d., SR test, *P =* 0.0327, *n* = 13 mice). l) Scatter plots of average parameter fits (*ρ* and *6*) for Stochastic (orange) and Deterministic (purple) patches under saline (grey outline, circles) and Muscimol (green outline, squares, means ± SEM). Muscimol reduces the magnitude of both parameters and the difference between them across contexts (quantified in Fig. 6m) m) Comparison of parameter differences (Euclidean distance of z-scored *ρ* and *6* values) for saline vs. Muscimol sessions (mean ± s.d., SR test, *P* = 0.0479, n = 13 mice). n) DV rollouts for a Deterministic/Stc:012 reward sequence (rewards at t = 0, 1, 2s) using parameter fits from an example Mouse 219 for saline (top) or Muscimol (bottom) sessions, visualizing the collapse of context-specific DV configurations. Standard deviation (s.d.); Standard error of mean (SEM); Mann-Whitney U test (U test); Wilcoxon signed-rank test (SR test). ***p<0.001; **p<0.01; *p<0.05; n.s. p>0.05.

To study this rapid reconfiguration at level of neural dynamics, we analyzed population trajectories during patch trials within and across contexts by measuring the Euclidean distance between them (Δ_Traj_, **Fig. 6f**). We confirmed that context-specific trajectories were more separated than within-context ones (**Fig. 6g**). We then examined trajectories for sequential cross-context (e.g. Det → Stc) trial pairs to see the immediate effect of context transitions on neural population dynamics. Visualizations showed divergence of paired-trial trajectories across contexts (**Fig. 6h**), and Δ_Traj_ for cross-context trials was greater than within-context comparisons (**Fig. 6i**). This shows that dFC dynamics do not gradually update over multiple trials but, mirroring the behavior, are immediately reconfigured upon volitional transitions, potentially enabling rapid mode switching.

Finally, to test whether volitional mode switching depends on dFC activity, we used muscimol to silence M2 neurons during the FM task (**Fig. 6j**). Like in VR, decision flexibility was significantly reduced under muscimol compared to saline (**Fig. 6k**). In the integrator model, this behavioral pattern should reflect a collapse of parameter configurations underlying distinct decision modes. Indeed, we found that ***δ*** and **ρ**, and their difference across contexts, were reduced under M2 inactivation (**Fig. 6l,m**), collapsing DV configurations (**Fig. 6n**). While mice still foraged effectively, M2 inactivation also decreased context transition rates and increased PRTs (**Extended Data Fig. 15**), suggesting a potential role for dFC in volitional strategy shifting for foraging.^18,53–55^ Our observation of longer PRTs under dFC silencing is also consistent with a role in waiting behavior and sustaining persistent activity for decision and motor timing.^39,47,49,50,56–59^

## Discussion

A comprehensive understanding of flexible cognition depends on mechanistic insight into how neural processes are contextually reconfigured to solve the diversity of problems that animals encounter in the natural world.^1,4^ We studied the neural basis of context-dependent decision making in the ecological setting of patch foraging. Our approach stands in contrast to classical task designs, which require animals to learn sensory cues or infer rules for perceptual choices, and mostly identify mechanisms of context-dependency that operate upstream of decision formation.^11–13,17,20^ Instead, we observed a strikingly different solution for foraging across naturalistic environments: a *subspace configuration* mechanism, wherein different contexts recruit separate activity subspaces that organize orthogonal coding of integrator decision variables (DV). In our foraging task, context-specific neurons encoded distinct configurations of an integrator DV that computes patch leaving decisions. In the dorsal frontal cortex (dFC), these orthogonal integrators are organized with experience over days, rapidly toggled by volitional context switching, and necessary for flexible expression of context-specific decision strategies. Together, these findings reveal orthogonal integrator dynamics as a general neural solution for context-dependent foraging decisions, deepening mechanistic insight into the biological basis of flexible cognition and setting a new foundation to study mammalian cognitive capacities for real-world ecological challenges.

### Configurations of a common integrator process explain context-specific foraging decisions

Recent examination of classical problems in behavioral ecology through the lens of decision theory^35,36^ has begun to clarify the algorithmic logic and neural basis of foraging decisions,^29,30,60^ raising timely questions about how neural dynamics are configured to support flexible cognition for ecological problems. We recently demonstrated that patch leaving decisions in mice are well described by an integrator process oppositely weighing time and rewards,^38^ and we and others have identified neural correlates of integrator DVs in the frontal cortex of rodents.^39,47,49,50,61,62^ For patch foraging, neural integrators can instantiate different decision algorithms, such as tracking time investment, time since rewards, relative reward rates, and reward quantity (**Fig. 2i,k**).^35,63^

Inspired by a ubiquitous challenge in the natural world, we wondered how animals adapt this cognitive process across different foraging environments. We studied mice foraging for water rewards across two environment contexts defined by distinct reward dynamics: a Stochastic context, yielding decaying-probability rewards, and a Deterministic context yielding fixed rewards (**Fig. 2b,c**). Mice calibrated patch leaving decisions to Deterministic reward structure (leaving after the final reward, **Extended Data Fig. 1f,g**), suggesting a possible “counting” algorithm (i.e. ***ρ*** > 0, Model 4) and raising the hypothesis that mice may change their cognitive algorithm for patch foraging across Deterministic and Stochastic environments. Surprisingly, we found that mice use the same reward integration algorithm (Model 3) under different parameter configurations (**Fig. 2m-o**). Model fits to Deterministic patches had larger magnitudes for both the drift rate **δ** and reward sensitivity ***ρ***, and critically, these configurations were reflected in the neural correlates of DVs (**Fig. 4d-g**). This suggests instead that, rather than switching algorithms, reconfiguration of a common computation weighing reward gains against time may be a general solution for flexible foraging. Future studies using different reward distributions should clarify how different configurations of integrator dynamics can be tuned to express adaptive decision strategies.^35,63^

What biological mechanisms underlie changes in configuration of the integrator process? By analogy to bounded accumulator (e.g. drift-diffusion) models for perceptual decision making, the drift rate **δ** in our model effectively captures the weight of passing time as evidence *for* patch leaving. In other task settings, drift rates for evidence accumulation can be modulated by the quality of sensory evidence, confidence, and value.^37,38,64–67^ For foraging decisions, tuning of drift rates may be driven by top-down cortical signals (e.g. from frontal areas),^50,56^ organization of recurrency sustaining persistent activity,^14,68^ and neuromodulators such as serotonin, which has been linked to behavioral patience for uncertain rewards.^69–71^ The reward sensitivity ***ρ*** in our model can be thought of as a weight of pulsatile evidence *against* patch leaving, a signal which is likely mediated by detection of water in the mouth through somatosensory pathways. The gain (or sign) of reward signals onto the integrator could be modulated by sensory gating and/or local microcircuit architectures. In general, what mechanisms control the parameters of integrator dynamics, and how these are learned and calibrated from reward feedback, are important questions for future investigation which can be readily tackled with our experimental paradigm.

### Subspace configuration of context-specific orthogonal integrator dynamics

The central aim of our study was to understand how neural dynamics are structured to support context-dependent decision computations. To express contingent behavior, context signals must configure neural network modes to alter how sensory information is integrated to form a decision (**Fig. 1a, 3g**).^8,9^ A critical mechanistic distinction is whether context-dependency is achieved by altering the inputs to a *common decision process* (“input configuration”) or by altering *the decision process itself* (“subspace configuration”, **Fig. 1b, 4a**). For rule-based perceptual choice problems, input configuration mechanisms (e.g. sensory gating or attentional shifting) are important and highly diverse. For example, a foundational study in monkeys suggested a mechanism that operates by tuning frontal cortex recurrent network dynamics to contextually gate the effect of sensory input on context-invariant decision activity for binary perceptual choices.^11^ A major recent advance has been the unification of such mechanisms in a framework that explains individual behavior variability as idiosyncratic expression of input configurations for a common decision process.^20^

However, an important limitation of input configuration models is that they are agnostic about the structure of decision coding itself. As hypothesized in other work, the brain could also use separate activity subspaces to organize context-specific instantiations of a decision process (**Fig. 1b, 4a**).^7,23,25,72^ Subspace coding has been studied in many domains of neuroscience, and while it appears to be crucial for decorrelating sensory representations,^25,73^ cortical communication,^23^ and shielding premotor activity from motor expression,^46,47,74^ it has not been observed as a coding structure for cognitive decision variables. Nonetheless, orthogonal coding of context-specific DVs could confer computational and behavioral benefits, including modular tuning during learning, independent readout by downstream circuits, and rapid toggling by context control signals.^7,27,42,75^

In contrast to previous studies, we find robust and convergent evidence of a subspace configuration mechanism structuring orthogonal coding of context-specific DVs in mice (**Fig. 4c-l, Fig. 5j-r**, **Fig. 6e**). In part, integrator DVs for foraging appear to be instantiated in largely separate neuron subpopulations – a unique subspace organization which effectively maps coding dimensions onto anatomical cell identities (**Fig. 4c-e**, **Fig. 5j,k**). This simple structure may allow the brain to calibrate and toggle context-specific cognitive modes by activating subsets of neurons in different contexts, expressing and exposing them selectively to teaching signals and plasticity. Indeed, rather than incurring a cognitive “switching cost,” typically observed as a performance decrease after context transitions, mice in our foraging task immediately toggle decision strategies when they volitionally switch environment contexts (**Extended Data Fig. 2h-j**). As this is mirrored in the immediate reconfiguration of dFC dynamics and depends causally on dFC activity (**Fig. 6h-n**), the subspace coding structure may support rapid strategy switching through this modular representation – though more precise causal studies are needed to test this hypothesis.

Given this departure from the extant literature, our findings beg the question: *why* do we observe mostly context-specific, rather than context-invariant, decision coding? Although our task design is unique in its use of environments to study context-dependency (**Extended Data Fig. 6**), we experimentally ruled out the possibility that orthogonal DVs are a spurious consequence of confounds from the sensory environment (**Fig. 5n-r, Extended Data Fig. 13a-d**). Rather, neural solutions for context-dependent decisions are likely sensitive to the nature of the cognitive problem and task setting.^8^ For rule-based perceptual choice, tuning of sensory inputs can embed context contingencies, and perhaps especially if context or rule cues are transient, unimodal stimuli.^11–13,18,20^ In our foraging task, however, context information is continuously available and is identified unambiguously by the sensory-spatial environment. Additionally, from the animal’s perspective the notion of context as a stable environment may be importantly different from sensory cues signaling cognitive rules, as such stimuli rarely carry abstract meaning in the natural world.^21^

We speculate that when context is an environment, which carries intrinsic ecological meaning (e.g. a place to explore and forage), the brain may use it to organize large-scale neural states that can shift behavior and cognition between specific functional modes adapted for each environment. Indeed, we found that environment context identity was not only encoded in sensory areas, as expected from the task design, but widely throughout the brain, including in frontal, subcortical, and thalamic areas.^8,73,76,77^ Foraging environments also recruited separate activity subspaces in many regions (**Fig. 3m-p**), organizing orthogonal DV coding (**Fig. 4j**) and likely reflecting context-specific population dynamics that may support other aspects of sensory processing and behavior. These findings are consistent with a picture of context-sensitivity and neural “configurability” as graded in the brain rather than localized.^8^ At the same time, some circuits, including in the frontal cortex, may be of particular importance for regulating cognitive modes.^1,3^

### Placement of the dorsal Frontal cortex in a network of configurable circuitry

We used the VR task to record widely across 32 brain regions and characterize how context broadly shapes neural activity (**Fig. 3-4**). We then focused on the dFC to study the organization, generality, and function of orthogonal DV coding (**Fig. 5-6**). We found consistently that inactivation of M2 (the secondary motor/agranular frontal cortex in dFC) disrupted context-dependent foraging decisions (**Fig. 4m-o**, **Fig. 6j-n**). Interestingly, M2 inactivation did not disrupt general foraging behavior in either task, suggesting redundancy for the core behavior. This is perhaps not surprising given the distributed nature of integrator dynamics for decisions and movement timing,^47,48,59,78,79^ which in rodents are sustained through multi-region recurrent interactions. However, this result suggests a more specific function for M2 in selecting and/or stabilizing context-specific cognitive modes – consistent with previous studies linking this cortical region to flexible behavior, action policies, and integrator dynamics based on sensory contingencies^19,59,80–82^

The anatomy also supports this picture, as M2 receives direct input from sensory areas and projects broadly to motor control circuits in the cortex, midbrain, and basal ganglia.^49,51,59,61,78^ As many other regions exhibited context-specific subspaces and DV coding for foraging (**Fig. 4j**), our findings focus questions on multi-region interactions and the microcircuit mechanisms underlying selection and maintenance of context-specific neural dynamics. These can be readily tackled with our paradigm using high-density recordings^41,52^ in molecularly-defined dFC microcircuit elements and input/output pathways, and by coupling these with targeted circuit perturbations.

### Ecological grounding and fluid methodology for the study of flexible cognition

Finally, our approach bridges the limitations of both “reduced” and “naturalistic” task designs (**Fig. 3a**), demonstrating the scientific advantage of fluid methodology and the importance of ecological grounding to drive robust cognitive behavior.^21,22,40^ We departed from classical methods to design a foraging task that encourages context-dependency for an intuitive natural problem.^29,30,40^ In the foraging setting, mice readily learned context-specific decision strategies within days (instead of weeks or months) and could rapidly, repeatedly, and volitionally shift cognitive modes. Practically, this allowed us to widely sample neural activity across regions with single-cell resolution and study neural dynamics over the entire time-course of learning, and across experimental paradigms, leading to key insights that have eluded previous studies. Conceptually, our approach shows how complementary tasks can be used synergistically to clarify, validate, and expand mechanistic models of brain function. The dual-task approach using VR and FM designs established generality of orthogonal integrator dynamics across levels of ecological complexity. Importantly for future work, our findings also expand the scope of inquiry around the neural basis of flexible cognition, opening a new avenue to study how decision processes are shaped by natural environments, adapt to changing conditions, and are dynamically reconfigured to meet real-life ecological challenges.

## Methods

### EXPERIMENTAL MODEL AND SUBJECT DETAILS

#### Mice

A total of 72 adult mice (C57/BL6j, 35 male, 37 female, age 2-6 months) were used in the experiments. Mice were housed on a 12h dark/12h light cycle (dark from 07:00 to 19:00) and performed behavior tasks in the dark period. All procedures were performed in accordance with the National Institutes of Health Guide for the Care and Use of Laboratory Animals and approved by the Harvard Animal Care and Use Committee.

### METHOD DETAILS

#### Surgical procedures

For virtual-reality (VR) and muscimol inactivation experiments, a custom titanium head-plate was attached to the skull for head-fixation. Mice were anaesthetized using isoflurane (4% induction, 1-2% maintenance) and a head-plate was attached to the skull with Metabond (Parkell).^83^ For both acute and chronic Neuropixels recordings, fiducial marks were made at the target sites for probe insertion using a fine-tipped pen, and a ground pin was inserted into the skull above the cerebellum and attached with Metabond. For muscimol inactivation experiments, a craniotomy was placed above the secondary motor cortex (M2, Bregma +2.35mm AP, +1.0mm ML) region of the dorsal frontal cortex (dFC) and muscimol or saline was injected 0.8mm ventral to the skull surface.

For chronic Neuropixels implants, mice were anaesthetized using Ketamine/medetomidine (Ketamine 10 mg/mL, Dexdomitor 0.04 mg/mL, 6 mL/kg). We used a custom adapted version of the Apollo implant described in Bimbard et al.,^52^ which uses separable “docking” and “payload” modules to secure and cement the probe to the skull after insertion and recover the probe after completion of experiments. Neuropixels 1.0 single-shank probes were used for chronic recordings in the frontal cortex, and we tracked electrophysiology signals online using SpikeGLX to guide probe placement. After implanting and securing the probe, we injected Antisedan (Atipamezole hydrochloride, 5mg/mL) to reverse the effects of Ketamine and allowed animals to fully recover for 1-2 weeks before behavior experiments.

#### Foraging behavior experiments

For all experiments, mice were water restricted to maintain body weight > 85% of their baseline weight before training to ensure task motivation and maintain health. Additional water supplementation was provided following training sessions with a total of task and post-task water ranging from 0-3mL per day, determined individually per mouse. Mice that showed task motivation at higher weights were given greater amounts of water supplementation accordingly to minimize stress from water restriction. Body weights ranged from 85-105% of baseline.

Male and female mice were used in approximately equal proportions for all behavior and recording experiments. We observed no significant differences in task engagement, general foraging behavior, or context-dependent foraging decisions across sexes (**Extended Data Fig. 5**).

#### Free-moving (FM) patch foraging task

##### Arena setup for FM foraging task

Mice navigated a hexagonal arena (inner radius 26.35 cm) with waterspouts placed at four locations in the center of the arena walls (see **Fig. 2c**) and a wall with a controllable door separating the arena into two chambers (the “Stochastic” and “Deterministic” environment contexts, each with two waterspouts). Mice were detected at waterspouts using IR beam breaks (3mm LED IR beam sensors, Adafruit), and water timing and amount was controlled using solenoid valves (LHDA 1221111H, The Lee Company). Task events (water delivery and door separating the chambers) were controlled and recorded using custom Arduino code running on a microprocessor (Teensy 4.1). Transparent acrylic barriers (walls schematized in **Fig. 2c**) were placed between the patches to increase and equalize the effective path length between patches (approximately 30 inches or 76.2 cm) and encourage greater time investment for patch trials. Mice could approach and nose-poke a reward port to initiate a trial, upon which they would immediately receive a 2µL water droplet (marking t = 0s in the patch trial). If they remained with their nose inserted, they could continue to receive water rewards each second with decreasing probability based on patch reward statistics (see below) and were free to leave at any time by removing their nose. After leaving a patch, that reward port was timed out and would not initiate another trial for 6 seconds. All rewards were 2µL water droplets.

##### Habituation

Mice began water restriction one day prior to the task and were handled for 15 minutes prior to starting experiments. Mice were not habituated to the arena before day 1.

##### Phase 1

For the first 10 sessions (30 minutes each), mice were exposed only to the right chamber (the Deterministic environment context - the door separating the chambers remained closed) which included two reward ports yielding Deterministic reward returns. After the initial reward at t = 0s, mice could receive additional rewards at t = 1 and t = 2s (for a total of three guaranteed rewards, **Fig. 2b**, bottom). Mice shaped their patch leaving decision strategies to this Deterministic reward structure during Phase 1 (**Extended Data Fig. 1d-g**).

##### Phase 2

Mice then proceeded to Phase 2, in which they were exposed to the Deterministic environment (the right chamber) and the Stochastic environment (the left chamber) once per session in 15-minute blocks, alternating the starting environment each day. After 15 minutes in the session, the door separating the chambers opened, and once the mouse explored the other chamber and completed a patch trial, the door was automatically shut, imposing a block structure. On Stochastic patches, after an initial reward at t = 0s, mice could receive additional rewards each second with decaying probability. Initial P(Reward) P_i_ at t = 1s for Stochastic patches was drawn randomly (uniformly) from P_i_ = [0.66, 0.81, 0.93, 0.99] and followed an exponential decay with ***τ*** = 0.325s (***P_t_*** = ***P_i_e***^−***τt***^), with P_t_ = 0 after 20 seconds (**Extended Data Fig. 1a**). 15% of the patches in the Stochastic environment were programmed with the Deterministic patch return (“Stc:012 patches” with rewards at t = 0, 1, 2s) to use as reward-matched probe trials. Mice experienced the Phase 2 task structure for 2 weeks, typically with 1 day off per week (6 days on, 1 day off).

##### Phase 3

After Phase 2, mice transitioned to Phase 3 of the task, in which the door separating the environments/chambers was always left open. Mice were therefore free to switch contexts and forage completely under their own volition. Although mice gained more rewards on average at Deterministic compared to Stochastic patches and preferred to exploit Deterministic patches (**Extended Data Fig. 1h,i**), they still explored both contexts extensively in every session, typically making 1-4 context transitions each minute (**Extended Data Fig. 1j,k**). Despite the lack of block structure or designed context cues, mice still exhibited the same differences in decision behavior as observed in Phase 2 and altered their behavior immediately on context transitions (**Extended Data Fig. 2h-j**), indicating rapid toggling between decision strategies.

For the alternative hidden state task variant (**Extended Data Fig. 3**), mice (2 males, 3 females) were only exposed to one chamber with two ports, and context was a hidden state that dictated the patch statistics (Deterministic or Stochastic) but was never cued. Phases 1 and 2 had identical block structure as the main task, and Phase 3 had alternating blocks of approximately 30 seconds to closely replicate the frequency of context transitions mice exhibit volitionally in the main task.

#### Virtual reality (VR) patch foraging task

##### Setup for VR foraging task

Virtual reality setups were identical to those used in Bukwich et al.^38^ Three monitors (width 53cm, height 30cm) were placed in front and on either side of the animal. Virtual reality scenes were generated using VirMEn software in MATLAB on a workstation computer (DELL Precision 5810). Mice were head-fixed on a cylindrical Styrofoam treadmill (diameter 20.3cm, width 10.4cm) and the rotational velocity of the treadmill was recorded using a rotary encoder. The output pulses of the encoder were converted into velocity signals using custom Arduino code running on a microprocessor (Teensy 3.2) and integrated within VirMEn to compute position. Water was delivered to mice from a waterspout placed in front of their mouth, and licks were monitored using an infrared sensor (OPB819Z, TT electronics). Voltage signals from the rotary encoder were digitized and recorded on the virtual reality computer using a data acquisition system (PCIe-6323, National Instruments). Water timing and amount was controlled using a solenoid valve (LHDA 1221111H, The Lee Company) and a switch (2N7000, On Semiconductor), with TTL pulses generated by the virtual reality computer via the PCIe-6323 data acquisition card.

##### Habituation

Mice were habituated to the treadmill prior to initial head-fixation with delivery of occasional water droplets from the reward spout while mice were free to explore on top of the treadmill, which was held in place to prevent rotation. Once mice showed consistent interest in consuming water droplets (usually 2 days with 15-minute exposures), head fixation began on the following session. Initial head-fixation habituation sessions were run for durations of 15 minutes, with gradual increases in session duration across consecutive sessions reaching a maximum duration of 50-60 minutes. During these sessions, water rewards (2µL) were delivered in 10 second intervals with additional rewards given during running bouts to reinforce running behavior. Once starting head-fixation, mice performed daily sessions with 1 day off per week (6 days on, 1 day off) for the remainder of the task. Once mice were consistently running for most of the session (usually 3-5 days), they proceeded on the next session to Phase 1 of the VR foraging task.

##### Phase 1

Once mice demonstrated consistent running behavior during habituation (above), they began training on the linear virtual track. Mice navigated forward down the virtual corridor and could stop at visually indicated patch locations to receive water rewards (they never received water between patches). The VR gain was set so that the travel distance between patches was 50cm. On patch entry, mice received a context-specific odor puff indicating that they were on the patch, and if they stopped moving for 0.05s, could receive an initial water reward (2µL). Following this reward, they could stay on the patch to receive additional rewards following patch-specific statistics. For the first 7-10 days, mice were only exposed to the Deterministic environment context, which was indicated with a unique visual pattern on the corridor walls (between the patches), unique visual landmarks for patch locations, and a unique patch odor (isoamyl acetate or ethyl butyrate). Deterministic patches always yield the patch-initial reward (at t = 0s) followed by rewards at t = 2s and t = 4s for a total of three guaranteed rewards. Mice could leave the patch at any time by moving forward on the treadmill to advance. The immobile time required to initiate the patch trial (starting at 0.05s) was gradually increased by 0.05s each day to a maximum of 0.5s. Occasionally, this was reduced for animals that showed faltering engagement rates and then increased again to 0.5s as engagement recovered.

##### Phase 2

Mice then proceeded to Phase 2 of the task, where they foraged in the Deterministic and the Stochastic environment once per session in two blocks (20-25 minutes each) with one context transition in between. The Stochastic environment was indicated with context-specific visuals and odors (isoamyl acetate or ethyl butyrate) on patch entry. On Stochastic patches, after an initial reward at t = 0s, mice could receive additional rewards each second with probability P(Reward) that decayed over time (**Extended Data Fig. 4a**). Initial P(Reward) P_i_ at t = 1s for Stochastic patches was drawn randomly (uniformly) from P_i_ = [0.15, 0.26, 0.58, 0.95] and followed an exponential decay with ***τ*** = 0.19s (***P_t_*** = ***P_i_e***^−***τt***^), with P_t_ = 0 after 30 seconds. 10% of the Stochastic patches were programmed with the Deterministic patch return (“Stc:024 patches” with rewards at t = 0, 2, 4s) as reward-matched probe trials. The starting environment alternated each day, and the context transition in the middle of each session was only triggered between patches and included a 10 second blank (black) screen before initiating the second environment.

#### Muscimol inactivation experiments

To test the functional role of dorsal frontal cortex (dFC) activity in context-specific decision making, we inactivated dFC neurons using local infusion of the GABA_A_ receptor agonist muscimol (Sigma-Aldrich M1523), targeting the secondary motor cortex (M2, within the dFC). For the VR task, mice were trained as described above for two weeks in Phase 2. Then, once per day about 2 hours before experiments, they were briefly (∼15 minutes) anesthetized and injected with either muscimol (1ng/nL, 150 nL) or saline (150 nL) over 5 minutes targeting the right side M2 using the coordinates (Bregma +2.35mm AP, +1.0mm ML, -0.8mm DV from the skull surface). Mice were completely recovered from anesthesia before experiments and performed the task as usual. We injected muscimol or saline for 2 sessions each on alternating days. Mice showed no major deficits in general foraging behavior or task engagement (**Extended Data Fig. 13e,f**).

For dFC inactivation experiments in the FM task, mice were trained as described above through Phase 1 and Phase 2, and for four days in Phase 3. We then injected either muscimol or saline into M2 on alternating days and performed behavior experiments (FM task Phase 3) about 2 hours after injection. In the FM task, mice exhibited several behavioral changes under muscimol (**Extended Data Fig. 15**), including reduced context-specificity in decision behavior (**Fig. 6k-n**), but did not show any obvious deficits in motor behavior and still completed 10s-100s of trials per session.

#### Electrophysiological recordings and spike sorting

##### Acute recordings in the VR task

Spiking data were collected using Neuropixels 1.0 single shank or Neuropixels 2.0 4-shank probes with SpikeGLX (https://billkarsh.github.io/SpikeGLX/). Craniotomies were performed 12-24 hours before the recording using a dental drill and covered with Kwik-cast (World Precision Instruments), and mice recovered fully before experiments. On the day of recording, a 3D-printed shield was placed above the animal’s head to block light and sight of the experimenter during preparation. Prior to insertion, probes were dyed with DiI, DiD, or DiO (Thermofisher) to label probe tracts from each recording session (see **Extended Data Fig. 7** for several examples). Neuropixels probes were lowered using a Thorlabs micromanipulator (PT1-Z8) at 10-15 µm/sec. After reaching the target depth, the probe was allowed to settle in the tissue for 20 minutes prior to the start of the recording. Recordings typically lasted 50-60 minutes.

##### Chronic recordings in the FM task

Spiking data were collected using Neuropixels 1.0 single shank probes targeting the dorsal frontal cortex (+2.25mm AP, +0.8mm ML) and recorded with Bonsai software (OpenEphys.Onix1 software library) using the ONIX PCIe acquisition system (open Ephys). Craniotomies and probe implantation were performed at least one week before behavior and recording sessions began. Artificial dura (Dura-Gel, Cambridge Neurotech) was applied during probe implantation to protect the site of the craniotomy. Probes were mounted with dental cement using a custom adaptation of the Apollo implant system to enable probe recovery and were dyed using DiI to mark the tissue trajectory of the probe.^52^

For all Neuropixels recording experiments, spiking data were sorted using Kilosort 4 (https://github.com/MouseLand/Kilosort). This was followed by manual curation using Phy (https://github.com/cortex-lab/phy) to ensure waveform quality and recording stability for isolated units and to split doublet clusters. Though obtained, multi-unit and LFP signals were not used for any analysis in this study. See **Extended Data Fig. 8** for comprehensive recording summaries.

#### Histology and probe tract analysis

After the last recording session, mice were transcardially perfused with phosphate buffered saline (PBS) followed by 4% paraformaldehyde in PBS. The brains were sliced in 100 µm coronal sections using a vibratome (Leica). Brain sections were mounted on glass slides and stained with 4’, 6-diamidino-2-phenylindole (DAPI, Vectashield) in mounting medium. Slides were then imaged with a fluorescence microscope (Zeiss, Axio Scan.Z1). Probe tracts (labeled with fluorescence from DiI, DiD, and DiO) were identified in histology images (processed using Zen 3.3, Carl Zeiss Microscopy) and aligned using histological and electrophysiological landmarks to a reference atlas (Allen brain atlas CCF) using MATLAB software written by Dr. Andrew Peters (https://github.com/petersaj/AP_histology). Single units were sorted into 32 region and 7 macro-region categories using region identities from this reference atlas (see **Supplementary Table 1**). Single units localized in white-matter structures within 50µm (vertical) of an identified region were assigned to that region. Units localized in white matter structures outside of 50µm from other regions, or those assigned to regions outside the 32 categories, were not used for analysis.

#### Mouse and trial inclusion criteria

For experiments in the VR foraging task, 41 mice (22 males and 19 females) were trained and used for analysis, of which 10 were used for acute Neuropixels recordings. 47 recording sessions were used for neural analysis. For analysis of decision behavior, trials from individual sessions were only included if at least 10 trials were performed in that session (70548/70827, 99.61% of all trials). In the head-fixed VR task, mice occasionally disengaged from the task by pausing for brief periods of time, which occasionally occurred on a patch trial. Therefore, to prevent contamination of decision behavior by the disengaged state, we did not include trials with PRT (patch residence time) greater than 60 seconds for behavior analysis (2243/70827, 3.17% of all trials dropped).

For task-naïve recordings in the VR task, 6 mice (4 females, 2 males) were habituated to head-fixation as described above and then proceeded directly to Phase 2 of the foraging task for 3 days, during which acute Neuropixels recordings were performed. 17 recording sessions were used for neural analysis (Mouse 166 session 1 was not usable due to corruption of the raw data file).

For muscimol inactivation experiments in VR, mice were required to engage at least 10 trials of each type (Det and Stc:024) during both muscimol and saline sessions to be used for analysis. 10/14 mice trained on the task passed this inclusion criterion and were used for behavior analysis.

For basic behavior and recording experiments in the FM foraging task, 20 mice (9 males and 11 females) were trained and used for analysis, of which 4 were recorded from using chronic Neuropixels. 66 recording sessions (50 from Phase 2 and 16 from Phase 3) were used for neural analysis. All sessions, and all trials with PRT < 6 seconds, were used for behavior analysis.

For muscimol experiments in the FM task, mice were required to engage at least 10 trials of each type (Det and Stc:012) during both muscimol and saline sessions to be used for analysis. 13/13 mice trained on the task passed this inclusion criterion and were used for behavioral analysis.

#### Analysis of patch residence time (PRT) decision behavior

We analyzed decision behavior for patch foraging trials using the patch residence time (PRT) for different trial types. To evaluate whether mice made patch leaving decisions in a value-sensitive manner, we compared PRTs across four hidden trial types within Stochastic patches which had different initial reward probabilities (P_i_). P_i_ was not cued and was drawn randomly on trial initiation (see task descriptions above). In both the VR and FM tasks, PRTs on Stochastic patches scaled positively with Pi (**Extended Data Fig. 2a, 4d**), indicating sensitivity to local reward rates.^38^

We compared distributions of PRTs across patch types as a measure of context-dependency in foraging decision behavior, visualizing them using the cumulative density function (CDF) of P(Leave) over time (as in **Fig. 2d**). We quantified the divergence in PRT distributions using the Kolmogorov-Smirnov (KS) statistic, which measures the difference in CDFs at their point of maximum vertical difference. We compared the KS statistic across Deterministic and reward-matched Stc:012 or Stc:024 trial types, or across subsets of within-context trials (**Fig. 2h, Extended Data Fig. 4e**), and found that PRT distributions across contexts were more distinct than direct within-context comparisons. However, the KS statistic can be sensitive to large, local deviations in the shape of a distribution and can be skewed by small sample sizes. Thus, for some

analyses which required comparing distributions with different sample sizes (e.g. for muscimol experiments in VR, which contained relatively few trials), we controlled for this by down-sampling trials to equalize sample sizes and taking the average re-sampled KS statistic across 1000 draws. We validated our results by also measuring distribution differences using an alternative metric, the Wasserstein distance (see **Extended Data Fig. 2f, Extended Data Fig. 13g**), which in 1D is equivalent to the normalized absolute difference between CDFs (ΔCDF) and is less sensitive to local deviations. These measures were also well correlated in both tasks (**Extended Data Fig. 5e,j**), showing that they robustly capture differences in PRT distributions and decision behavior.

#### Analysis of patch leaving decisions using optimal foraging theory

To understand how mice might make patch leaving decisions for Stochastic or Deterministic reward dynamics, we simulated patch leaving behavior under a theoretically optimal decision policy based on the Marginal Value Theorem (MVT).^31,32^ A foundational result in optimal foraging theory, the MVT gives the optimal leave time for patch foraging (in the sense of long-term reward maximization) as the time when the marginal (in the limit instantaneous) reward rate equals the long-term future expectation (the global reward rate). Intuitively, the optimal time to leave on a given patch is the time when the marginal gain from staying an additional small time (i.e. **_δ_**_reward_ in **δ**t) is no longer worth the time opportunity cost compared to the long-term reward rate expected from leaving the patch (Δ_reward_ in Δt). To implement an MVT-based policy, a forager must therefore use estimates of both the local on-patch reward rate (**δ**_reward_/**δ**t) and the long-term global reward rate (Δ_reward_/Δt). To simulate MVT decision behavior for our task, we assumed that mice could estimate the global reward rate from their experience, which we computed as 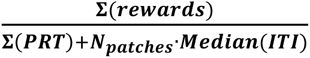 for all patch trials in a session. We used PRTs and the median ITI length to estimate the total time spent foraging, rather than the session duration, because mice often spent additional time doing non-foraging activities (e.g. exploration, grooming) during later times in the session. We took the average of this quantity across mice and sessions (0.43 rewards/second) to be an empirical measure of the global reward rate mice could potentially use, and we simulated leaving decisions (described below) against this as well as a range of surrounding values (**Extended Data Fig. 2c-e**, “emp” = empirical reward rate, 0.43 rewards/second).

To approximate an MVT policy, a forager must also have some time-varying estimate of the local reward rate on the current patch (**δ**_reward_/**δ**t). However, in our task design (and in many real-world scenarios), rewards are stochastic and sparse, and therefore local reward rates must be estimated based on ongoing experience and knowledge of reward statistics. If the forager has an estimate of the reward rate from experience or knowledge of reward dynamics, they could simply compare this expectation to Δ_reward_/Δt. Therefore, for simulations of MVT decisions on Deterministic patches, we assumed that mice are experienced and use a reward expectation (**δ**_reward_/**δ**t = 1 reward/second for 0 < t <= 2s, and **δ**_reward_/**δ**t = 0 for t > 2s) to guide their decisions. Simulating MVT leave times against the empirical estimate of Δ_reward_/Δt = 0.43 yielded a clear step-like increase in the P(Leave) CDF (**Extended Data Fig. 2d**) which was closely matched by the behavior of experienced mice on Deterministic patches (**Extended Data Fig. 1f,g**).

For Stochastic patches, which do not yield deterministic rewards, mice must instead rely on ongoing estimates of the local reward rate **δ**_reward_/**δ**t on each patch trial. Therefore, for simulations of MVT decisions on Stochastic patches, we assumed instead that mice could approximate **δ**_reward_/**δ**t using the local objective reward rate, which they directly experience. We computed this in a rolling 1-second window and compared it against Δ_reward_/Δt to generate simulated patch leaving decisions. In contrast to the Deterministic patch policy (above), this yielded a graded PRT distribution (**Extended Data Fig. 2c**), reflecting the decaying-probability reward dynamics and again matching the pattern of mouse behavior (**Extended Data Fig. 2b**).

To generate behavioral patterns for context-specific foraging strategies, we used these two policies (using reward expectations for Deterministic patches, and the local reward rate for Stochastic patches) to simulate PRTs on patches with rewards at t = 0, 1, 2, providing a direct comparison as between Deterministic and Stc:012 patches (**Fig. 2d,e**). This yielded PRT distributions that were distinct despite identical rewards and which, as constructed, we could understand as a consequence of the different time-varying reward expectations on Deterministic vs. Stochastic patches. Mouse behavior in both the FM (**Fig. 2d,e**) and VR tasks (**Fig. 3e,f**) matched this pattern, with greater P(Stay) early on Deterministic patches (reflecting a higher Deterministic reward expectation), and greater P(Leave) later on Deterministic patches (reflecting a lower reward expectation after the final deterministic reward, when the patch is depleted).

As optimality simulations such as this require many assumptions (e.g. mice have estimates of the global reward rate, can measure the local reward rate, and learn a time-varying reward expectation for Deterministic patches), we do not strongly conclude that mice use these precise strategies or follow an MVT computation. Rather, this analysis shows that, under the assumptions required to apply an MVT decision policy, different patterns of decision behavior could naturally emerge as a consequence of learned differences in Stochastic vs. Deterministic reward dynamics.

#### Integrator models of patch leaving decisions

Following our previous report (Bukwich et al.),^38^ we modeled patch leaving decisions in both the VR and FM foraging tasks as a drift-diffusion decision variable (DV) that integrates time on patch (***δ***) and reward receipt (***ρ***) toward a boundary (akin to evidence for or against the leaving decision).^35,38^ In the model (schematized in **Fig. 2i,j**), a decision variable **X_t_** is initialized to 0 at t = 0s (for each patch trial), incremented by a “drift rate” ***δ*** each time step, and is perturbed upon reward receipt by ***ρ*** (the “reward sensitivity”). At each time step, **X_t_** is converted into a leaving probability (**0 <= P_Leave_ <= 1**) using a sigmoid function centered at 0.5 with temperature ***T*** 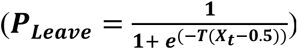. Thus, the model contains three parameters (***δ***, ***ρ***, ***T***), and different algorithms for the decision computation (“Models 1-4”, **Fig. 2j**) are produced by setting different constraints and definitions for ***δ*** and ***ρ***. Specifically, the “reward insensitive” Model 1 has (***δ* > 0, *ρ* = 0**); the “reward reset” Model 2 has (***δ* > 0, *ρ* = -X_t_**); the “reward integration” Model 3 has (***δ* > 0, *ρ* < 0**); and the “reward accumulation” model 4 has (***δ* > 0, *ρ* > 0**). Integrator models were fit by using empirical reward returns and mouse PRT behavior to generate DV rollouts, and then estimating the parameters (***δ***, ***ρ***, ***T***) using the fmincon function in MATLAB (model fitting toolbox) to maximize the log-likelihood between **P_Leave_** over time and the empirical patch leave times (PRTs). For parameter comparisons across patch types (**Fig. 2n, Extended Data Fig. 4i**), full models were used to estimate ***δ*** and ***ρ*** using trials with 0.5s <= PRT < 30s (for VR behavior) and 0.5s <= PRT < 6s (for FM behavior). For comparison and quantification of model performance, we fit models on 90% of the trials, computed the Bayesian Information Criterion (BIC) between predicted **P_Leave_** and ground truth leave times for held-out trials, and compared the average (across folds) cross-validated BIC for Models 1-4 (**Fig. 2k,l, Extended Data. Fig. 4f-h**).

We reported in our previous study (Bukwich et al.)^38^ that, in a similar VR patch foraging task, patch leaving decisions were best described using an integrator model (Model 3) that is scaled by a latent “patience” state **ϕ**, which captures individual variability in waiting behavior as well as within-session slow fluctuations in PRTs that are not explained by patch rewards. For behavior in the present VR task, we used a similar scaling parameter in the models to improve fits to behavior while preserving interpretability of the parameters ***δ*** and ***ρ***. For each patch trial, we computed the current value of a “patience” variable **ϕ** by taking the average of Gaussian-smoothed PRTs of the two adjacent trials (in either direction) and then scaled these to the average **ϕ** across all trials used for model fitting (***φ*** = ***φ***/***mean***(***φ***)).Then, to compute the DV **X_t_** in the model fitting procedure, we used ***δ*/ϕ** as the drift rate. As smaller drift rates yield longer PRTs, this captures the effect of the patience state on leaving decisions by scaling the drift rate to bias trial PRTs in the direction of those for neighboring trials. As we did not observe substantial mouse variability in the FM task, we did not include this “patience” scaling by **ϕ** for fits of the integrator models for FM behavior.

#### Generating predicted PRTs from model outputs

To simulate PRTs from specific DV rollouts fit for example mice (**Extended Data Fig. 2k,l**), we took the DVs **X_t_** generated from model fits, added gaussian noise (**σ = 0.1**), computed **P_Leave_** as described above, took the first timepoint where P^Leave^ > 0.5 as one simulated PRT, and repeated this procedure 100 times. We used this distribution of simulated PRTs to compute **P_Leave_** CDFs for different parameterizations of the DV (fit to behavior from Stochastic and Deterministic patch trials) and plotted these next to the empirical CDFs.

#### Normalization of single-neuron firing rate traces

For analysis of neural data, firing rate traces for each unit were generated by binning spikes into 50ms time bins and then z-scoring across the entire time series. Throughout the study, *σ* in plots of neural activity indicates 1 standard deviation (s.d.) of firing rate. For some analyses, local single-neuron activity was further normalized by subtracting activity within a pre-event (e.g. such as pre-patch trial) window and is therefore shown as “Δ activity” (still in units of *σ*, e.g. **Fig. 4d-e**).

#### Smoothing and de-noising for principal component projections

Where principal component analysis (PCA) was used to visualize low-dimensional projections of population activity, single-unit traces were further smoothed or averaged to reduce noise contributions to PC identification.^43^ For PC projections of activity across long timescales capturing the context transition (**Fig. 3k**, **Fig. 5g**), single-neuron traces were smoothed using a 10-second left moving-mean filter (i.e. only using previous, and not future, timepoints). For PC projections of patch trial activity (**Fig. 6h**), single-neuron activity was averaged within trial types to reduce noise from trial variability and then smoothed with a 1-second left moving-mean filter.

#### Analysis of single-neuron and population coding of environment context

We identified individual neurons as encoding environment context/block identity by measuring consistent changes in firing rate corresponding to environment identity (i.e. Stochastic vs. Deterministic) that occur at the time of context transition. For each neuron, we sampled activity in 60-second bins across the entire session and performed a two-sample t-test at α = 0.05 comparing the mean activity across samples from context 1 and 2. We then measured the change in mean activity “Δ activity” across 5-minute windows directly before and after the context transition and obtained a p-value (testing at α = 0.01) by comparison to a null distribution generated from sampling Δ activity around random within-context timepoints. Neurons that independently passed both statistical tests and exhibited the same direction of change (i.e. consistent global and transient changes) were considered to encode context identity **(Fig. 3l**, **Fig. 5h**). For regional comparisons of single-neuron coding (**Fig. 3l**), session/region samples with at least 5 neurons were used.

To quantify the separation of context activity states and rule out spurious contributions driven by slow activity fluctuations or drift artifacts (**Extended Data Fig. 9b, Extended Data Fig. 14a**), we split the single-neuron traces into rough quarter bins (for each session) corresponding to the first and second half of each context block, and then took the average activity of each neuron in each of these bins, forming population vectors for context 1 (half 1 and half 2) and for context 2 (half 1 and half 2). We then computed three angles comparing these vectors by first normalizing and then taking the inverse cosine of their dot product (***θ*** = ***cos***^−**1**^(***V*_1_** · ***V*_2_**)). For within-context comparisons, we computed the angle between the two halves within each context, and for across-context comparisons, we took the angle between vectors for the second half of context 1 and the first half of context 2. This directly compares the separation of activity states across contexts to within-session measures of activity change driven by variability unrelated to context identity.

#### Analysis of divergence between neural population activity subspaces

To analyze how context reconfigures population activity, we quantified the divergence of activity subspaces that separately capture neural activity for Deterministic and Stochastic contexts. To do this, we used canonical correlation analysis (CCA), which finds directions of maximum covariance between two sets of data and can be used to measure their subspace overlap (see **Extended Data Fig. 10** for graphical description, toy examples, and synthetic data control analyses).^44^ For each session or session/region sample, we first separately applied PCA to the epochs corresponding to the Stochastic and Deterministic contexts to define directions in the state space (the principal components) which capture within-context neural dynamics, and retained the top **N_PC_** components for each, where **N_PC_** is the number of PCs computed for the full dataset that capture 90% of total variance. These sets of PCs (**S_Stc_** and **S_Det_**) form orthonormal bases that define subspaces of the full state space.^25,43^ To measure their alignment, we first computed their cross-covariance matrix (***C*** = ***S_Stc_^T^S_Det_***) and then performed a singular value decomposition of C (***U*Σ*V^T^*** = ***C***). This yields two sets of singular vectors (or canonical directions) **U** and **V** which are successively maximally correlated with each other under the constraint that **U** and **V** are themselves orthogonal.^44,45^ The diagonal of ∑ contains the corresponding singular values (or canonical correlations) for each vector pair, which are related to their alignment and the angle between them. These angles (***θ***) can be used to quantify subspace alignment; completely overlapping subspaces yield all angles ***θ* = 0°**, while completely orthogonal subspaces yield all ***θ* = 90°** (see **Extended Data Fig. 10c-f** for illustrative examples in 3D with toy data). We measured overall subspace divergence as 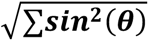, the “chordal distance” between the subspaces, as well as the number of singular vector pairs with ***θ* > 80°** (“orthogonal vectors”), which can be interpreted as the dimensional size of the components of the subspaces that are orthogonal to one another.

Alignment of neural subspaces has also been measured in studies of motor preparatory and movement activity by quantifying the amount of shared neural variance captured across subspaces defined during different trial epochs. To compare our geometric measure (the chordal distance, see above) with this variance-based measure, we computed the subspace alignment index introduced in Elsayed et al. 2016 for context-specific subspaces (**S_Stc_** and **S_Det_**) in our dataset. This alignment index quantifies the amount of neural variance in one epoch (e.g. context A) captured by projection into the subspace defined by the other epoch (e.g. context B), normalized by the maximum variance capturable with the same dimensionality (e.g. variance in A captured by subspace A). It therefore ranges from 0 to 1, where 0 indicates completely orthogonal subspaces (i.e. zero shared variance explained) and 1 indicates completely overlapping/aligned subspaces. We computed this metric for **S_Stc_** and **S_Det_**, subtracted it from 1, and multiplied by 100 to quantify the percentage (0-100%) of unexplained variance across subspaces. We found that this variance-based measure closely tracked the geometric chordal distance, using both full measurements and measurements normalized by subtraction of within-context comparisons (**Extended Data Fig. 10b**).

To evaluate whether subspace divergence was greater than expected from within-context neural variability (**Fig. 3n, Extended Data Fig. 10f**), we performed this analysis across epochs corresponding to the second half of context 1 and the first half of context 2 (across-context) and compared these by subtraction to the first and second halves of one context (within-context) as direct within-session controls. For comparison across regions, we measured subspace difference metrics for each brain region as the median of all region/session samples with at least 5 neurons.

Because we observed a strong correlation between subspace divergence and environment context/block coding across regions (**Fig. 3o**), we analyzed whether the alignment metrics were obligately correlated with linear firing rate changes along one dimension (e.g. along a “context coding” vector), as this could potentially yield a spurious association. To test this, we repeated the subspace alignment analysis using data from each session taken from two halves of one context and synthetically added activity with gaussian noise (µ = 0-2 s.d., σ = 0-0.1 s.d.) to different sized subpopulations of real neurons to replicate the presence of linear “context coding” at different levels. Although adding synthetic context coding to real activity slightly increased the chordal distance across contexts (**Extended Data Fig. 10i**), as expected from vector addition geometry, the relationship (β = 0.436, measured using regression) was weaker than that observed in the real data (β = 1.185). Importantly, adding synthetic context coding did not increase the number of orthogonal vectors between subspaces (**Extended Data Fig. 10j**), while real context coding was significantly correlated with the number of orthogonal vectors (**Extended Data Fig. 10h**).

#### Analysis of single-neuron decision variable (DV) coding

To identify single-neuron correlates of the foraging decision variable (DV), we used a similar approach as in our previous study (Bukwich et al).^38^ We first fit the cognitive DV to decision behavior for each mouse using the integrator model as described, extracted the DV for trials corresponding to each trial type (e.g. Deterministic trials or Stochastic trials), and concatenated these separately. For each neuron, we then extracted the (0.5s mean smoothed) firing rate corresponding to the same patch trials, concatenated these, and computed the correlation (***r***) between the unit activity and the DV, separately for Stochastic and Deterministic trials), which we took as a measure of DV coding strength. To test statistical significance, we repeated this process 1000 times, shuffling the trial order before concatenating, and computed a null distribution of ***r*** to test the observed value against. Neurons that exceeded either 0.5% tails (α = 0.01) of this distribution were considered “DV coding” for either the Stochastic or Deterministic DV. Neurons that had a significant correlation with either the Stochastic DV or the Deterministic DV, but not both, were context-specific DV coding cells (Stc DV and Det DV, respectively, as in **Fig. 4c**).

To measure general single-neuron DV coding across sampled brain regions, we computed, for each session/region sample with at least 5 units, the fraction of Stc DV cells and the fraction of Det DV cells, averaged these, and subtracted the chance value for that sample resulting from shuffling trial identities (**Fig. 4h**). Because we effectively applied two statistical tests (for Det DV and Stc DV coding), we used a Bonferroni-corrected (α/2) p-value cutoff for this analysis.

#### Analysis of reward responses and ramping activity in DV coding neurons

To assess whether DV coding neurons represented differently parameterized decision variables as predicted from fits of the integrator models, we measured local reward responses and ramping activity in neurons encoding the Stochastic DV (Stc DV) or Deterministic DV (Det DV). We included all neurons (pooled across all sampled regions) which passed our bootstrap statistical test at α = 0.01 for the Det or Stc DV with a positive correlation. We measured average activity in Stc DV and Det DV neurons around three trial events: the reward marking trial initiation (t = 0s, **Extended Data Fig. 11e**), additional patch rewards (at t > 0s, **Fig. 4f**, **Extended Data Fig. 11d**), and patch leave (the time the mouse moves off the patch location, **Fig. 4g**). For the trial initiation response, we included activity from all trials with PRT >= 1s, measured the reward response as the local change in mean activity across surrounding 0.5s windows, and measured ramping activity as the slope (β) for regression of activity against time in the interval [0.25 to 1s] after trial initiation (reported as change in activity/sec, or Δ*σ*·s^-1^). Patch reward responses were analyzed the same way, except centering the window to rewards that occur after t = 0s. For ramping activity before patch leave, we included trials with PRT >= 7s and no rewards after 4s so that activity was not contaminated by dynamics from recent reward responses. As this plot (**Fig. 4g**) is centered around the time the mouse moves off the patch, the time the animal makes the leaving decision and initiates movement is slightly before t = 0. We therefore measured ramping activity as the slope β for regression of activity against time in the interval [-2 to -0.5s], reported as Δ*σ*·s^-1^.

#### Analysis of population decision variable (DV) coding and DV coding vectors

To define DV coding vectors for measurements of population coding geometry, we concatenated Stochastic or Deterministic DVs as described above for the single-neuron analysis, extracted corresponding single-neuron activity for all neurons in the population (within sessions), and performed linear regression of the neuron traces (**X**) against the DV (i.e. ***DV*** = ***X^T^β***). The normalized coefficients **β/norm(β)** can be interpreted as a direction in the activity space that best captures how neural activity linearly covaries with the DV and thus defines a coding vector (or coding dimension) **β_Stc_** (for the Stochastic DV) or **β_Det_** (for the Deterministic DV).^11,43^ The angle between these coding vectors is the inverse cosine of their dot product (***θ*** = ***cos***^−**1**^(***β_Det_*** · ***β_Stc_***)). Within-context control comparisons of the angle between DV coding vectors were computed in the same way but instead using trial data corresponding to two subsampled halves of trial data within each context rather than across contexts. For both the VR and FM tasks, the DV coding angle within dFC (dorsal Frontal cortex) populations was significantly greater across contexts compared to direct within-context vector angles (**Fig. 4l**, **Fig. 6e**).

As some sessions included large populations of neurons, the concentration of measure for vector angles in high dimensions (concentrated around 90 degrees by chance) may potentially obscure the geometric analysis. Therefore, to rule out the possibility that we observe orthogonal DV coding vectors spuriously due to this dimensionality effect, we repeated the coding vector angle analysis in a low-dimensional (3D) space by first projecting the population activity onto 3 components defined by partial least squares (PLS) regression between the neural data and the model-fit DV response variable (**Extended Data Fig. 12d-f**, **Extended Data Fig. 14f,g**); we then computed the angle as described above. The 3D PLS space captured 80-100% of the explainable variance in the DV, indicating that it retains the majority of DV coding present in the population. For dFC populations in both VR (*θ* = 96.2°) and FM (*θ* = 77.6°) tasks, the coding vector angle was still close to 90 degrees and significantly greater than within-context comparisons.

#### Analysis of sequential trial dFC population trajectories within and across contexts

We analyzed trajectories of dFC population dynamics on patch trials in Phase 3 of the FM task to quantify and visualize the immediate effect of volitional context transitions on neural activity underlying foraging decisions (**Fig. 6f-i**). To measure trajectory distances (**Δ_Traj_**), we first projected dFC population activity into a reduced PC space capturing 90% of the total neural variance, and then for each sequential trial pair in each session (e.g. trial_n_ and trial_n+1_), we computed Δ_Traj_ between those trials’ trajectories (e.g. Traj_n_ and Traj_n+1_) as the sum of point-by-point Euclidean distances between them (∑***δ_Traj_***(***t***), where ***δ_Traj_***(***t***) = ***norm***(***Traj_n_***(***t***) − ***Traj_n_***_+**1**_(***t***)), as schematized in **Fig. 6f**). To visualize trial trajectories for sequential trials within or across contexts (**Fig. 6h**), we projected population activity onto the top 3 PCs and averaged non-overlapping trial trajectories within context (e.g. Det_n_ → Det_n+1_) or across contexts (e.g. Det_n_ → Stc_n+1_).

#### Population decoder analysis of decision variables

For linear decoding of DVs, we fit integrator DVs to mouse behavior (Model 3, as in **Fig. 2j**) and concatenated trial DVs and corresponding neural activity (for region/session populations) as described above for the coding vector analysis. However, instead of concatenating all trials, we performed leave-one-out cross-validation by iterating through trials, leaving each trial out once, computing the DV coding vector using the remaining data, projecting the neural data (**X**) for the held-out trial onto the coding vector using **DV_pred_ = X^T^β**, and computing the correlation ***r*** between the true DV and **DV_pred_**. For each session/region sample, we took the median ***r*** across all held-out trial predictions as a summary measure of decoder performance (cross-validated *r,* **Extended Data. Fig. 12a**, **Fig. 4l**, **Fig. 5d**, **Fig. 6d**). For cross-context decoder analyses, we projected trial data from one context onto the coding vector for the other context and computed cross-validated *r.* For shuffle comparisons, we randomly circularly permuted each neuron activity trace and then repeated the analysis as described. For DV decoders for the VR task, trials with 0.5s < PRT < 30s were used, and for the FM task, trials with 0.5s < PRT < 6s were used. Sessions were only used for the decoder analysis if they had at least 10 trials to train the decoder, and if the number of neurons (population size) for that region/session sample was at least 5.

#### Population decoder analysis of context identity

For linear binary decoding of context identity, we took the average activity for each patch trial (trial decoder) or each inter-trial-interval (ITI) period in the session as population vector samples, then randomly up-sampled the group (i.e. Stochastic or Deterministic trials/ITIs) with fewer samples to equalize sample sizes and remove bias. Iterating 10 times, we then held out 10% of the samples (non-overlapping across iterations), fit a Fisher’s Linear Discriminant (FLD) model on the remaining 90% to discriminate Stochastic vs. Deterministic identity (fitcdiscr in MATLAB), and tested predicted labels against ground truth, using the area under the receiver operating characteristic curve (auROC) as a performance metric. The average across auROCs for the 10 folds (cross-validated auROC) was used as an overall summary measure. For shuffle comparisons, class labels (Deterministic vs. Stochastic context identity) were randomly shuffled before model fitting.

### QUANTIFICATION AND STATISTICAL ANALYSIS

All data were analyzed with custom code written in MATLAB (version 2019b, Mathworks). All box and whisker plots represent the median, interquartile range, and range of the distribution unless otherwise specified. For statistical tests, normality of data and equal variance of groups were not assumed by default, and non-parametric tests were used for unpaired and paired group comparisons. All 95% confidence intervals are reported, and when necessary, were estimated by re-sampling. Tests for normality were used before applying any parametric tests (t-tests, ANOVA), and for all significance testing we used two-tailed tests at α = 0.05 or α = 0.01 (noted in figure legends). For statistical comparisons of PRT distributions, two-sample Kolmogorov-Smirnov tests were used. Resampling methods based on temporally permuted neuron traces or shuffled time bins were used to assess significance of single-neuron coding (described in detail above). Statistical significance of population decoder analyses was assessed by comparison to null distributions of performance statistics generated from shuffled trial sequences or class labels. The sizes of mouse groups were not pre-specified and approximated those of previous studies. Comprehensive details for all statistical tests are provided in **Supplementary Table 2**.

## ACKNOWLEDGEMENTS

We thank Mike Bukwich and Malcolm Campbell for help setting up the VR foraging task and discussions on experiment design; Edward Soucy and Yuwei Li of the Harvard Neurotechnology Core for engineering assistance; the Harvard Center for Biological Imaging for support with microscopy; Mitsuko Watabe-Uchida, Mark Burrell, Daniel Cardozo Pinto, and Adam Lowet for critical input and feedback on the manuscript; and Mark Churchland and Michael Shadlen, and their research groups, for fruitful discussion. This work was supported by grants from the National Institutes of Health (NIH) (5U19NS113201 to N.U. and 5R01DA05975 to N.U.) and the Simons Collaboration on Global Brain (to N.U.). L.K. was supported by a Helen Hay Whitney Fellowship.

## AUTHOR CONTRIBUTIONS

L.K. designed the foraging tasks, basing the virtual-reality (VR) foraging task in part on a previous design reported in Bukwich et al.^38^ L.K. built behavioral and recording setups. L.K. performed all behavior and recording experiments for the VR task, L.K. performed all behavior experiments for the free-moving (FM) task, and L.K. and J.S.S. performed chronic Neuropixels recordings for the FM task. L.K. performed muscimol inactivation experiments with assistance from G.Z. L.K. analyzed all behavioral and neural data. L.K. wrote the manuscript with feedback from N.U. No generative AI tools were used for any aspect of this work.

## DECLARATION OF INTERESTS

The authors declare no competing interests.

## DATA AND CODE AVAILABILITY

All code and data will be uploaded to a public repository upon publication

---

**Extended Data Figure 1.**
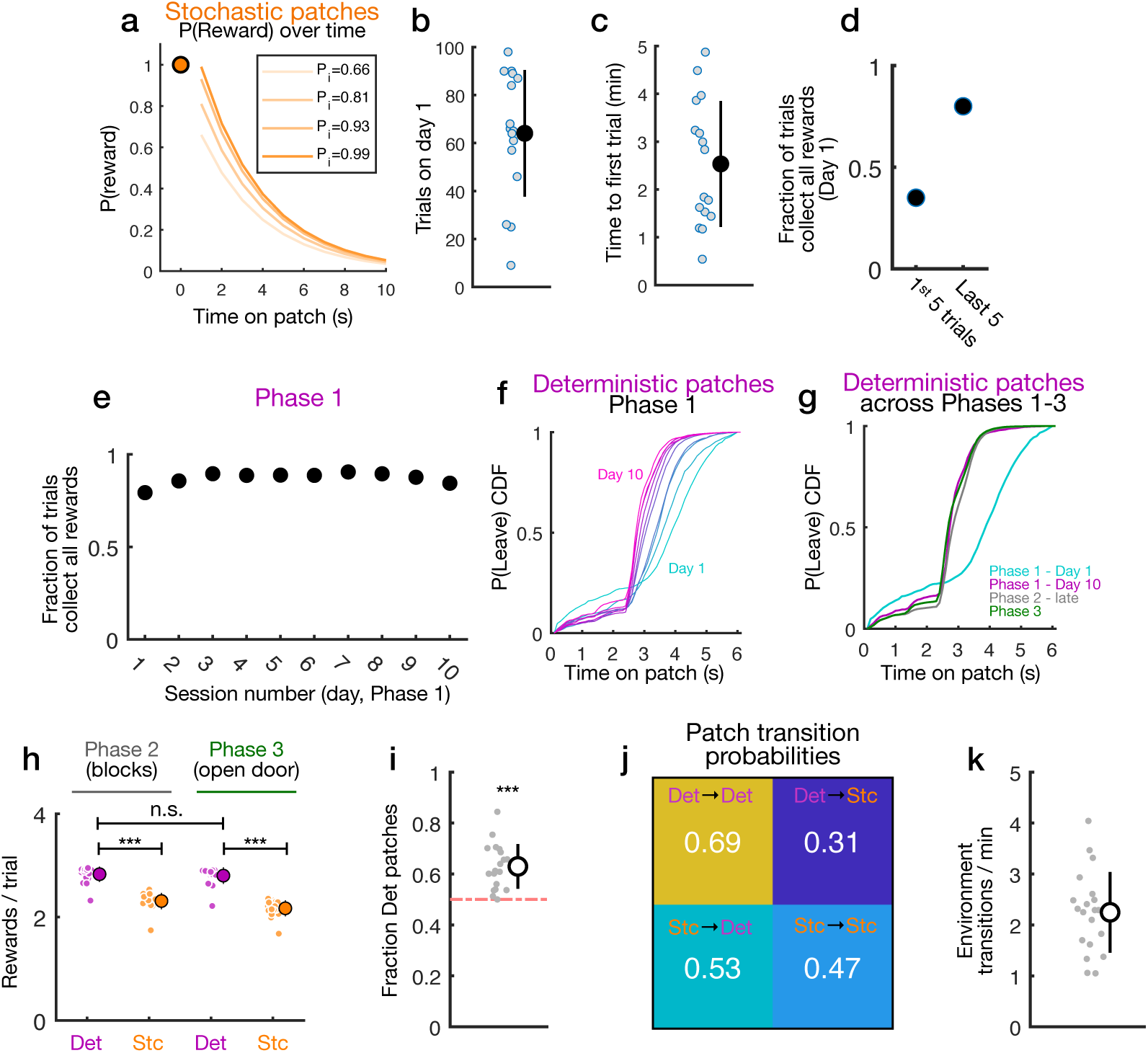
Additional analysis of behavior in two-context flexible foraging task. a) Reward probabilities over time for Stochastic patches. Each trial yields one 2µL water drop at t = 0s, and then one each second after with decaying probability (*ı* = 0.325s) starting with random P_i_ = 0.66, 0.81, 0.93, or 0.99 (see **Methods**). b) Number of patch trials performed by naive mice in the first session of Phase 1 of the foraging task (each dot is one mouse, mean ± s.d., *n* = 16 mice). c) Latency (minutes in the first session of the task) to the first patch trial (each dot is one mouse, mean ± s.d., *n* = 16 mice). d) Fraction of trials among the first 5 in the first session (left) and the last 5 in the first session (right) in which mice collected all three Deterministic rewards (data pooled across 16 mice). e) Fraction of trials in which mice collected all three Deterministic rewards for each session across the time-course of Phase 1 (days 1-10, data pooled across 16 mice). f) CDF of leaving probabilities P(Leave)_t_ for trials on each session (days 1-10) across Phase 1 (data pooled across mice, cyan = day 1, pink = day 10). g) CDF of leaving probabilities P(Leave)_t_ for Deterministic trials on day 1 in Phase 1 (cyan), day 10 in Phase 1 (pink), late (last 4 days) in Phase 2 (grey), and Phase 3 (green) showing similar PRT distributions and decision behavior across all phases of the task. h) Average number of rewards collected per trial for Deterministic and Stochastic patches in Phase 2 and Phase 3 (each dot is one mouse, mean ± s.d., SR test, *P* = 8.86e-5 [Phase 2 Stc vs. Det], *P* = 8.86e-5 [Phase 3 Stc vs. Det], *P* = 0.0522 [P2 Det vs. P3 Det], *n* = 20 mice). i) Choice probability for Deterministic vs. Stochastic patches in Phase 3 of the task when animals can volitionally switch environments (each dot is one mouse, mean ± s.d., t-test > 0.5, *P* = 2.4e-6, *n* = 20 mice). j) Transition matrix for Deterministic and Stochastic patches in Phase 3 of the task. k) Rate of volitional context transitions in Phase 3 (each dot is one mouse, mean ± s.d.). Standard deviation (s.d.); Wilcoxon signed rank test (SR test); \*\*\**P* < 0.001; \*\**P* < 0.01; \**P* < 0.05; n.s. p > 0.05.

**Extended Data Figure 2.**
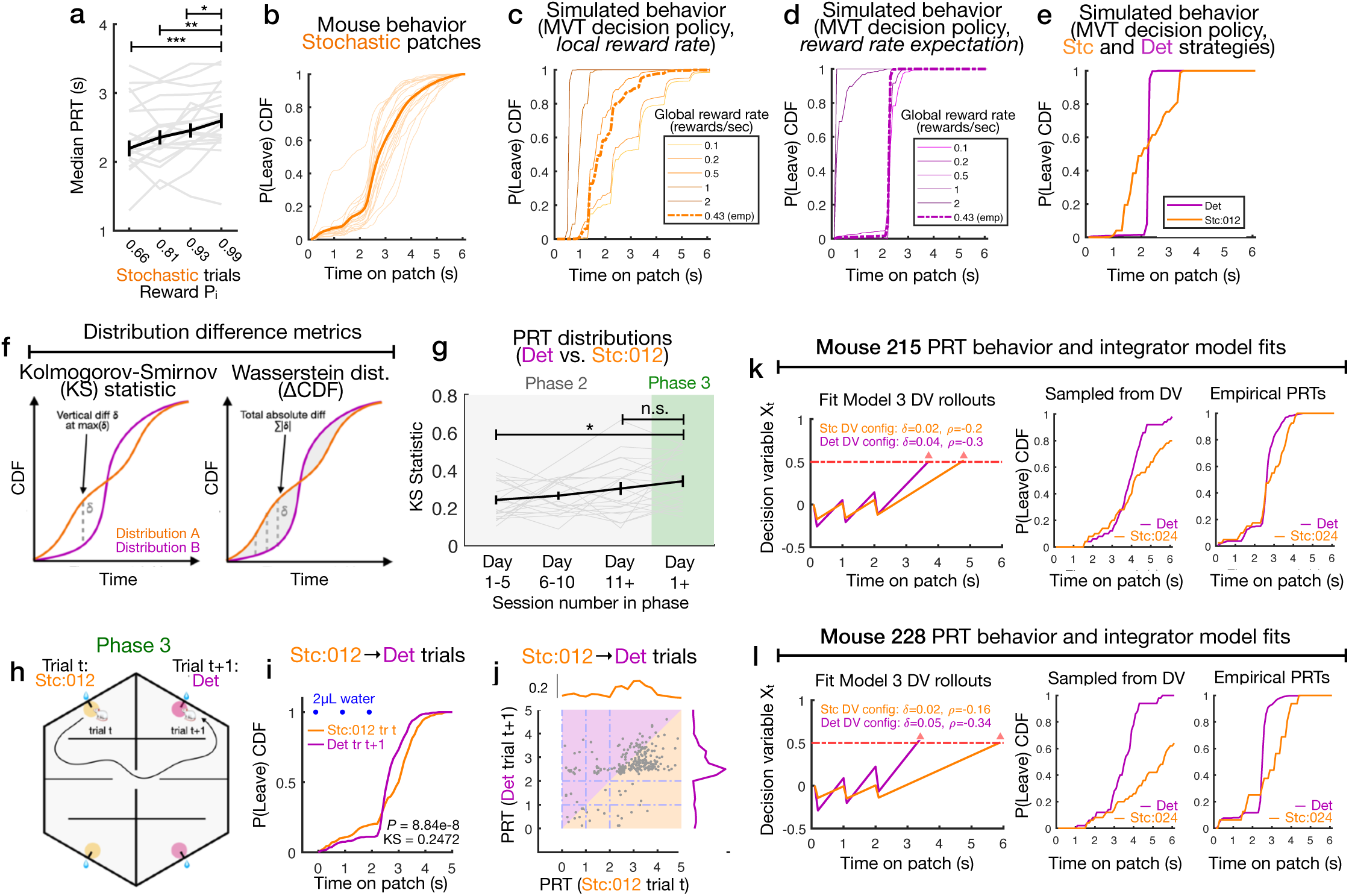
Additional analysis of decision behavior and context-dependency. a) Patch residence times (PRT, seconds) for hidden Stochastic trial types with different starting reward probabilities P_i_ = 0.66, 0.81, 0.93, or 0.99 (see also **Extended Data** Fig. 1a, median ± SEM; SR tests, *P* = 8.86e-5 [0.66 vs. 0.99], *P* = 0.004, [0.81 vs. 0.99], *P* = 0.0479 [0.93 vs. 0.99], *n* = 20 mice). b) CDF of leaving probability P(Leave)_t_ for Stochastic patches (excluding Stc:012 patches). Light orange lines are individual mice, and dark thick line is average across mice (*n* = 20 mice). c) Simulated leaving behavior for Stochastic patches with application of the Marginal Value Theorem (MVT, foragers compare local reward rate on the current patch with an estimate of the global reward rate and leave when these are equal, see **Methods**). PRT distributions were simulated against different global reward rates (shades of orange) and against an empirical estimate of the global reward rate (0.43 rewards/s, thick dashed line) from mouse performance (total rewards/time spent foraging). On Stochastic patches, experienced foragers are assumed to estimate the local reward rate as actual rewards/s over the last second on the patch. d) Simulated leaving behavior for Deterministic patches with application of the MVT against different global reward rates (shades of purple) and an empirical estimate of the global reward rate (0.43 rewards/s, thick dashed line). On Deterministic patches, experienced foragers are assumed to use learned reward rates (i.e. 1 reward/s for 0s < t <= 2s, and 0 reward/s for t > 2s). e) Patch residence time (PRT) distributions (CDF of leaving probability) for simulated leaving decisions on Deterministic and Stc:012 patches using decision policies reflecting strategies for Stochastic and Deterministic patches (**Extended Data** Fig. 2c**,d**, **Methods**). f) Diagram showing difference metrics for comparison of PRT distributions. The Kolmogorov-Smirnov (KS) statistic measures the maximum vertical distance δ between CDFs (left), and ΔCDF (equivalently the 1D Wasserstein metric) measures the total or average absolute difference between CDFs (right). g) KS statistic comparing Deterministic vs. Stc:012 PRT distributions across early (days 1-5), middle (days 6-10), and late (days 11+) Phase 2 progression and Phase 3 (mean ± SEM, SR tests, *P* = 0.0438 [early Phase 2 vs. Phase 3], *P* = 0 .3703 [late Phase 2 vs. Phase 3], *n* = 20 mice). h) Cartoon showing example of a Stc:012 (trial t) followed directly by a Deterministic (trial t+1) sequence in Phase 3 with volitional transitions. i) P(Leave) CDFs comparing PRT distributions for Stc:012→Deterministic trial sequences (KS test, *P* = 8.84e-8, *n* = 271 trials). j) Scatter plot showing PRTs for Stc:012→Deterministic trial sequences in Phase 3 and histograms of PRT distributions. k) Left, context-specific DV rollouts using parameters δ and ρ fit to integrator Model 3 for example Mouse 215. Right, simulated (using fit DV parameters, see **Methods**) and real PRT distributions as P(Leave) CDFs for Deterministic and Stc:012 patches. Context-specific configurations of the integrator model yield PRT distributions that closely match mouse behavior. l) The same as **Extended Data** Fig. 2k but for different example Mouse 228. Kolmogorov-Smirnov test (KS test); Wilcoxon signed-rank test (SR test); Standard deviation (s.d.); Standard error of mean (SEM); \*\*\**P* < 0.001; \*\**P* < 0.01; \**P* < 0.05; n.s. *P* > 0.05.

**Extended Data Figure 3.**
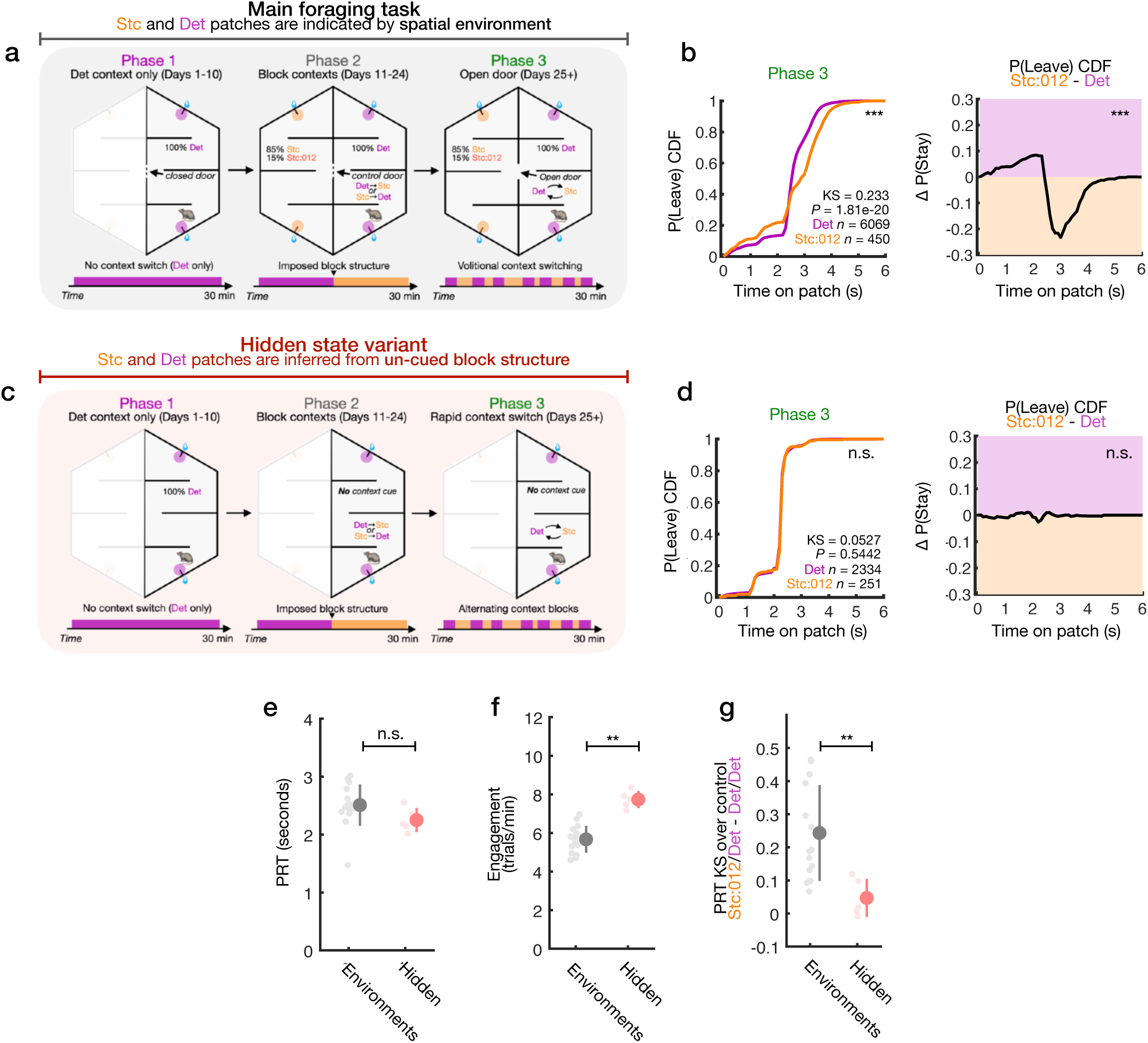
Alternative variant task design to test necessity of spatial context cue. a) The main foraging task design which uses spatial environments (separated right and left chambers) to indicate context of Deterministic or Stochastic patch reward dynamics (Fig. 1c). Training progresses from exposure to only the Deterministic context (Phase 1) through an imposed block structure (Phase 2) to volitional cortex switching (Phase 3, door left open). b) Left, CDFs of leaving probability P(Leave) for Deterministic patches (purple) and reward-matched patches in the Stochastic context (Stc:012, orange). The divergence of PRT distributions indicates context-dependency in the patch leaving decision process (KS test, *P* = 1.18e-20, *n* = 6069 [Det trials], *n* = 450 [Stc:012 trials]). Right, the difference in CDFs plotted separately on the left (Stc:012 - Deterministic), like in Fig. 2e. c) Alternate task design in which Deterministic vs. Stochastic context is a hidden state which is not cued, requiring mice to infer the current context from trial outcomes. Mice always forage from two ports in one spatial environment. d) Left, P(Leave) CDFs for Deterministic and Stc:012 patches (*n* = 5 mice) in the hidden state variant (KS test, *P* = 0.5442, *n* = 2334 [Det trials], *n* = 251 [Stc:012 trials]); right, the difference in CDFs, as in **Extended Data** Fig. 3b. e) Average patch residence time (PRT) across all trial types for the main spatial environment task (*n* = 16 mice) and the hidden state variant (*n* = 5 mice, mean ± s.d., U test, *P* = 0.0523). f) Average task engagement (patch trials / minute) for the main task (*n* = 16 mice) and the hidden state variants (*n* = 5 mice, mean ± s.d., U test, *P* = 0.0011). g) The divergence across Deterministic vs. Stc:012 PRT distributions quantified using the Kolmogorov-Smirnov (KS) statistic, normalized by within-Deterministic PRT variability (KS of sub-sampled Det trials), for the main task (*n* = 16 mice) and the hidden state variant (*n* = 5 mice, mean ± s.d., U test, *P* = 0.0015). Kolmogorov-Smirnov test (KS test); Mann-Whitney U test (U test); Standard error of mean (SEM); \*\*\**P* < 0.001; \*\**P* < 0.01; \**P* < 0.05; n.s. *P* > 0.05.

**Extended Data Figure 4.**
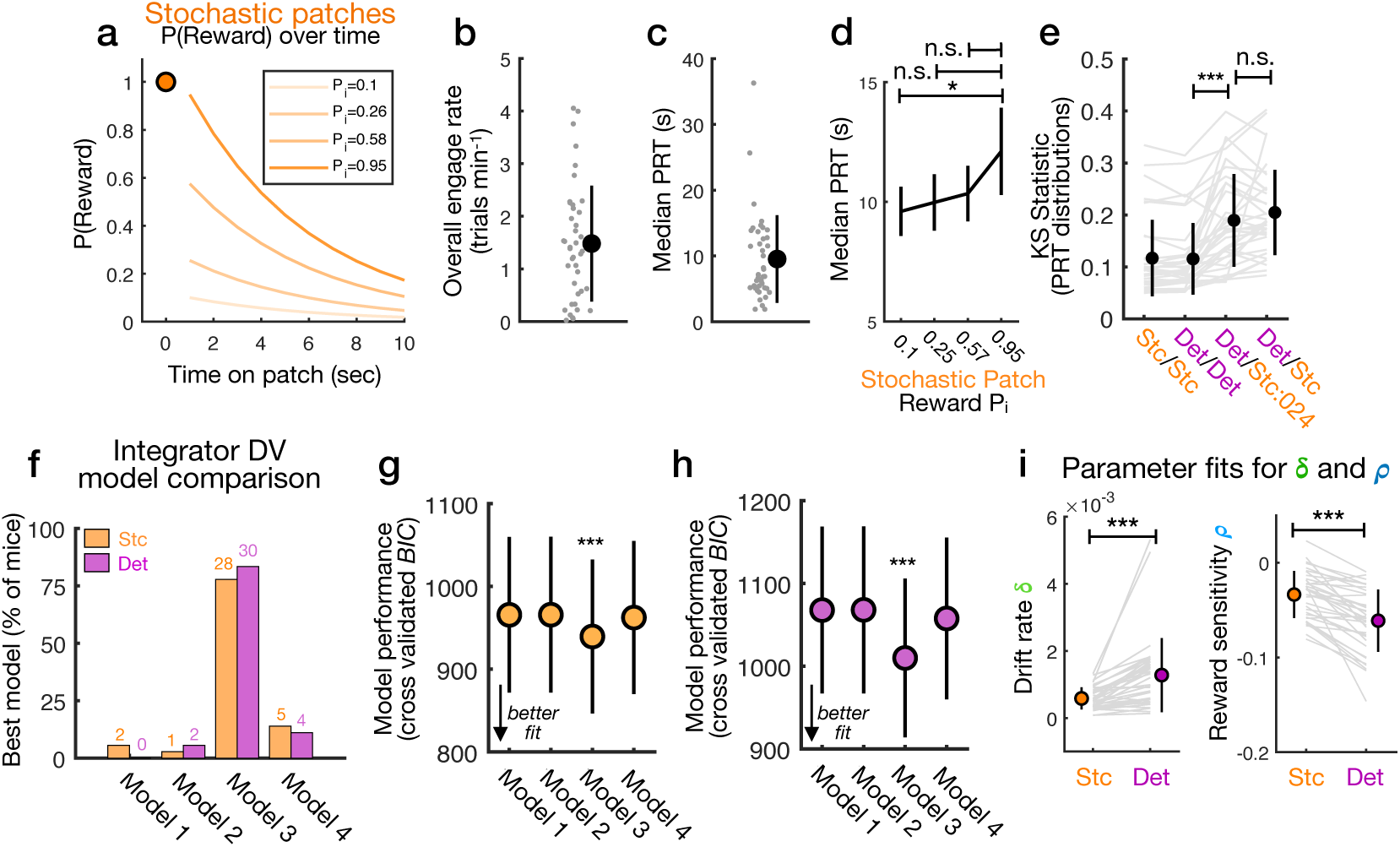
Additional details about virtual-reality (VR) foraging task and behavior. a) Reward probabilities over time for Stochastic patches. Each trial yields one reward at t = 0s, and then one each second after with decaying probability (*ı* = 0.19s) starting with random P_i_ = 0.15, 0.255, 0.575, or 0.947 (also see **Methods**). b) Patch engagement rate (trials per minute) for mice performing VR foraging task (each dot is one mouse, mean ± s.d., *n* = 40 mice). c) Patch residence time (PRT) for mice performing virtual foraging task (each dot is the median PRT for one mouse, mean ± SEM, *n* = 40 mice). d) PRTs for hidden Stochastic trial types in the VR task with different starting reward probabilities P_i_ = 0.1, 0.25, 0.57, or 0.95 (see also **Extended Data** Fig. 4a, mean ± SEM; SR tests, *P* = 0.0315 [0.1 vs. 0.95], *P* = 0.01746 [0.25 vs. 0.95], *P* = 0.0996 [0.57 vs. 0.95], *n =* 40 mice). e) Kolmogorov-Smirnov (KS) statistics for differences in PRT distributions across trial type comparisons (mean ± s.d., SR tests, *P* = 5.7e-7 [Det/Det vs. Det/Stc:024], *P* = 0.4372 [Det/ Stc:024 vs. Det/Stc], *n* = 36 mice). f) Model comparison analysis showing, for each integrator model (Models 1-4, Fig. 2j) and patch type (Stochastic or Deterministic), the fraction of mice for which that model was the best fit (quantified using the cross-validated BIC on held-out patch trials). The time/reward integrator Model 3 was also the best fit quantitatively (see **Extended Data** Fig. 4g**,h**). g) Performance (cross-validated Bayesian information criterion, BIC) of integrator models fit to mouse PRT behavior in the VR task and tested on held-out data for Stochastic patches (mean ± SEM, SR tests between models (*P* = 1.35e-5 [Model 1 vs. 3], *P* = 1.35e-5 [Model 2 vs. 3], *P* = 5.78e-5 [Model 3 vs. 4], *n* = 36 mice). h) Performance of integrator models on Deterministic patches (mean ± SEM, SR test between models (*P* = 2.07e-5 [Model 1 vs. 3], *P* = 1.92e-5 [Model 2 vs. 3], *P* = 5.05e-5 [Model 3 vs. 4], *n* = 36 mice). i) Left, comparison of Model 3 drift rate parameter *δ* fit to Stochastic vs. Deterministic trials (mean ± s.d., SR test, *P* = 5.85e-7, *n* = 36 mice). Right, comparison of reward sensitivity parameter *ρ* fit to Stochastic vs. Deterministic trials (mean ± s.d., SR test, *P* = 4.22e-7, *n* = 36 mice). Wilcoxon signed-rank test (SR test); Standard deviation (s.d.); Standard error of mean (SEM); \*\*\**P* < 0.001; \*\**P* < 0.01; \**P* < 0.05; n.s. *P* > 0.05.

**Extended Data Figure 5.**
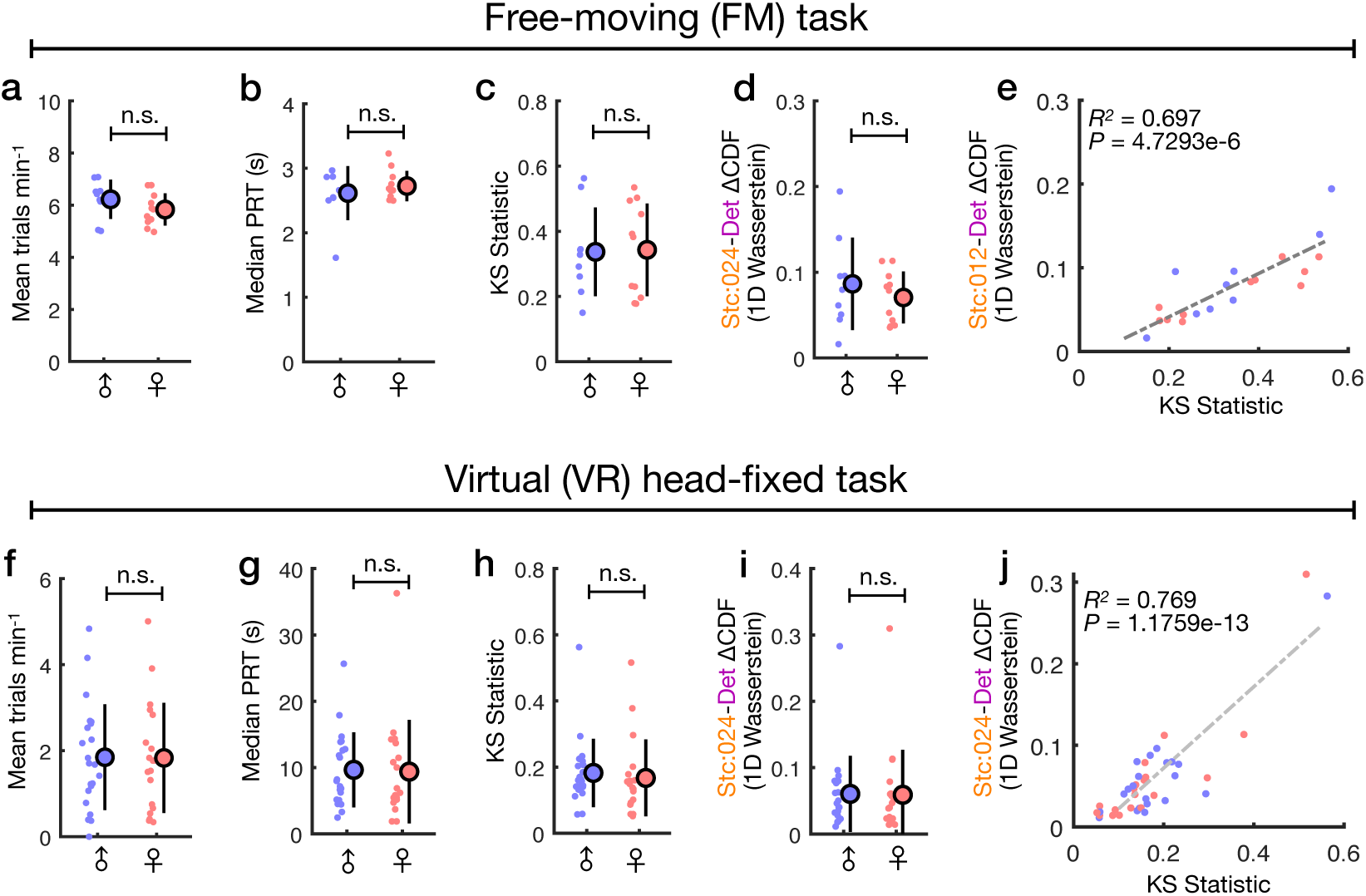
Lack of sex differences in general foraging behavior and context-dependent foraging decisions. a) Comparison of patch engagement rate (trials per minute) between male vs. female mice in the free-moving (FM) foraging task (mean ± s.d., U test, *P* = 0.1965, *n* = 9 males, *n* = 11 females). b) Comparison of median patch residence time (PRT, seconds) between male vs. female mice in the FM task (mean ± s.d., U test, *P* = 0.9697). c) Comparison of KS statistic for Stc:012 vs. Deterministic patch PRTs between male vs. female mice in the FM task (mean ± s.d., U test, *P* = 1). d) Comparison of ΔCDF for Stc:012 vs. Deterministic patch PRTs between male vs. female mice in the FM task (mean ± s.d., U test, *P* = 0.4941). e) Correlation between KS statistic and ΔCDF for all mice in the FM task (linear regression, R^2^ = 0.69, *P* = 4.73e-6, *n* = 20 mice). f) Comparison of patch engagement rate between male vs. female mice in the virtual (VR) foraging task (mean ± s.d., U test, *P* = 0.9296, *n* = 21 males, *n* = 19 females). g) Comparison of median patch residence time between male vs. female mice in the VR task (mean ± s.d., U test, *P* = 0.5513). h) Comparison of KS statistic for Stc:024 vs. Deterministic patch PRTs between male vs. female mice in the VR task (mean ± s.d., U test, *P* = 0.203). i) Comparison of ΔCDF for Stc:024 vs. Deterministic patch PRTs between male vs. female mice in the VR task (mean ± s.d., U test, *P* = 0.5157). j) Correlation between KS statistic and ΔCDF for all mice in the VR task (linear regression, R^2^ = 0.77; *P* = 1.18e-13, *n* = 40 mice). For all plots, each dot is one mouse (blue = males; red = females). Mann-Whitney U test (U test); \*\*\**P* < 0.001; \*\**P* < 0.01; \**P* < 0.05; n.s. p > 0.05.

**Extended Data Figure 6.**
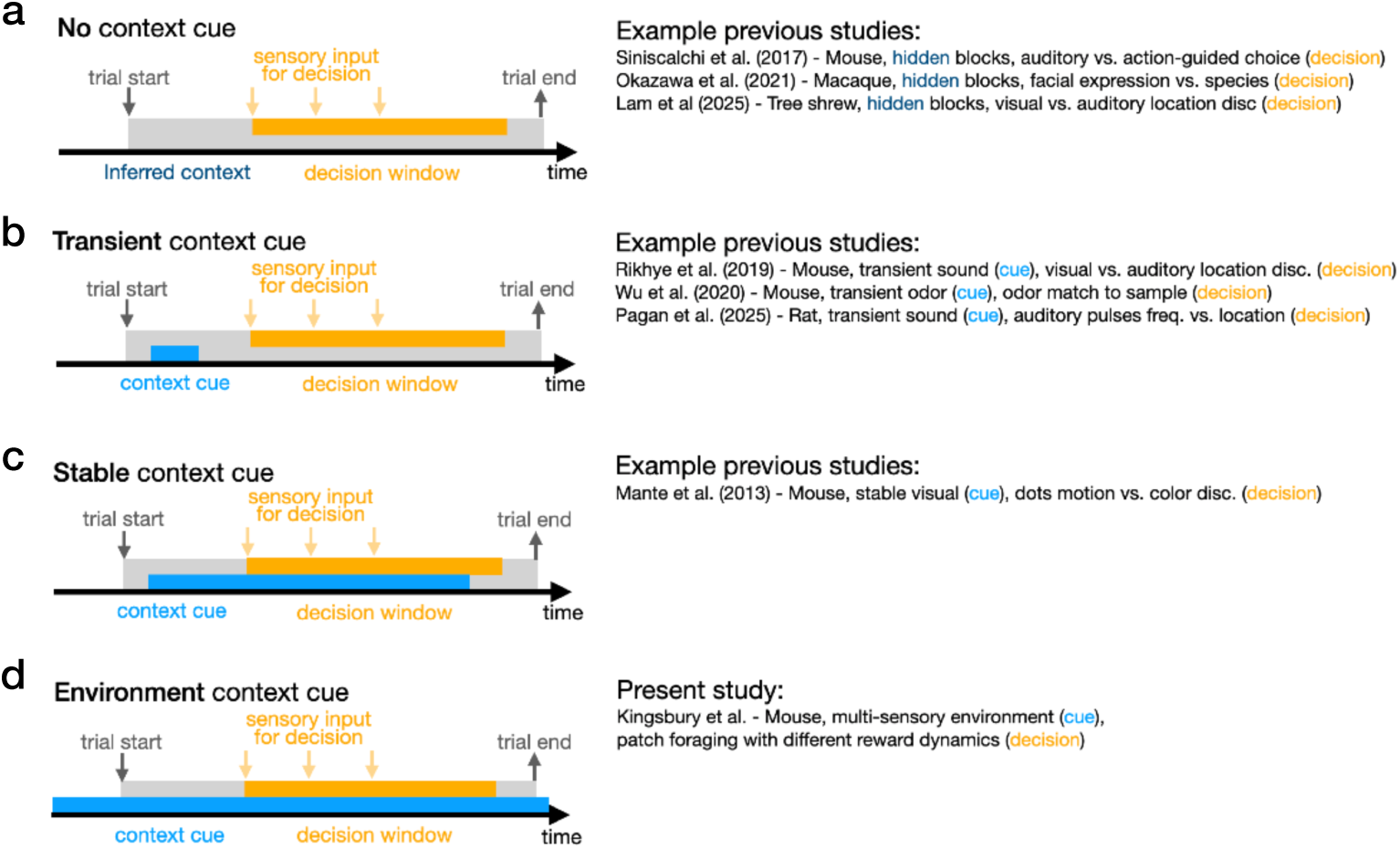
Comparison of different experimental designs and trial structures used to study context-dependent decision making. a) Tasks requiring animals to infer rule context for a decision process provide no explicit cue, and animals use error feedback to determine the correct context, typically presented in alternating blocks. Context identity is not correlated with external cues (see also **Extended Data** Fig 3c**-d**). b) Tasks providing transient sensory cues require internal maintenance of the context identity across a short (usually a few seconds) delay period before decision formation. Context identity is correlated with recent sensory history. c) Tasks providing a stable sensory cue indicate context unambiguously during and through the period of decision formation. Context identity is directly correlated with ongoing sensory input. d) Tasks using different environments (the main foraging task in this paper) indicate context continuously during decision trials and during inter-trial periods through visuospatial and other sensory information. Context identity is continuously correlated with sensory input.

**Extended Data Figure 7.**
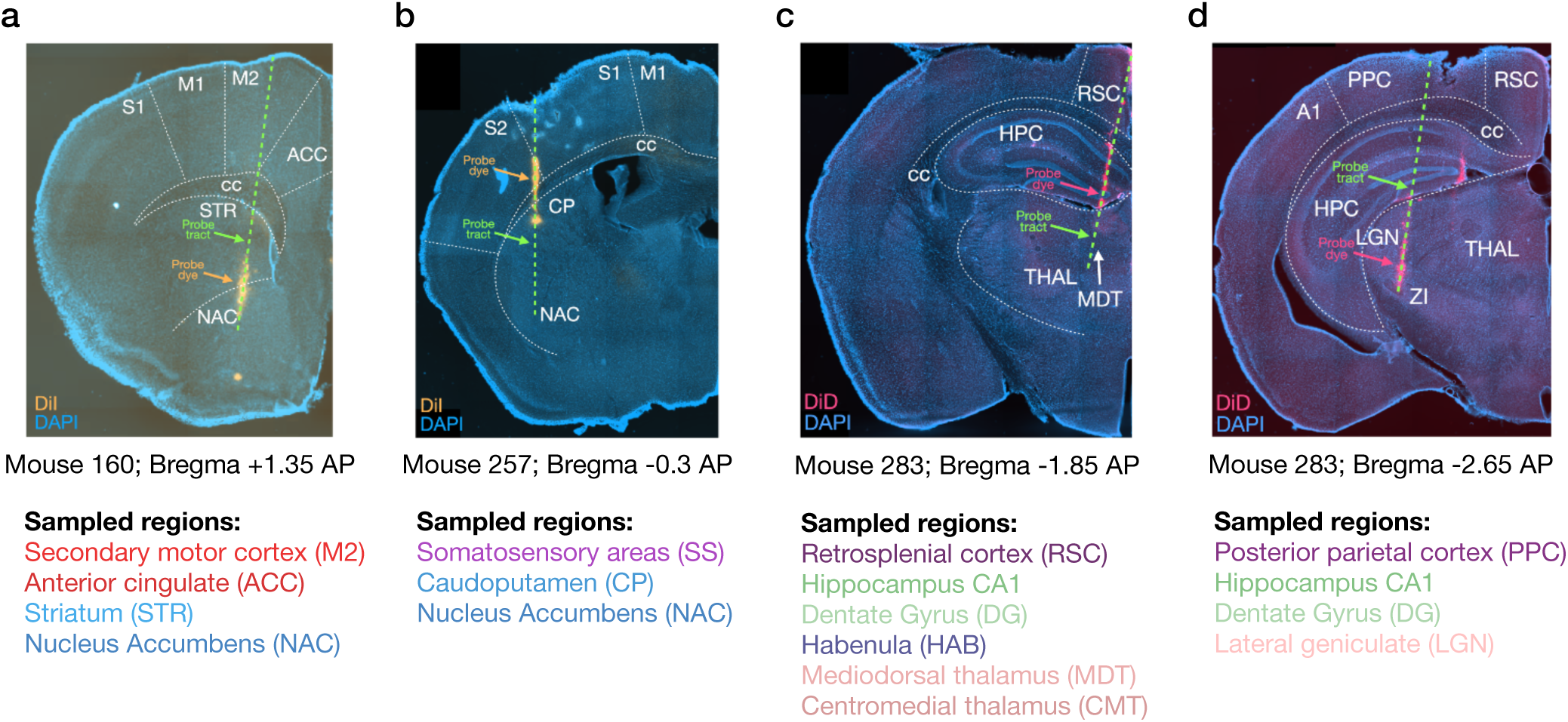
Examples of Neuropixels insertions targeting different brain regions. a) Example histology (Mouse 160) showing a 1-shank Neuropixels probe insertion targeting frontal cortex and basal ganglia regions (M2, ACC, STR, NAC). b) Example histology (Mouse 257) showing a probe insertion targeting sensory cortex and basal ganglia regions (SS, CP, NAC). c) Example histology (Mouse 283) showing a probe insertion targeting sensory cortex, hippocampal, subcortical and anterior thalamus regions (RSC, CA1, DG, HAB, MDT, CMT). d) Example histology (Mouse 283) showing a probe insertion targeting sensory cortex, hippocampal, and posterior thalamus regions (PPC, CA1, DG, LGN). For all images, blue = DAPI, orange/red = probe dye (DiI/DiD), green dashes = probe tract); insertion coordinates in millimeters (mm) from Bregma. See **Supplementary Table 1** for more detail on region identification.

**Extended Data Figure 8.**
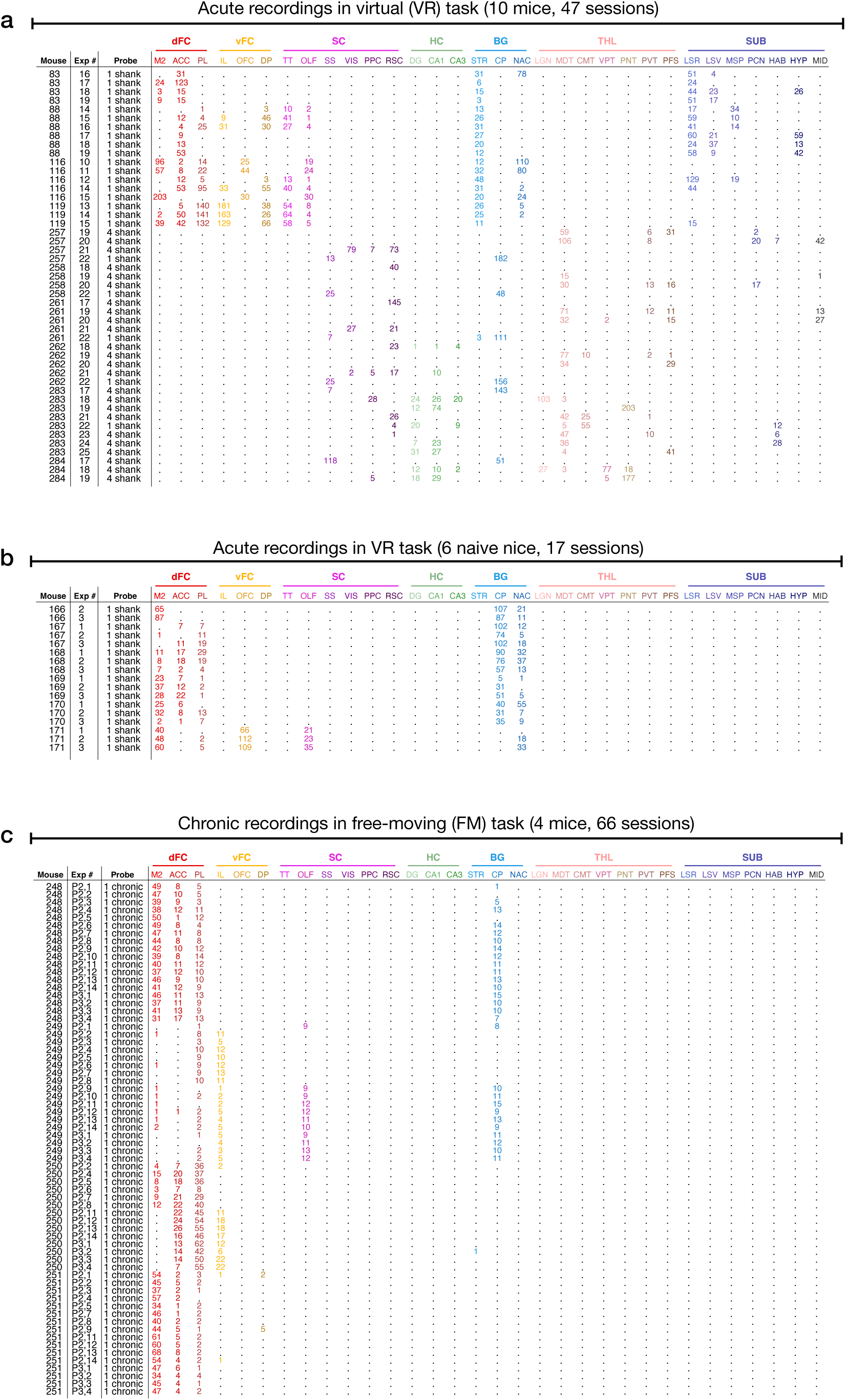
Additional information about Neuropixels recordings. a) Extended summary table for acute Neuropixels recordings in trained mice performing the virtual (VR) foraging task, showing mouse, session number, probe type (1-shank or 4-shanks), and number of recorded neurons from each region (*n* = 47 recording sessions from 10 mice). b) Extended summary table for acute Neuropixels recordings in task-naive (first 3 days) mice performing the VR task (*n* = 17 recording sessions from 6 mice). c) Extended summary table for chronic Neuropixels recordings in mice performing the free-moving (FM) foraging task across Phase 2 (14 days) and Phase 3 (4 days, *n* = 66 sessions from 4 mice).

**Extended Data Figure 9.**
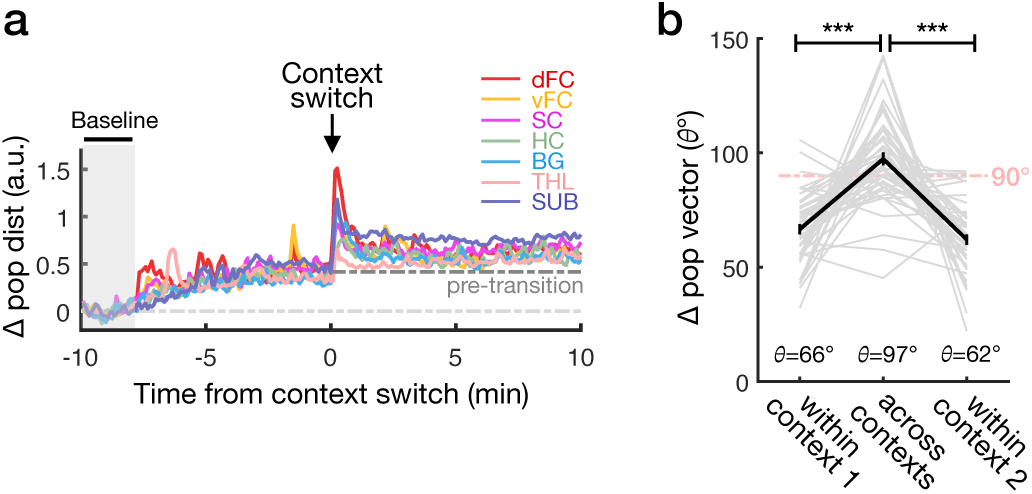
Analysis of neural activity states and dynamics around context transition. a) Time-course of population vector distances (Euclidean norm) from pre-transition time window (-10 to -8 min) across context transition showing transient dynamics and persistent activity shift. Pseudo-populations (across sessions and mice) of neurons were pooled within macro-regions. b) Angle measurements ($ in degrees) comparing similarity of across- vs. within-context population vectors (mean ± SEM, SR tests, *P* = 9.09e-8 [within context 1 vs. across], *P* = 3.13e-8 [across vs. within context 2], *n* = 47 sessions). Wilcoxon signed-rank test (SR test); Standard error of mean (SEM); \*\*\**P* < 0.001

**Extended Data Figure 10.**
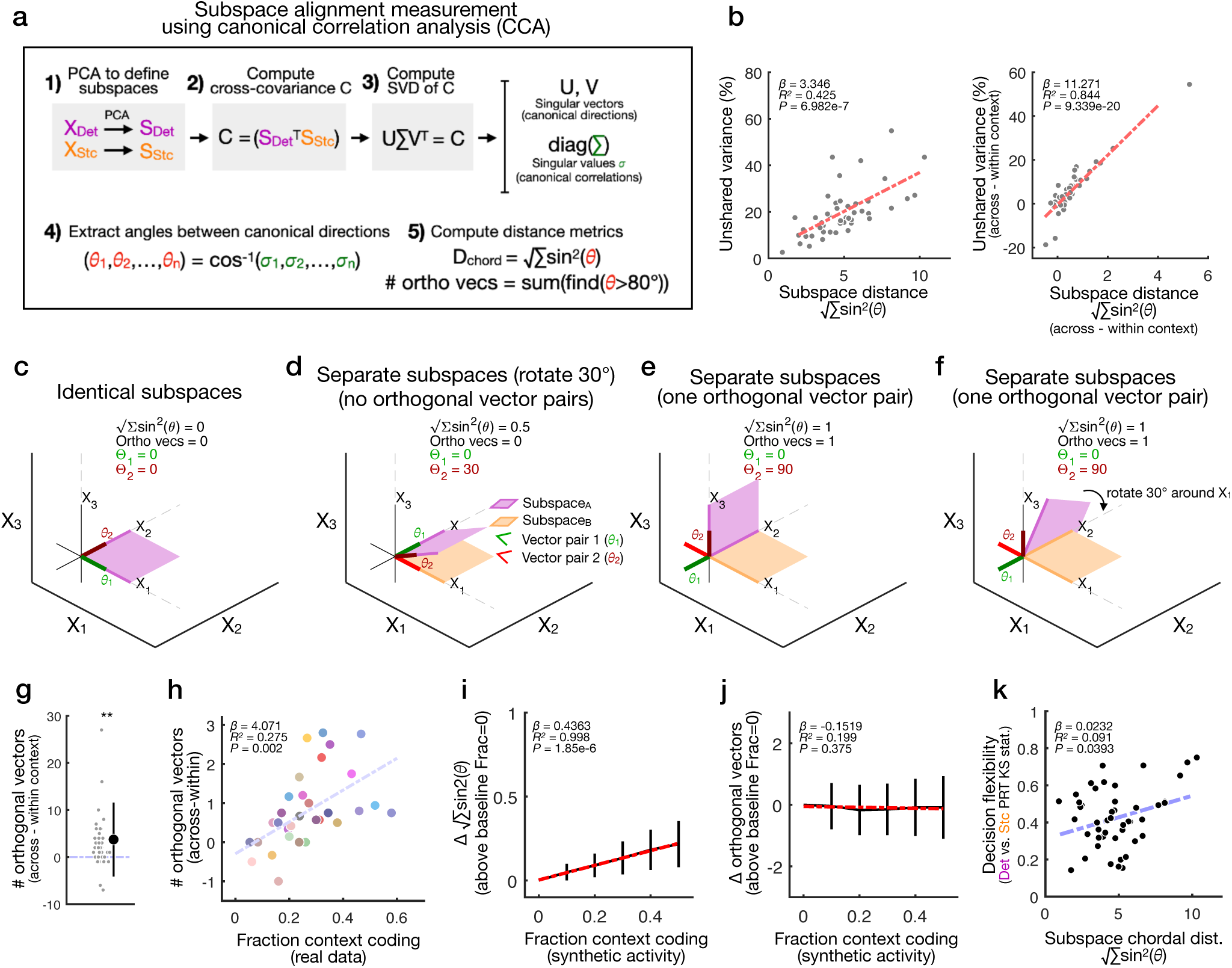
Description of subspace alignment analysis and additional controls. a) Graphical description of subspace alignment analysis pipeline. For each session, principal component analysis (PCA) is used on neuron spiking data within each context (Stochastic or Deterministic block) to determine subspaces **S_Det_** and **S_Stc_**, and canonical correlation analysis (CCA) is used to find canonical correlations and compute subspace distance metrics (the chordal distance and number of orthogonal vectors) between **S_Det_** and **S_Stc_** (see **Methods**). b) Left, correlation between the chordal distance metric √∑sin^2^(ϴ) and the amount of unexplained variance across subspaces. Unshared variance measures the percentage of neural variance captured by one subspace (e.g. **S_Det_**) that is not is not captured by the dimensions forming the other subspace (e.g. **S_Stc_**), equivalent (by complement) to the “subspace alignment index” introduced in Elsayed et al. 2016 and computed the same way. The chordal distance measures alignment of the dimensions forming the subspaces and closely tracks the amount of unexplained variance between them (linear regression, β = 3.35, R^2^ = 0.43, *P* = 6.98e-7). Right, the same relationship plotted after normalizing each measure by subtracting within-context values (linear regression, β = 11.27, R^2^ = 0.84, *P* = 9.34e-20). c) Example visualization of subspace alignment analysis applied to identical 2D subspaces (green = 1^st^ pair of canonical directions; red = 2^nd^ pair of canonical directions; ϴ = angle between vector pairs). d) Visualization of analysis applied to separate 2D subspaces with 30° angle between them and one shared canonical direction. e) Visualization of analysis applied to separate 2D subspaces with one orthogonal pair of canonical directions and one shared canonical direction. f) Visualization after rotating (30°) one subspace in **Extended Data Figure 10e**, maintaining the same subspace distance. g) Normalized (between - within context) number of orthogonal vectors across Deterministic and Stochastic context subspaces (mean ± s.d., t-test, *P* = 0.0021, *n* = 47 sessions). h) Correlation across regions between coding of environment context (fraction of neurons) and number of orthogonal vectors between context subspaces (R^2^ = 0.275, *P* = 0.002, *n* = 32 regions). i) Analytical control to test the effect of linear context coding on subspace distance measures. Relationship between chordal distance √∑sin^2^(ϴ) and addition of synthetic context coding activity (see **Methods**) in single neurons as observed empirically in Fig. 3o (mean ± s.d., fraction of synthetic context-coding neurons, linear regression, β = 0.44, R^2^ = 0.99, *P* = 1.85e-6). j) Relationship between orthogonal vectors and addition of synthetic context coding activity in single neurons (mean ± s.d., linear regression, β = -0.1519, R^2^ = 0.199, *P* = 0.375). k) Correlation across sessions between full chordal distance √∑sin^2^(ϴ) and decision flexibility, indexed as the KS statistic between Deterministic vs. Stochastic PRT distributions as in Fig. 3p (R^2^ = 0.091, *P* = 0.039, *n* = 47 sessions). Standard deviation (s.d.); \*\*\**P* < 0.001; \*\**P* < 0.01; \**P* < 0.05; n.s. *P* > 0.05.

**Extended Data Figure 11.**
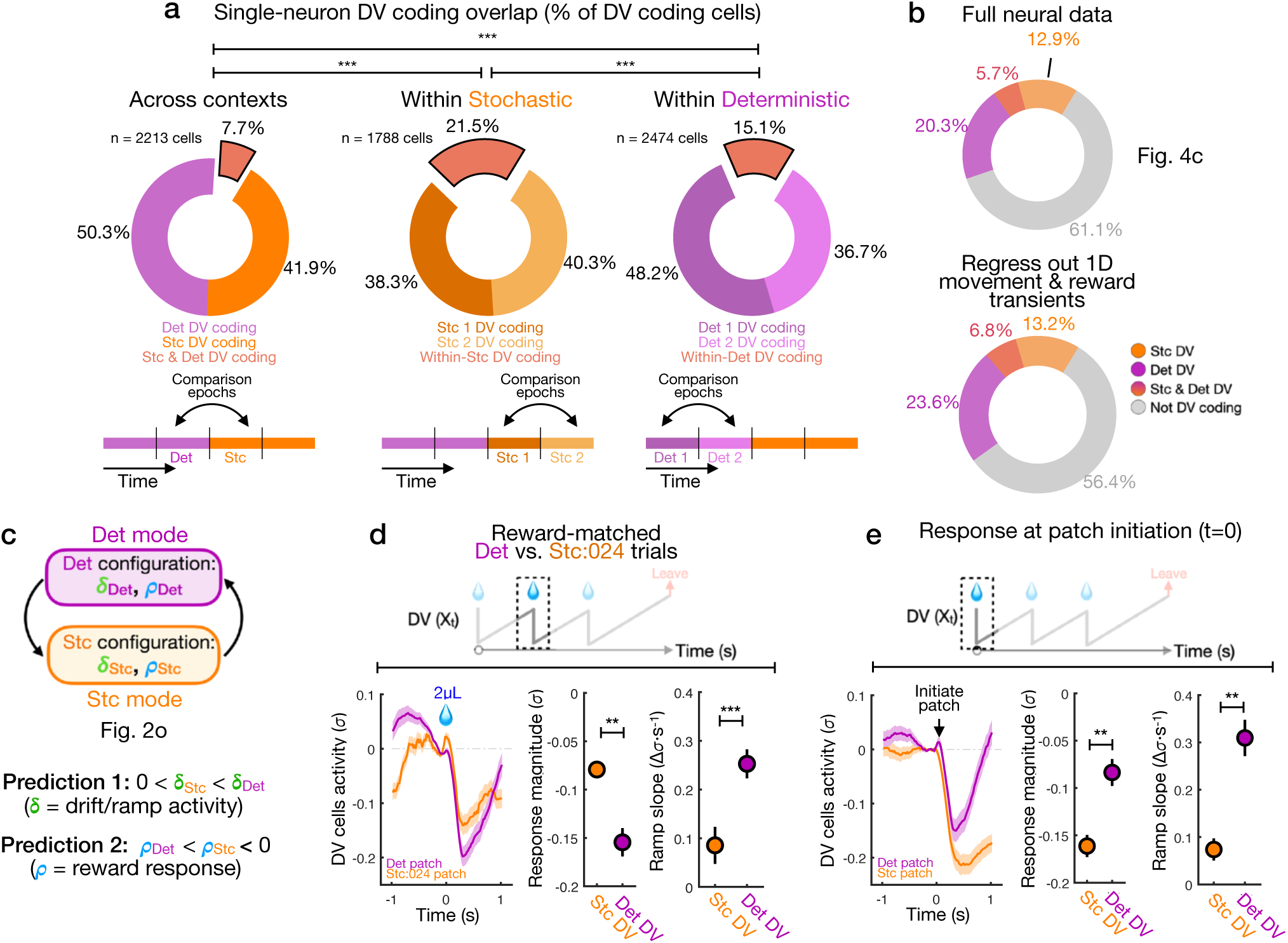
Additional analysis of context-specific decision coding in the VR task. a) Comparison of single-neuron DV coding and overlap across within-context control conditions. DV coding for individual neurons was computed using correlation analysis as in Fig. 4c (see **Methods**), except using even splits of within- or across-context trial data as schematized at bottom. Overlap of DV coding is quantified in pie charts. Across-context DV coding (left) overlaps significantly less than direct within-context comparisons (middle, right, Fisher’s exact tests, *P* = 1.1967e-27 [across vs. within-Stc], *P* = 1.842e-12 [across vs. within-Det], *P* = 8.079e-6 [within-Stc vs. within-Det]). b) Top, copy of Fig. 4c; bottom, breakdown of Stochastic (Stc) decision variable (DV), Deterministic (Det) DV, both DVs, and non-coding neurons from among all recorded (as in Fig. 4c) after regressing out linear correlations with movement (1D treadmill speed) and transient reward responses (0.5s window after each reward) for each neuron in the dataset (see **Methods**). c) Predictions of distinct activity dynamics for Stc DV and Det DV neural populations on patch trials based on parameter fits from integrator models (context-specific configurations of δ and ρ, Fig. 2n**,o** and **Extended Data** Fig. 4i). Cartoon illustration of DV predictions. d) Same as in Figure 4f but only using Stc:024 (reward-matched) patches. Average responses of Stc DV (orange, *n* = 387) and Det DV (*n* = 708, purple) cells around trial rewards (mean ± s.d.); middle, comparison of response magnitude (t-test, *P* = 1.1e-3); right, comparison of ramping slope following response (t-test, *P* = 7.14e-4). e) Left, average responses of Stc DV (orange, *n* = 575) and Det DV (*n* = 708, purple) cells at patch initiation (mean ± s.d.); middle, comparison of response magnitude (t-test, *P* = 4.08e-5); right, comparison of ramping slope following response (t-test, *P* = 6.87e-7). Standard deviation (s.d.); Standard error of the mean (SEM); Wilcoxon signed rank test (SR test); \*\*\**P* < 0.001; \*\**P* < 0.01; \**P* < 0.05; n.s. *P* > 0.05.

**Extended Data Figure 12.**
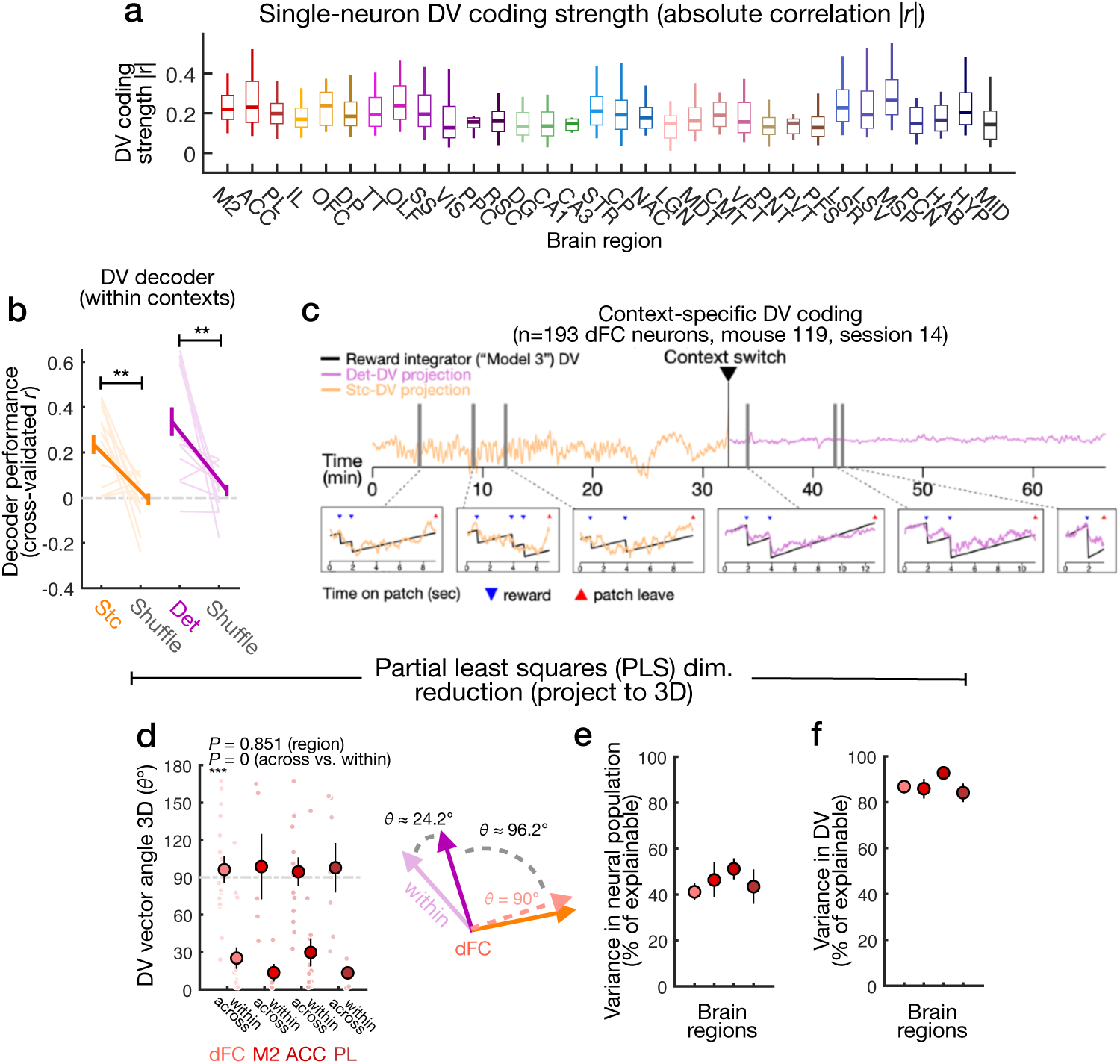
Population coding and decoding of context-specific decision variables. a) Breakdown across regions of single-neuron decision variable (DV) coding strength (median IQR and 5^th^-95^th^ percentile, absolute correlation |*r|*). b) Performance of **l**inear decoders for Stochastic or Deterministic DVs (cross-validated correlation *r* between predicted and model-fit DVs on held out trials) from dFC activity (mean ± SEM, SR tests, *P* = 0.0085, *n* = 14 sessions [Stochastic], *P* = 0.0017 [Deterministic], *n* = 13 sessions). c) dFC activity projected onto Det DV or Stc DV coding vectors (defined by linear regression coefficients, see **Methods**) from an example session (Mouse 116, session 10), showing population level DV coding in the same session both on Stochastic and Deterministic trials. d) Angle (ϴ degrees) as in Fig. 4k between Det DV and Stc DV coding vectors after projection into 3D space defined by partial least squares regression (PLS) components, and comparisons with within-context controls, for dFC neurons and each dFC region separately (mean ± SEM, t-tests against ϴ = 90°, *P* = 0.5662 [dFC], *P* = 0.7579 [M2], *P* = 0.7023 [ACC], *P* = 0.7103 [PL]; 2-way anova, *P* = 0 (within vs. between), *P* = 0.8516, region, *n* = 50 session samples). e) Variance explained (% of total, mean ± SEM) in the neural population data (spanning concatenated patch trials) by the top 3 PLS components used for the analysis in **Extended Data** Fig. 12d. f) Variance explained (% of total explainable, mean ± SEM) in the DV response variable by the top 3 PLS components used for the analysis in **Extended Data** Fig. 12d. Standard deviation (s.d.); Standard error of the mean (SEM); Wilcoxon signed rank test (SR test); \*\*\**P* < 0.001; \*\**P* < 0.01; \**P* < 0.05; n.s. *P* > 0.05.

**Extended Data Figure 13.**
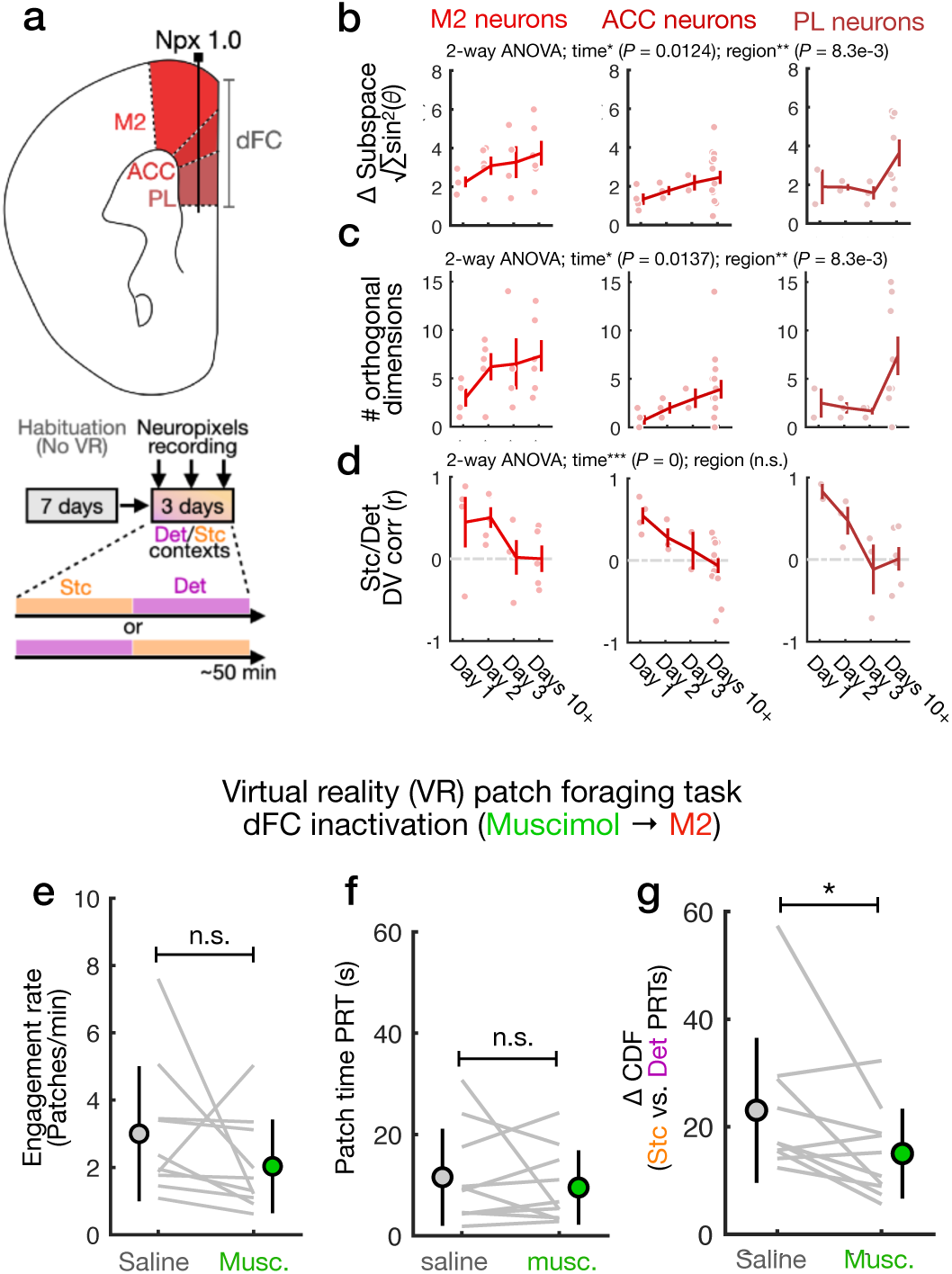
Neural recordings in task-naive mice and additional analysis of behavior under dFC inactivation. a) Schematic of experiment to measure neural activity in task-naive mice; After habituation to running wheel and head-fixation (see **Methods**), acute Neuropixels recordings targeting dFC were performed on the first 3 days of Phase 2 of the VR foraging task (Phase 1 was omitted). b) Subspace divergence (chordal distance √∑sin^2^(ϴ)) in dFC neurons from task-naive (days 1-3) and experienced sessions (mean ± SEM, 2-way anova, *P* = 0.0083, region, *P* = 0.0124, time, *d.f.* = 57). c) Number of orthogonal vectors (ϴ > 80°) in dFC neurons from task-naive (days 1-3) and experienced sessions (mean ± SEM, 2-way anova, P = 0.0083, region, *P* = 0.0137, time, *d.f.* = 57). d) DV coding similarity (correlation between Stc DV and Det DV coding strength *r*) across days (mean ± SEM, 2-way anova, *P* = 0.8174, region, *P* = 0, time, *d.f.* = 53). e) Comparison of patch engagement (trials/min) for mice in the virtual (VR) foraging task under Muscimol injection in M2 vs. saline (mean ± s.d., SR test, *P* = 0.0645, *n* = 10 mice). f) Comparison of overall patch residence time (median PRT for each mouse) behavior under Muscimol vs. saline control sessions (mean ± s.d., SR test, *P* = 0.9219, *n* = 10 mice). g) Comparison of ΔCDF (the 1D Wasserstein metric) of Deterministic vs. Stochastic PRT distributions for mice in the VR task under Muscimol injection in M2 vs. saline (mean ± s.d., SR test, *P* = 0.0273, *n* = 10 mice, alternative to KS statistic, see **Extended Data** Fig. 2f). Standard deviation (s.d.); Standard error of the mean (SEM); Wilcoxon signed rank test (SR test); \*\*\**P* < 0.001; \*\**P* < 0.01; \**P* < 0.05; n.s. *P* > 0.05.

**Extended Data Figure 14.**
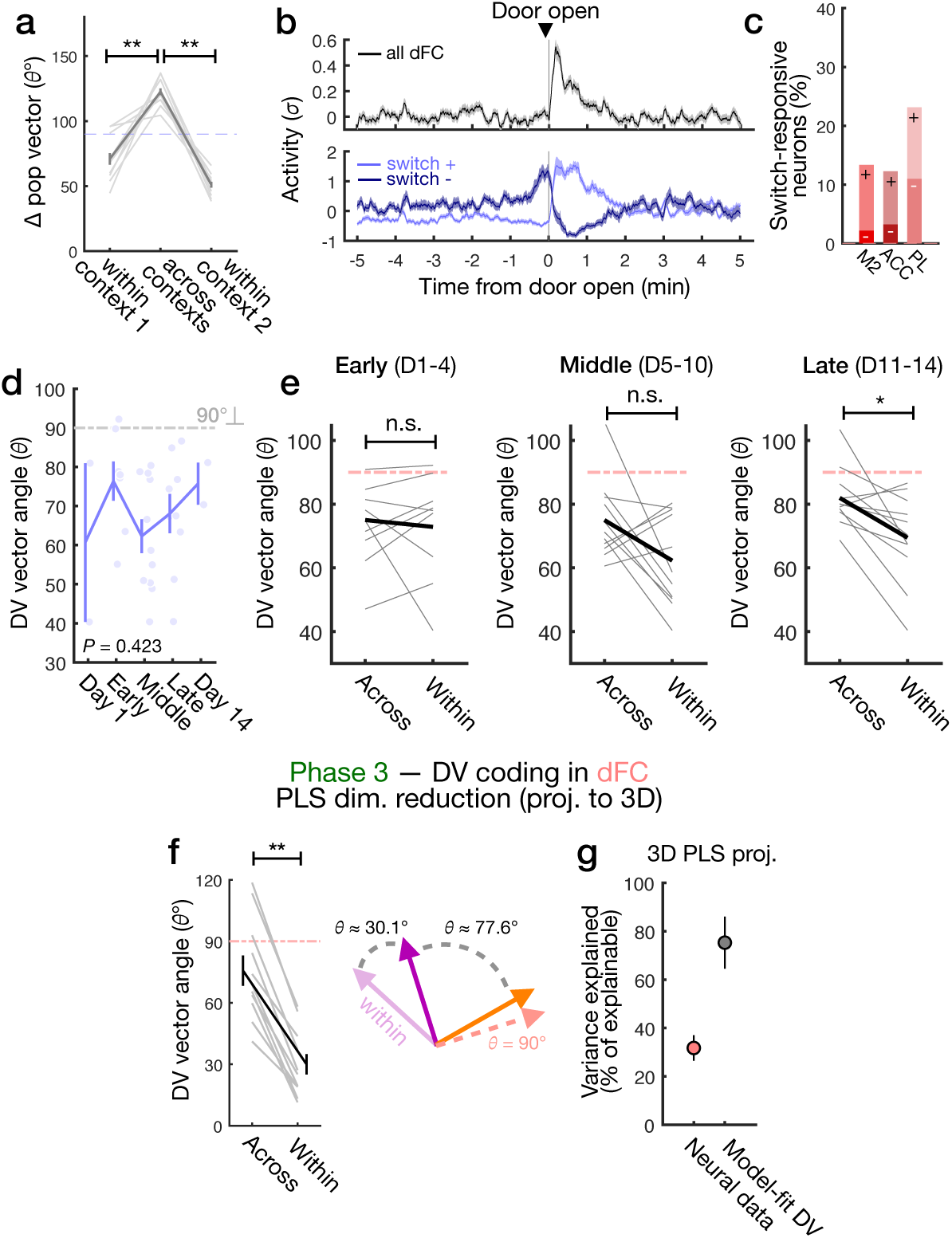
Additional analysis of chronic dFC recordings in free-moving (FM) task a) Angles (ϴ°) comparing within- vs. between-context dFC population vectors (mean ± SEM, SR tests, *P* = 0.002 [within context 1 vs. across contexts], *P* = 0.002 [within context 2 vs. across], *n* = 10 sessions). b) Top, transient response in mean population activity of dFC neurons centered around the door open time (mean ± s.d.); bottom, averages of subpopulations that are excited (light blue, *n* = 93) or inhibited (dark blue, *n* = 41) transiently at context transition (“switch-responsive” neurons, permutation tests at α = 0.01). c) Breakdown of switch-responsive neurons across dFC regions (M2, ACC, PL, light, positive; dark, negative responses). d) Angle (ϴ°) between within-context DV coding vectors (measured from even splits of patch trials) in dFC populations across days during Phase 2 progression (mean ± s.d., linear regression against time bins, *P* = 0.423). Within-context ϴ = 67.95° ± 14.93 (mean ± s.d., plotted as light blue line in Fig. 5q for comparison). e) Paired comparisons of across-context vs. within-context DV vector angles for different time-points in Phase 2 (black line = mean, SR tests, *P* = 0.6523, *n* = 9 sessions [early days 11-14]; *P* = 0.1016, *n* = 11 sessions [middle days 5-10]; *P* = 0.0137, *n* = 11 sessions [late days 11-14]). f) Angle (ϴ degrees) as in Fig. 6e between Det DV and Stc DV dFC coding vectors in Phase 3 of the FM task after projection into 3D space defined by partial least squares (PLS) components (mean ± SEM, SR test, *P* = 4.8828e-4, *n* = 12 sessions); illustration at right of within-context and across-context DV coding vector geometry. g) Variance explained (% of total, mean ± SEM) in the neural population data (spanning concatenated patch trials) and the model-fit DV (% of total explainable, mean ± SEM) by the top 3 PLS components used for the analysis in **Extended Data** Fig. 14f. Standard deviation (s.d.); Standard error of the mean (SEM); Wilcoxon signed rank test (SR test); \*\*\**P* < 0.001; \*\**P* < 0.01; \**P* < 0.05; n.s. *P* > 0.05. Each line or dot is one session.

**Extended Data Figure 15.**
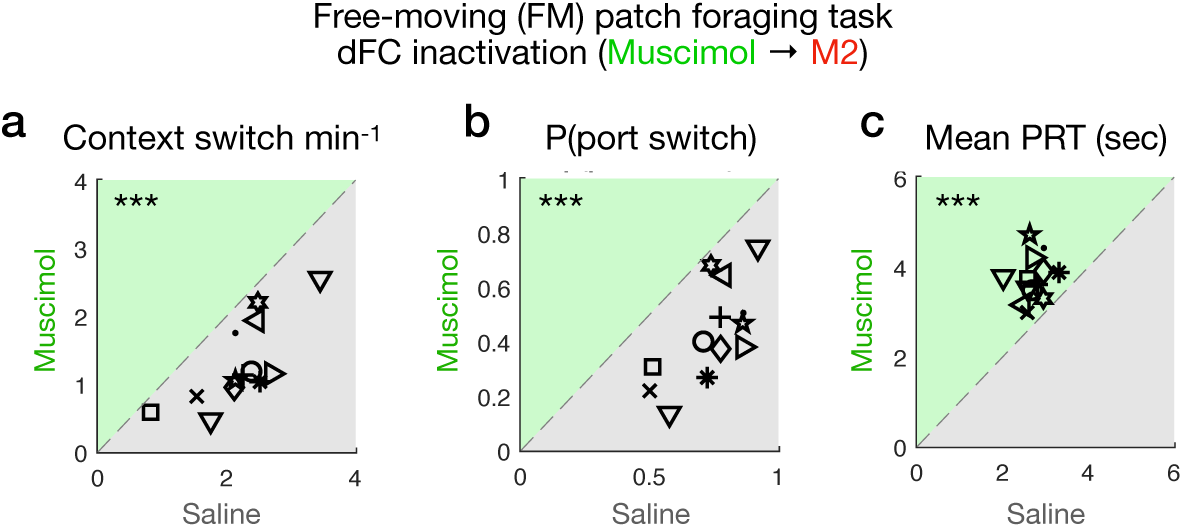
Additional analysis of behavior in dFC inactivation experiments in free-moving (FM) foraging task. a) Comparison of context transition rate (switches/min) for mice in the free-moving (FM) foraging task under Muscimol injected in M2 vs. saline (SR test, *P* = 2.44e-4, *n* = 10 mice). b) Comparison of port switch rate (average probability of port transition) for mice in the FM task under Muscimol vs. saline (SR test, *P* = 2.44e-4, *n* = 10 mice). c) Comparison of average patch residence times (PRT) for mice in the FM task under Muscimol vs. saline (SR test, *P* = 2.44e-4, *n* = 10 mice). Each dot is one mouse; Wilcoxon signed rank test (SR test); \*\*\**P* < 0.001

**Supplementary Table 1.** Brain region categories and Allen atlas correspondence.

| Region name | Tag | Macro-region category | Allen atlas labels |
| --- | --- | --- | --- |
| Secondary motor cortex | M2 | Dorsal frontal cortex (dFC) | 'Secondary motor area' |
| Anterior Cingulate cortex | ACC | Dorsal frontal cortex (dFC) | 'Anterior cingulate area' |
| Prelimbic cortex | PL | Dorsal frontal cortex (dFC) | 'Prelimbic' |
| Infralimbic cortex | IL | Ventral frontal cortex (vFC) | 'Infralimbic' |
| Orbitofrontal cortex | OFC | Ventral frontal cortex (vFC) | 'Orbital' |
| Dorsal peduncle | DP | Ventral frontal cortex (vFC) | 'Dorsal peduncular' |
| Tenia tecta | TT | Sensory cortex (SC) | 'Tenia tecta' |
| Olfactory areas | OLF | Sensory cortex (SC) | 'Olfactory areas' |
| Somatosensory areas | SS | Sensory cortex (SC) | 'somatosensory' |
| Visual areas | VIS | Sensory cortex (SC) | 'visual areas' |
| Posterior parietal cortex | PPC | Sensory cortex (SC) | 'anterior area' |
| Retrosplenial cortex | RSC | Sensory cortex (SC) | 'Retrosplenial' |
| Dentate gyrus | DG | Hippocampal (HC) | 'Dentate gyrus' |
| Hippocampus CA1 | CA1 | Hippocampal (HC) | 'Field CA1' |
| Hippocampus CA3 | CA3 | Hippocampal (HC) | 'Field CA3' |
| Striatum | STR | Basal ganglia (BG) | 'Striatum' |
| Caudoputamen | CP | Basal ganglia (BG) | 'Caudoputamen' |
| Nucleus accumbens | NAC | Basal ganglia (BG) | 'accumbens' |
| Lateral geniculate nucleus | LGN | Thalamic (THL) | 'lateral geniculate' |
| Mediodorsal thalamus | MDT | Thalamic (THL) | 'Mediodorsal nucleus of the thalamus' |
| Centromedial thalamus | CMT | Thalamic (THL) | 'Central medial nucleus of the thalamus'<br>'Rhomboid nucleus' |
| Ventral thalamus | VPT | Thalamic (THL) | 'Ventral posteromedial nucleus of the thalamus'<br>'Ventral posterolateral nucleus of the thalamus' |
| Posterior thalamus | PNT | Thalamic (THL) | 'posterior nucleus of the thalamus'<br>'Posterior complex of the thalamus'<br>'Posterior triangular thalamic nucleus'<br>'Suprageniculate nucelus'<br>'Ethmoid nucleus of the thalamus' |
| Paraventricular thalamus | PVT | Thalamic (THL) | 'Paraventricular nucleus of the thalamus' |
| Parafascicular nucleus | PFS | Thalamic (THL) | 'Subparafascicular area' |
| Lateral septum rostral | LSR | Subcortical (SUB) | 'Lateral septal nucleus rostral' |
| Lateral septum ventral | LSV | Subcortical (SUB) | 'Lateral septal nucleus ventral' |
| Medial septum | MSP | Subcortical (SUB) | 'Medial septal nucleus' |
| Precommissural nucleus | PSN | Subcortical (SUB) | 'Precommissural nucleus' |
| Hebnula | HAB | Subcortical (SUB) | 'habenula' |
| Hypothalamus | HYP | Subcortical (SUB) | 'Hypothalamus'<br>'Preoptic area' |
| Midbrain | MID | Subcortical (SUB) | 'Midbrain' |

**Supplementary Table 2.**
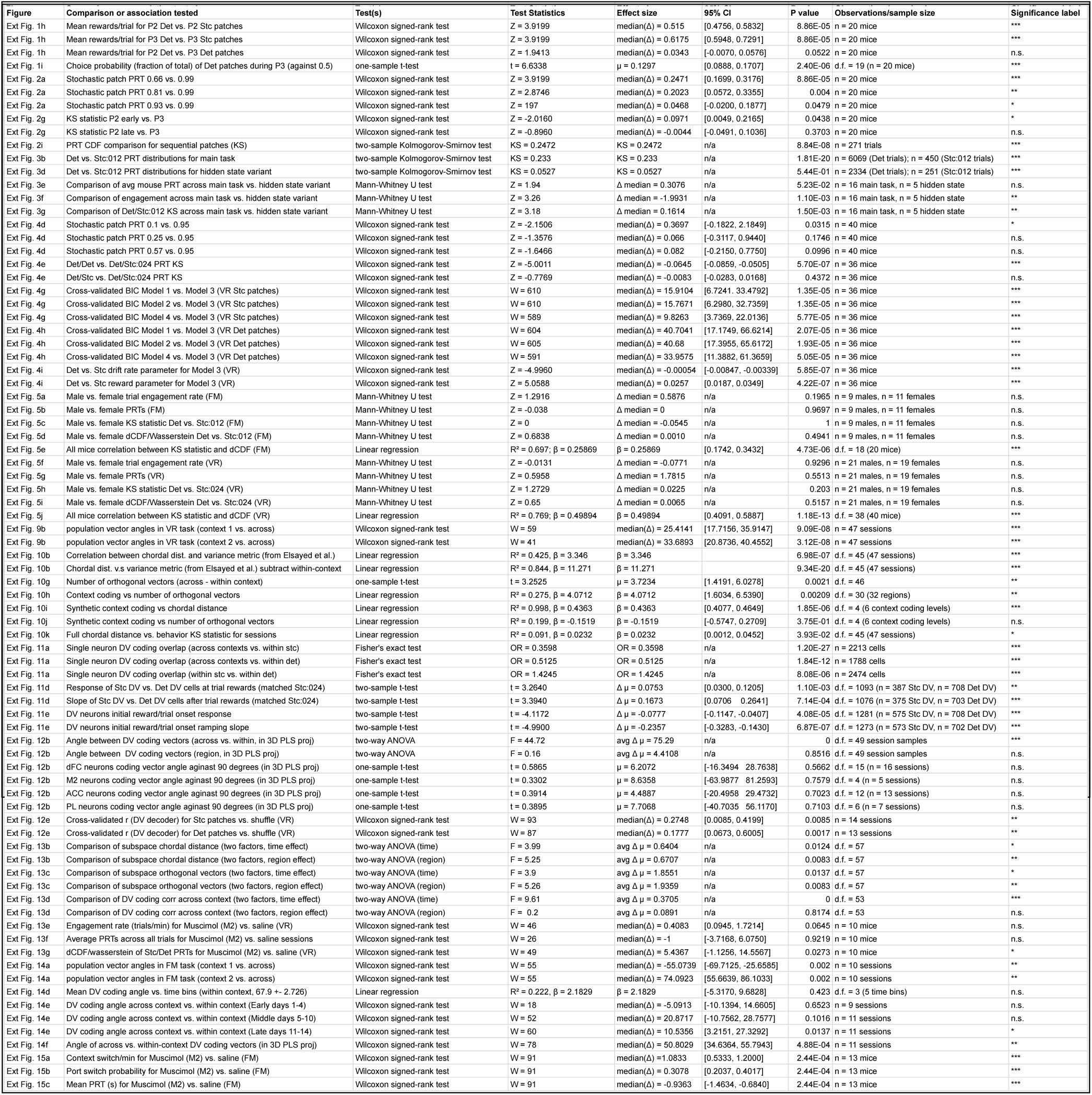
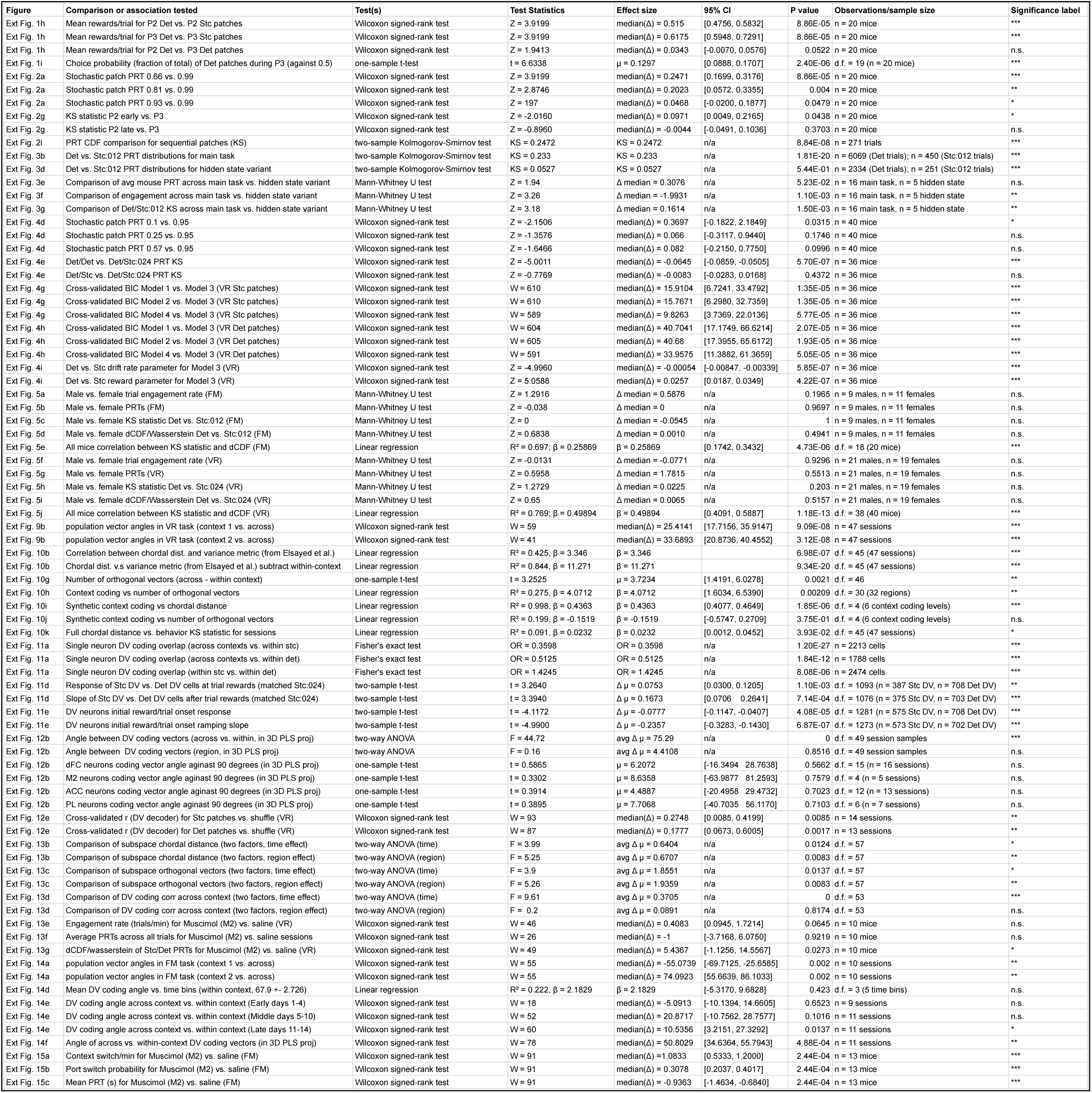
Additional details for statistical tests and quantification (Main figures).

## Notes

### Competing Interest Statement

The authors have declared no competing interest.

### Summary of Updates

Additional experiments and analysis, re-structuring of figure presentation, and substantial textual edits and changes to manuscript.

